# Predicting T Cell Quality During Manufacturing Through an Artificial Intelligence-based Integrative Multi-Omics Analytical Platform

**DOI:** 10.1101/2021.05.05.442854

**Authors:** Valerie Y. Odeh-Couvertier, Nathan J. Dwarshuis, Maxwell B. Colonna, Bruce L. Levine, Arthur S. Edison, Theresa Kotanchek, Krishnendu Roy, Wandaliz Torres-Garcia

## Abstract

Large-scale, reproducible manufacturing of therapeutic cells with consistently high quality is vital for translation to clinically effective and widely accessible cell therapies. However, the biological and logistical complexity of manufacturing a living product, including challenges associated with their inherent variability and uncertainties of process parameters, currently make it difficult to achieve predictable cell-product quality. Using a degradable microscaffold-based T cell process as an example, we developed an Artificial Intelligence (AI)-driven experimental-computational platform to identify a set of critical process parameters (CPP) and critical quality attributes (CQA) from heterogeneous, high dimensional, time-dependent multi-omics data, measurable during early stages of manufacturing and predictive of end-of-manufacturing product quality. Sequential, Design-of-Experiment (DOE)-based studies, coupled with an agnostic machine-learning framework, were used to extract feature combinations from media assessment that were highly predictive of total live CD4^+^ and CD8^+^ naïve and central memory (CD63L^+^CCR7^+^) T cells and their ratio in the end-product. This computational workflow could be broadly applied to any cell therapy and provide a roadmap for discovering CQAs and CPPs in cell manufacturing.

## Introduction

T cell-based immunotherapies have received great interest from clinicians and industry due to their potential to treat, and often functionally cure some cancers and their potential applicability in many other diseases^1,2^. Since 2017, four genetically modified autologous Chimeric Antigen Receptor (CAR) T cell therapies (*Yescarta*™, *Kymriah*™, *Tecartus*™, *Breyanzi*^®^) have received FDA approval to treat certain B-cell malignancies. Despite these successes, CAR-T cell therapies are constrained by poorly-understood manufacturing processes that are time-intensive, expensive, and difficult to scale^3,4^ with a lack of methods and tools to predict product quality during manufacturing and identify product Critical Quality Attributes (CQAs) and the associated Critical Process Parameters (CPPs).

Translating laboratory-scale T cell expansion experiments into a large-scale manufacturing process is hindered by the incomplete understanding of cell properties and how they are affected by process variables, lack of detailed characterization, and high variability of materials during manufacturing^5^. These challenges of manufacturing a “living product” are further magnified since current chemistry, manufacturing, and control (CMC), analytics, regulations, and product-specifications are designed for conventional chemical and biopharmaceutical manufacturing systems^6^. This underscores the need to develop innovative tools, methods, and standards to ensure appropriate quality controls, and new strategies involving quality by design (QbD) and good manufacturing practices (GMP) for cell-based therapies^7–9^. The intricate manufacturing process for T cells and other cell therapies must be deeply assessed and appropriately controlled to ensure scalability, predictability, and a high-quality manufacturing process at the most reasonable cost. A key step for reaching this goal is to identify putative CQAs and CPPs early in the manufacturing process that can predict the quality of the manufactured cell-therapy product. We hypothesized that rigorous characterization of process parameters along with longitudinal measurements of cell-secreted cytokine, chemokine, and metabolites from the culture media early during manufacturing will allow us to develop an AI-based mathematical-computational framework for the identification of multivariate parameters that are predictive of the end-of-manufacturing product phenotypes.

Characterization studies of approved autologous anti-CD19 CAR-T cell therapies have recently revealed initial sets of candidate quality attributes, i.e. percent transduction, vector copy number, and interferon-γ production for Axicabtagene ciloleucel (Yescarta™)^10^ while CAR expression and release of interferon-γ are a few of those identified for Tisagenlecleucel (Kymriah™)^11^. Many of these attributes are calculated as endpoint responses and thus a deeper understanding of the cell growth process impacted by starting conditions and performance during their manufacturing is essential. Hence, CQAs that enable early monitoring through real-time process measurements such as multi-omics cell characterization can overcome current challenges in assessing product consistency. Yet, the computational complexity of dealing with the heterogeneity and multivariate nature of multi-omics measurements to characterize T cell quality, i.e., high definition phenotyping of naïve and memory subsets, remains a challenge.

Generally, T cells with a lower differentiation state such as naïve and stem cell or central memory cells have been shown to provide superior anti-tumor potency, presumably due to their higher potential to replicate, migrate, and engraft, leading to a long-term, durable response^18–21^. Likewise, CD4 T cells are similarly important to anti-tumor potency due to their cytokine release properties and ability to resist exhaustion^22,23^. Our group has developed a novel degradable microscaffold (DMS)-based method using porous microcarriers functionalized with anti-CD3 and anti-CD28 mAbs for use in T cell expansion cultures. We showed that compared to commercially available microbeads (Miltenyi), degradable microscaffolds (DMSs) generated a higher number of migratory naïve (T_N_) and central-memory (T_CM_) (CCR7^+^CD62L^+^) T cells and CD4^+^ T cells across multiple donors^12^. We used this manufacturing process as an exemplar to develop an experimental-computational AI-based tool to predict product quality from early process measurements. This two-phase approach consists of (1) the optimization of process parameters through experimental designs, and (2) the extraction of early predictive signatures of T cell quality by multi-omics integration using regression models. This agnostic computational approach provides a platform to discover early predictive CQAs and CPPs to ensure consistent product quality, that can be widely applicable for other cellular therapies.

## Results

### I. Overall multi-omics study design

T cells were expanded *ex vivo* for 14 days and 100 μL of supernatant media samples were collected at days 4, 6, 8, 11, and 14 to measure cytokine profiles and perform NMR analysis. Endpoint responses on DMS-based T cell extracts were measured for different combinations of DMS parameters: IL2 concentration, DMS concentration, and functionalized antibody percent. Two experimental regions were determined using a design-of-experiments (DOE) methodology to maximize the yields of CD62L^+^CCR7^+^ cells (i.e. naïve and central memory T cells, T_N_+T_CM_) as a function of these process parameters. The first DOE resulted in a randomized 18-run I-optimal custom design where each DMS parameter was evaluated at three levels. To further optimize this DOE in terms of total live CD4^+^ T_N_+T_CM_ cells, a sequential adaptive design-of-experiment (ADOE) was designed with 12 additional samples (Fig.1b). All 30 runs from both experiments (DOE, ADOE) were molecularly characterized to model total live T_N_+T_CM_ (a) CD4^+^, (b) CD8^+^, and (c) their ratio. The extraction of early predictive CPPs and CQAs for the expansion of T_N_+T_CM_ cells during *ex vivo* culture was performed in two phases: (1) optimization of process parameters, and (2) integration of multi-omics for predictive modeling (Fig.1).

**Fig. 1.**
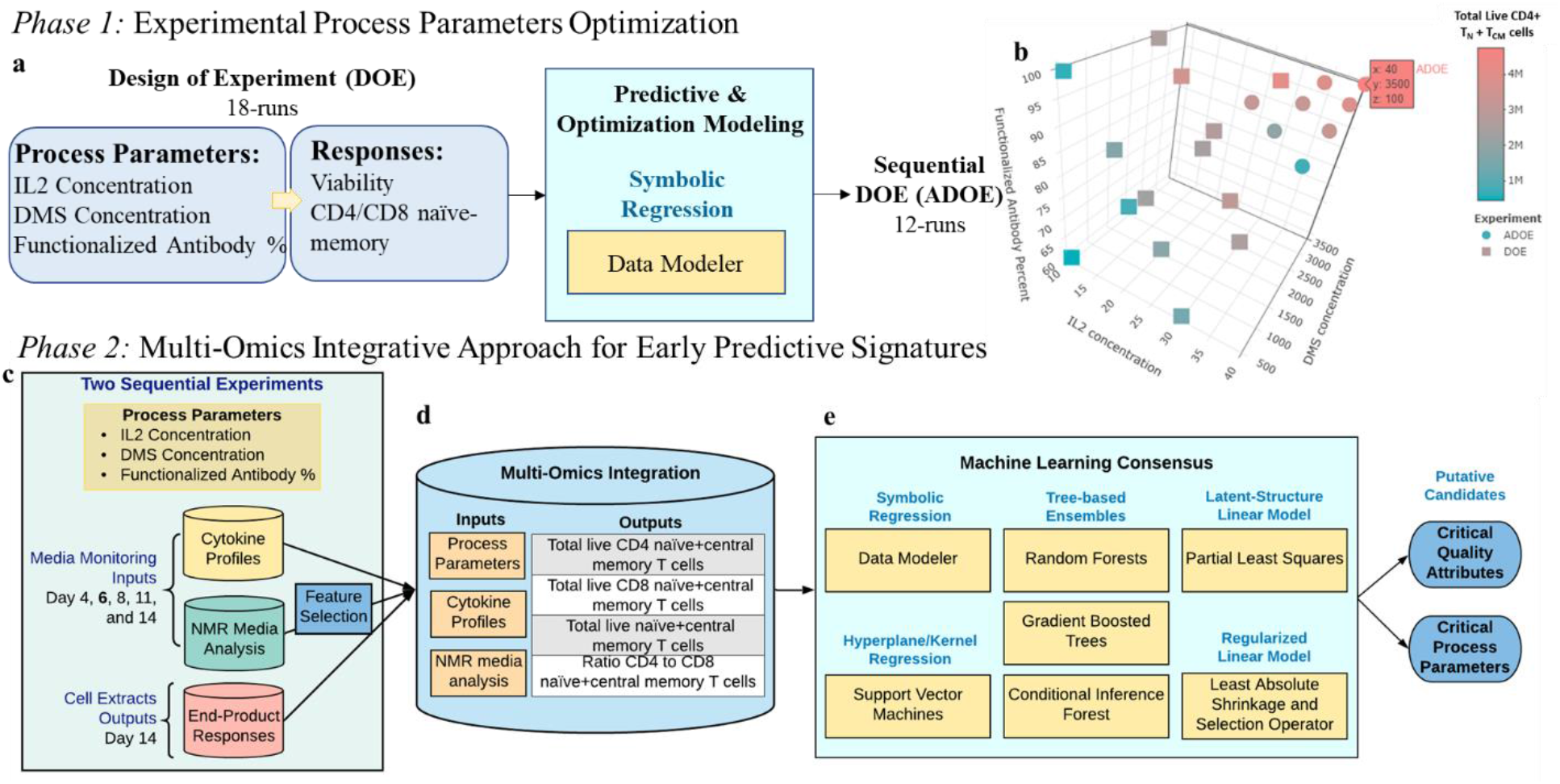
Two-phase approach to extract early predictive CPPs and CQAs for CD4^+^/CD8^+^ T_N_+T_CM_ cells. **a** DOE modeling and optimization of process parameters. **b** Experimental region studied and optimized for total live CD4^+^ T_N_+T_CM_ cells. **c** Overall study design (two experiments varying process parameters while measuring multi-omics and T_N_+T_CM_ responses. e). **d** Integrative multi-omics approach through **e** a machine learning consensus analysis to identify early predictive CPPs and CQAs putative candidates for both total live CD4^+^ and CD8^+^ T_N_+T_CM_ cells.

### II. Optimization of T_N_+T_CM_ cells as a function of process parameters

Using symbolic regression (DataModeler software from Evolved Analytics LLC), we examined the interactive effects of the DMS parameters on yield to simultaneously predict and optimize both CD4^+^ and CD8^+^ T_N_+T_CM_. A model ensemble predicted 4.2 × 10^6^ CD4^+^ T_N_+T_CM_ cells at an optimum setting of 30 U/μL IL2, 2500 carriers/μL, and 100% functionalized mAbs (Supp.Fig.S1,S3,S4). This result was consistent with the observed maximum value of 4.0 × 10^6^, highlighting that CD4^+^ T_N_+T_CM_ yield was maximized at high levels of DMS parameters (Fig.1b). In contrast, the predicted optimum yield for CD8^+^ T_N_+T_CM_ was 1.9 × 10^7^ cells at a setting of 30 U/μL IL2, 600 carriers/μL, and 100% functionalized mAbs (Supp.Fig.S2,S3,S4). Although this combination was not experimentally tested, the closest measured record (30 U/μL IL2, 500 carriers/μL, 100% functionalized mAbs) achieved the predicted maximum yield. Hence, the CD8^+^ T_N_+T_CM_ yield was maximized at high IL2 concentration and functionalized mAbs percentage but low DMS concentration.

The DOE analysis highlighted the potential for further optimization of total live CD4^+^ T_N_+T_CM_ cells, as well as the potential to optimize the CD4^+^ to CD8^+^ T_N_+T_CM_ cells ratio, at DMS levels greater than those originally evaluated (DOE). Therefore, to test and validate, a second adaptive design of experiment (ADOE) was designed to maximize the total live CD4^+^ T_N_+T_CM_ cells. We expanded the parameter range, assessing IL2 concentration>30 U/μL and DMS concentration>2500 carriers/μL (Fig.1b). CD4^+^ T_N_+T_CM_ and its ratio to CD8^+^ T_N_+ T_CM_, 4.7 × 10^6^ cell and 0.49 respectively, were maximized when IL2 concentration (40 U/μL) and DMS concentration (3500 carriers/μL) were maximized (Fig.1b;Supp.Table.S2;Supp.Fig.S1-S11). Utilizing the ADOE dataset, new response ensembles were generated enabling more robust prediction over the expanded parameter space (↑ IL2 and ↑ DMS concentrations).

### III. Multi-omic integrative analysis for early monitoring of T cell manufacturing

Due to the heterogeneity of the multivariate data collected and knowing that no single model structure is perfect for all applications, we implemented an agnostic modeling approach to better understand these T_N_+T_CM_ responses. To achieve this, a consensus analysis using seven machine learning (ML) techniques, Random Forest (RF), Gradient Boosted Machine (GBM), Conditional Inference Forest (CIF), Least Absolute Shrinkage and Selection Operator (LASSO), Partial Least-Squares Regression (PLSR), Support Vector Machine (SVM), and DataModeler’s Symbolic Regression (SR), was implemented to molecularly characterize T_N_+T_CM_ cells and to extract predictive features of quality early on their expansion process (Fig.1d-e).

SR models achieved the highest predictive performance (R^2^>93%) when using multi-omics predictors for all endpoint responses (Table.1). SR achieved R^2^>98% while GBM tree-based ensembles showed leave-one-out cross-validated R^2^ (LOO-R^2^) >95% for CD4^+^ and CD4^+^/CD8^+^ T_N_+T_CM_ responses. Similarly, LASSO, PLSR, and SVM methods showed consistent high LOO-R^2^, 92.9%, 99.7%, and 90.5%, respectively, to predict the CD4^+^/CD8^+^ T_N_+T_CM_. Yet, about 10% reduction in LOO-R^2^, 72.5%-81.7%, was observed for CD4^+^ T_N_+T_CM_ with these three methods. Lastly, SR and PLSR achieved R^2^>90% while other ML methods exhibited exceedingly variable LOO-R^2^ (0.3%,RF-51.5%,LASSO) for CD8^+^ T_N_+T_CM_ cells.

**Table 1.** LOO-R^2^ prediction performance results for all ML models when evaluating process parameters, and features from cytokine and NMR media analysis at day 6 or day 4.

| LOO-R2<br>Response/Predictors | ML |  |  |  |  |  |  |
| --- | --- | --- | --- | --- | --- | --- | --- |
|  | SR | RF | GBM | CIF | LASSO | PLSR | SVM |
| **Ratio.of.CD4.to.CD8.TN+TCM.Cells** |  |  |  |  |  |  |  |
| PP+N4 | **99%** | 86.8% | 96.3% | **84.5%** | 88.6% | 92.5% | 88.5% |
| PP+N6 | **99%** | 73.6% | 95.9% | 70.1% | 81.0% | 95.8% | 79.7% |
| PP+S6 | **99%** | **87.1%** | **99.9%** | 83.4% | 87.2% | 97.9% | 86.8% |
| PP+S6+N6 | **99%** | 85.5% | 95.3% | 83.4% | **92.9%** | **99.7%** | **90.5%** |
| **Total.live.CD4+.TN+TCM.cells** |  |  |  |  |  |  |  |
| PP+N4 | 97% | 67.0% | 93.6% | 69.3% | 34.3% | 90.1% | 75.5% |
| PP+N6 | 96% | 45.9% | 92.6% | 51.2% | 42.8% | **92.1%** | **79.4%** |
| PP+S6 | **98%** | **71.4%** | **99.9%** | **75.0%** | **74.9%** | 80.0% | 75.5% |
| PP+S6+N6 | **98%** | 68.2% | 95.6% | 74.4% | 72.5% | 81.7% | 77.0% |
| **Total.live.CD8+.TN+TCM.cells** |  |  |  |  |  |  |  |
| PP+N4 | 93% | 4.7% | **44.4%** | 9.2% | 1.2% | 65.1% | 9.1% |
| PP+N6 | 86% | 2.0% | 29.9% | **15.8%** | 28.5% | 63.3% | 30.6% |
| PP+S6 | **93%** | **7.8%** | 28.0% | 15.1% | **76.2%** | **98.4%** | **49.8%** |
| PP+S6+N6 | **93%** | 0.3% | 32.7% | 9.8% | 51.5% | 96.4% | 37.8% |

ML models prediction performance is measured as the leave-one-out cross-validated R<sup>2</sup> (LOO-R<sup>2</sup>) while SR prediction performance is measured as R<sup>2</sup> of the ensemble prediction where the ensemble is composed of diverse models with complexity constrained. Predictors evaluated: (PP) Process parameters, (N) NMR, (S) Cytokines measured at day 4 or 6. **max R<sup>2</sup> within each ML method are shown in bold.**

The top-performing technique, SR, showed that the median aggregated predictions for CD4^+^ and CD8^+^ T_N_+T_CM_ cells increases when IL2 concentration, IL15, and IL2R increase while IL17a decreases in conjunction with other features. These patterns combined with low values of DMS concentration and GM_CSF uniquely characterized maximum CD8^+^ T_N_+T_CM_. Meanwhile, higher glycine but lower IL13 in combination with others showed maximum CD4^+^ T_N_+T_CM_ predictions (Fig.2).

**Fig. 2.**
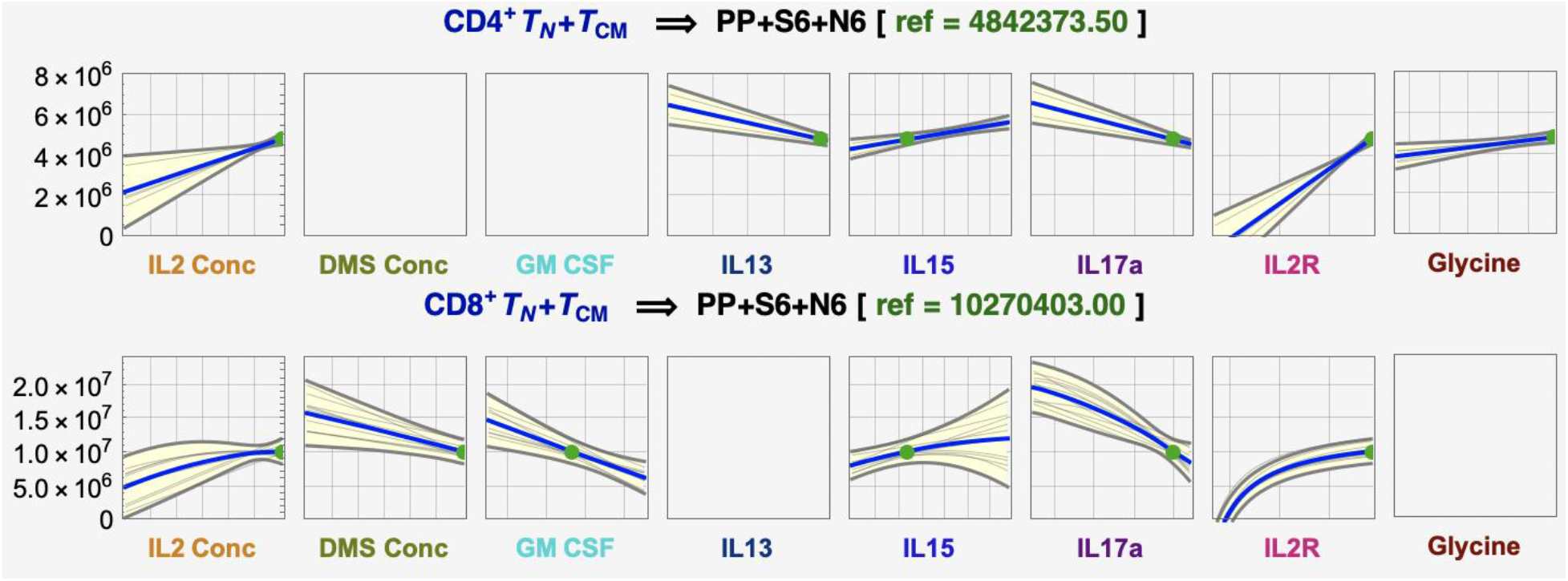
Multi-omics culturing media prediction profiles at day 6 from DataModeler. Prediction model profiles from day 6 culturing media monitoring where total live CD4^+^ T_N_+T_CM_ is maximized.

Selecting CPPs and CQAs candidates consistently for T cell memory is desired. Here, TNFα was found in consensus across all seven ML methods for predicting CD4^+^/CD8^+^ T_N_+T_CM_ when considering features with the highest importance scores across models (Fig.3a;Methods). Other features, IL2R, IL4, IL17a, and DMS concentration, were commonly selected in ≥5 ML methods (Fig.3a,c). Moreover, IL13 and IL15 were found predictive in combination with these using SR (Supp.Table.S4).

**Fig. 3.**
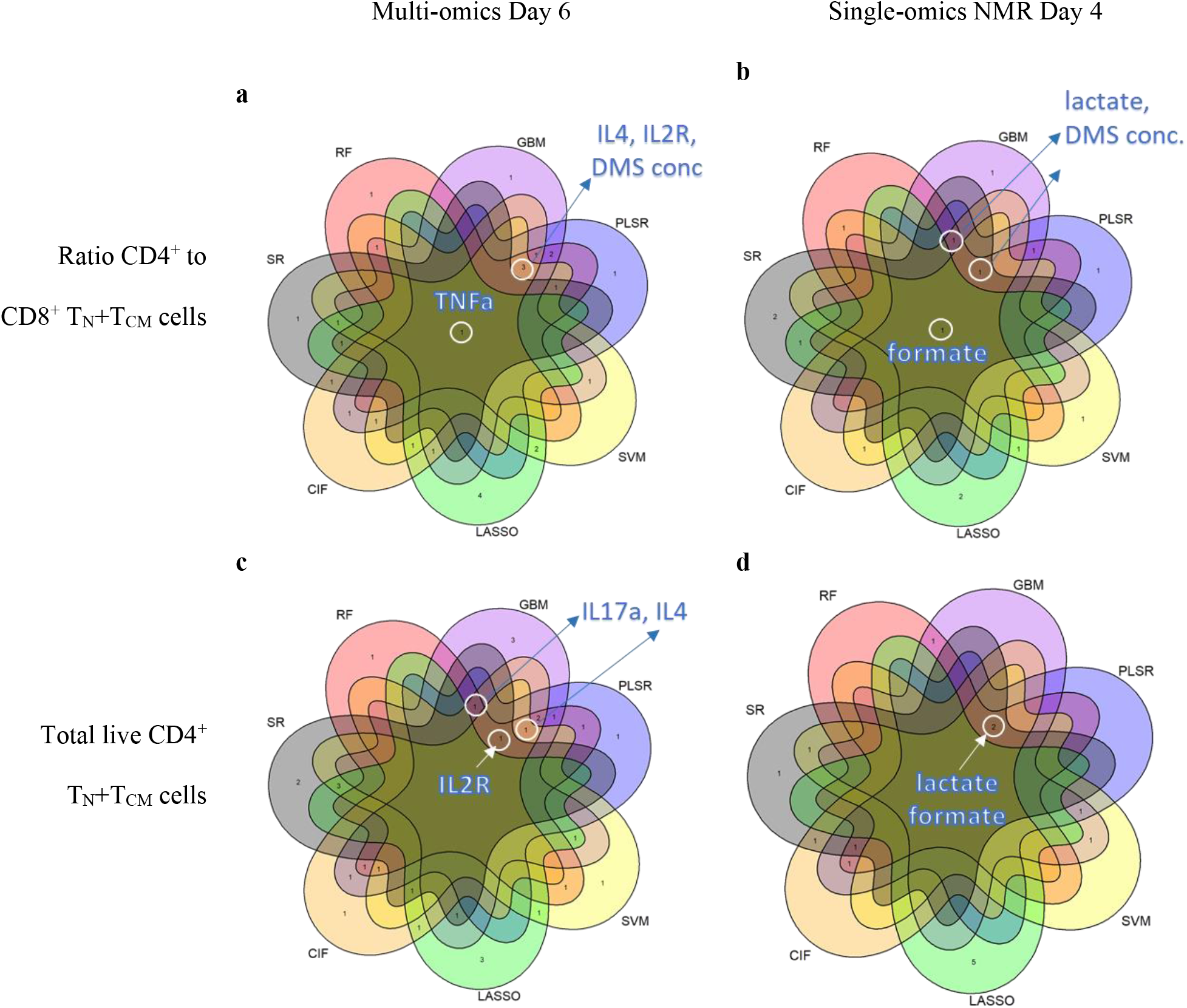
ML model consensus of highly predictive for early monitoring of T cell manufacturing. ML models consensus for **a-b** ratio CD4^+^ to CD8^+^ T_N_+T_CM_ cells, and **c-d** total live CD4^+^ T_N_+T_CM_ cells for both multi-omics modeling at day 6 and single-omics with NMR at day 4, respectively. Feature names are shown for consensus with 5 or more ML models at the highest-ranking standing (see Methods).

This integrative analysis of cytokine and NMR media analysis monitored at early stages of the T cell process provided highly predictive feature combinations of end-product quality. However, when translating a real-time monitoring strategy to a large-scale manufacturing process, measuring both cytokine and NMR features from media can be difficult and expensive. To be cost-efficient and translatable, we demonstrated that either cytokine profiles or NMR media analysis alone is sufficient to find predictive features without compromising prediction performance.

### IV. Cytokine media profiles for early prediction

ML models using solely media cytokine profiles at day 6 reached similar or higher R2 than those of the multi-omics models (CD4^+^ T_N_+T_CM_: 71.4%-99.9%; CD4^+^/CD8^+^: 83.4%-99.7%). However, CD8^+^ T_N_+T_CM_ still had variable LOO-R^2^, 7.8%-93%. Overall, higher cytokine media profiles showed higher CD4^+^ T_N_+T_CM_ and consequently its ratio with CD8^+^ (Fig.4a). This behavior was evident, even beyond day 6, for TNFα, IL2R, IL17a, and IL4 which were frequently selected as predictive features across models (Fig.4b-c;Supp.Fig.S20). A more complex behavior was detected for CD8^+^ T_N_+T_CM_ which cannot be explained by cytokine secretion alone (Fig.4d).

**Fig. 4.**
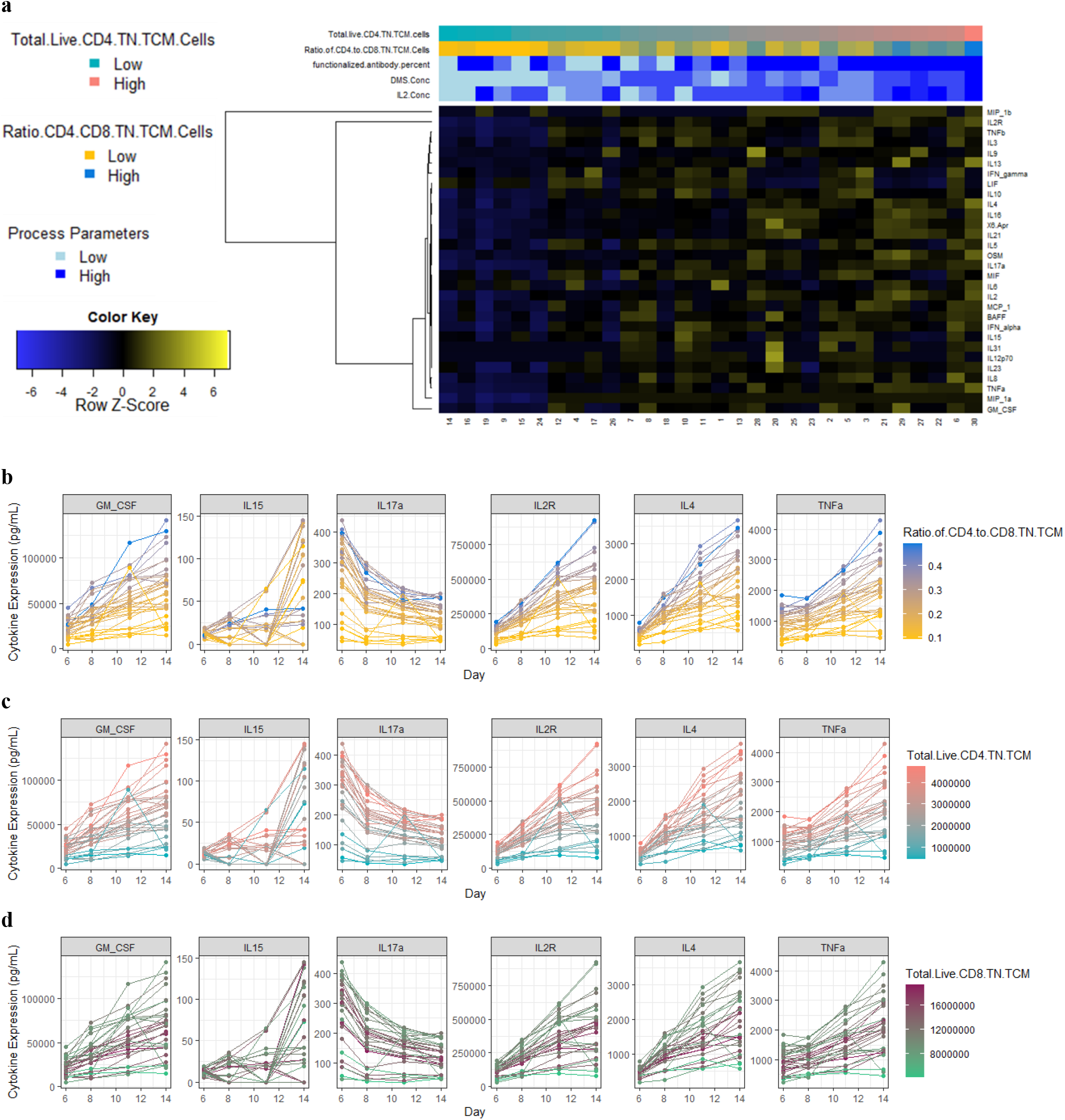
General characteristics of cytokine media profiles. **a** Heatmap for cytokine profiles from media samples on day 6. Expression in picograms/milliliter across time points for relevant cytokine features for **b** ratio CD4+ to CD8+ T_N_+T_CM_ cells, **c** total live CD4^+^ T_N_+T_CM_ cells, and **d** total live CD8^+^ T_N_+T_CM_ cells.

### V. NMR media analysis for early prediction

Models using only NMR media intensities on day 6 revealed an R^2^ decrease of 8.8% and 11.1%, on average, compared with the multi-omics and cytokine models, respectively. Yet, SR, GBM, and PLSR reached high LOO-R^2^ (92.1%-99%), specifically for CD4^+^/CD8^+^ and CD4^+^ TN+T_CM_. Although good prediction was achieved with NMR media analysis on day 6, we obtain slightly better predictions with NMR media analysis on day 4 (Table.1). From these models, formate, lactate, DMS concentration were highly ranked to predict both, ratio CD4^+^/CD8^+^ and CD4^+^ T_N_+T_CM_ (Fig.3b,d;Supp.Fig.19d). Some variable combinations also contained histidine, ethanol, dimethylamine, branch chain amino acids (BCAAs), glucose, and glutamine (Supp.Table.S3). Lower intensity values for BCAAs, dimethylamine, glucose, and glutamine displayed higher CD4^+^ T_N_+T_CM_ cells across the different media monitoring times (Supp.Fig.S25). Inversely, higher intensities of formate and lactate showed higher CD4^+^ T_N_+T_CM_ and its ratio with CD8^+^ consistently across time (Fig.5a,b).

**Fig. 5.**
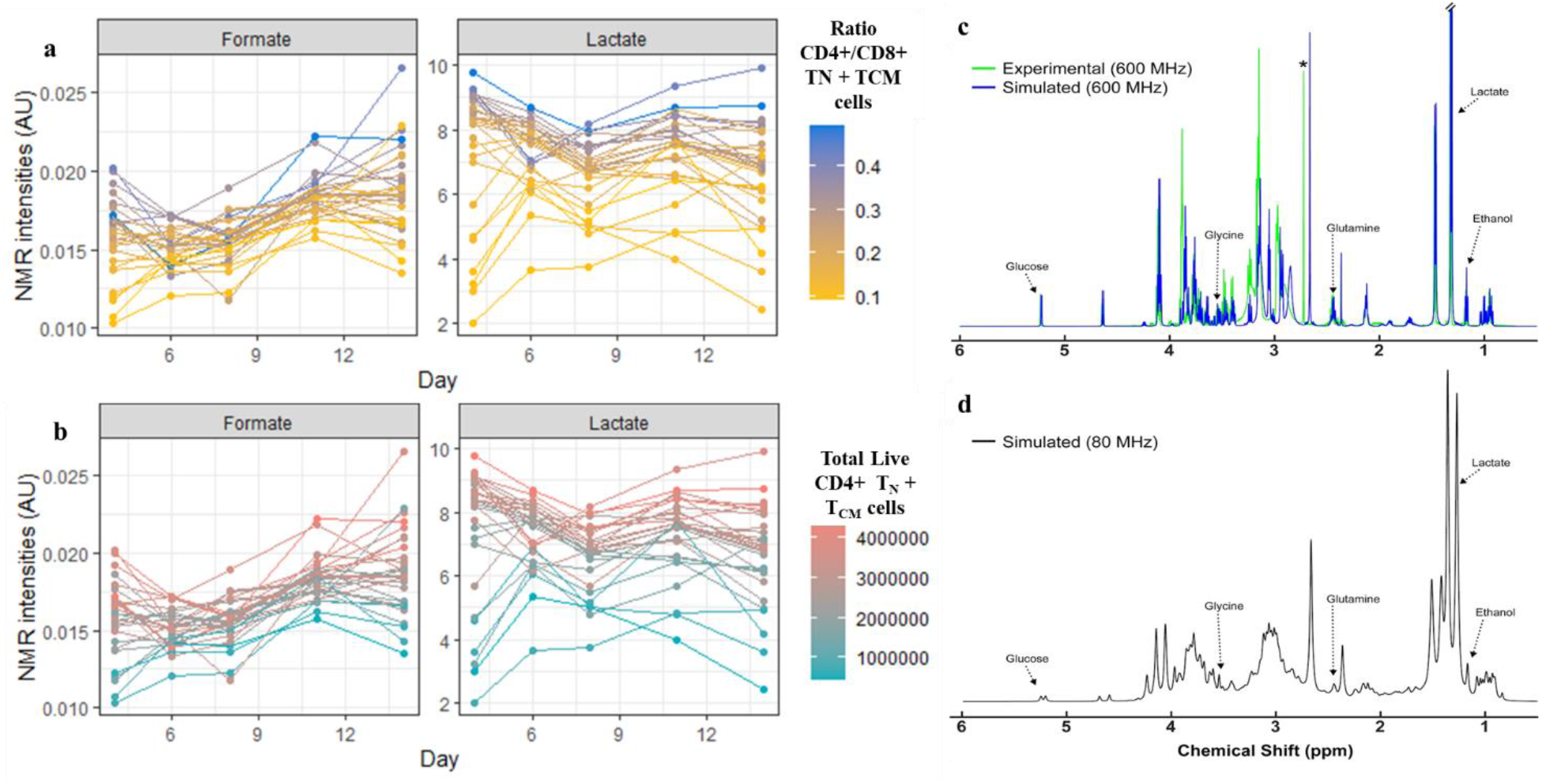
Top-performing features NMR media analysis. NMR intensities in arbitrary units (AU) across time points for **a** Ratio CD4^+^/CD8^+^ T_N_+T_CM_ cells, and **b** total live CD4^+^ T_N_+T_CM_ cells. **c** Simulation of ^1^H NMR spectrum shows the potential to detect multiple predictive features at lower magnetic fields. Overlay of a pooled experimental spectrum of T-cell culture medium (green) and GISSMO^27,28^ simulated spectrum (blue), composed of 19 compounds that reasonably approximate the experimental spectrum acquired at 600 MHz. *indicates an unknown feature of high intensity that was simulated with 2,3-dimethylamine (blue feature to right). Annotated features in the spectrum correspond to those identified as being highly predictive of output responses across computational methods. **d** GISSMO^27,28^ simulated spectrum at 80 MHz, corresponding to a field strength of commercially available benchtop NMR systems.

## Discussion

### I. Optimization of process parameters

CPPs modeling and understanding are critical to new product development and in cell therapy development, it can have life-saving implications. The challenges for effective modeling grow with the increasing complexity of processes due to high dimensionality, and the potential for process interactions and nonlinear relationships. Another critical challenge is the limited amount of available data, mostly small DOE datasets. SR has the necessary capabilities to resolve the issues of process effects modeling and has been applied across multiple industries^12^. SR discovers mathematical expressions that fit a given sample and differs from conventional regression techniques in that a model structure is not defined *a priori*^13^. Hence, a key advantage of this methodology is that transparent, human-interpretable models can be generated from small and large datasets with no prior assumptions^14,15^.

Since the model search process lets the data determine the model, diverse and competitive (e.g., accuracy, complexity) model structures are typically discovered. An ensemble of diverse models can be formed where its constituent models will tend to agree when constrained by observed data yet diverge in new regions. Collecting data in these regions helps to ensure that the target system is accurately modeled, and its optimum is accurately located^14,15^. Exploiting these features allows adaptive data collection and interactive modeling. Consequently, this adaptive-DOE approach is useful in a variety of scenarios, including maximizing model validity for model-based decision making, optimizing processing parameters to maximize target yields, and developing emulators for online optimization and human understanding^14,15^.

### II. Early predictive features

An in-depth characterization of potential DMS-based T-cell CQAs includes a list of cytokine and NMR features from media samples that are crucial in many aspects of T cell fate decisions and effector functions of immune cells. Cytokine features were observed to slightly improve prediction and dominated the ranking of important features and variable combinations when modeling together with NMR media analysis and process parameters (Fig.3b,d).

Predictive cytokine features such as TNFα, IL2R, IL4, IL17a, IL13, and IL15 were biologically assessed in terms of their known functions and activities associated with T cells. T helper cells secrete more cytokines than T cytotoxic cells, as per their main functions, and activated T cells secrete more cytokines than resting T cells. It is possible that some cytokines simply reflect the CD4^+^/CD8^+^ ratio and the activation degree by proxy proliferation. However, the exact ratio of expected cytokine abundance is less clear and depends on the subtypes present, and thus examination of each relevant cytokine is needed.

IL2R is secreted by activated T cells and binds to IL2, acting as a sink to dampen its effect on T cells^16^. Since IL2R was much greater than IL2 in solution, this might reduce the overall effect of IL2, which could be further investigated by blocking IL2R with an antibody. In T cells, TNF can increase IL2R, proliferation, and cytokine production^18^. It may also induce apoptosis depending on concentration and alter the CD4^+^ to CD8^+^ ratio^17^. Given that TNF has both a soluble and membrane-bound form, this may either increase or decrease CD4^+^ ratio and/or memory T cells depending on the ratio of the membrane to soluble TNF^18^. Since only soluble TNF was measured, membrane TNF is needed to understand its impact on both CD4^+^ ratio and memory T cells. Furthermore, IL13 is known to be critical for Th2 response and therefore could be secreted if there are significant Th2 T cells already present in the starting population^19^. This cytokine has limited signaling in T cells and is thought to be more of an effector than a differentiation cytokine^20^. It might be emerging as relevant due to an initially large number of Th2 cells or because Th2 cells were preferentially expanded; indeed, IL4, also found important, is the conical cytokine that induces Th2 cell differentiation (Fig.3). The role of these cytokines could be investigated by quantifying the Th1/2/17 subsets both in the starting population and longitudinally. Similar to IL13, IL17 is an effector cytokine produced by Th17 cells^21^ thus may reflect the number of Th17 subset of T cells. GM-CSF has been linked with activated T cells, specifically Th17 cells, but it is not clear if this cytokine is inducing differential expansion of CD8^+^ T cells or if it is simply a covariate with another cytokine inducing this expansion^22^. Finally, IL15 has been shown to be essential for memory signaling and effective in skewing CAR-T cells toward the Tscm phenotype when using membrane-bound IL15Ra and IL15R^23^. Its high predictive behavior goes with its ability to induce large numbers of memory T cells by functioning in an autocrine/paracrine manner and could be explored by blocking either the cytokine or its receptor.

Moreover, many predictive metabolites found here are consistent with metabolic activity associated with T cell activation and differentiation, yet it is not clear how the various combinations of metabolites relate with each other in a heterogeneous cell population. Formate and lactate were found to be highly predictive and observed to positively correlate with higher values of total live CD4^+^ T_N_+T_CM_ cells (Fig.5a-b;Supp.Fig.28-S30,S38). Formate is a byproduct of the one-carbon cycle implicated in promoting T cell activation^24^. Importantly, this cycle occurs between the cytosol and mitochondria of cells and formate excreted^25^. Mitochondrial biogenesis and function are shown necessary for memory cell persistence^26,27^. Therefore, increased formate in media could be an indicator of one-carbon metabolism and mitochondrial activity in the culture.

In addition to formate, lactate was found as a putative CQA of T_N_+T_CM_. Lactate is the end-product of aerobic glycolysis, characteristic of highly proliferating cells and activated T cells^28,29^. Glucose import and glycolytic genes are immediately upregulated in response to T cell stimulation, and thus generation of lactate. At earlier time-points, this abundance suggests a more robust induction of glycolysis and higher overall T cell proliferation. Interestingly, our models indicate that higher lactate predicts higher CD4^+^, both in total and in proportion to CD8^+^, seemingly contrary to previous studies showing that CD8^+^ T cells rely more on glycolysis for proliferation following activation^30^. It may be that glycolytic cells dominate in the culture at the early time points used for prediction, and higher lactate reflects more cells.

Ethanol patterns are difficult to interpret since its production in mammalian cells is still poorly understood^31^. Fresh media analysis indicates ethanol presence in the media used, possibly utilized as a carrier solvent for certain formula components. However, this does not explain the high variability and trend of ethanol abundance across time (Supp.Fig.S25-S27). As a volatile chemical, variation could be introduced by sample handling throughout the analysis process. Nonetheless, it is also possible that ethanol excreted into media over time, impacting processes regulating redox and reactive oxygen species which have previously been shown to be crucial in T cell signaling and differentiation^32^.

Metabolites that consistently decreased over time are consistent with the primary carbon source (glucose) and essential amino acids (BCAA, histidine) that must be continually consumed by proliferating cells. Moreover, the inclusion of glutamine in our predictive models also suggests the importance of other carbon sources for certain T cell subpopulations. Glutamine can be used for oxidative energy metabolism in T cells without the need for glycolysis^30^. Overall, these results are consistent with existing literature that show different T cell subtypes require different relative levels of glycolytic and oxidative energy metabolism to sustain the biosynthetic and signaling needs of their respective phenotypes^33,34^. It is worth noting that the trends of metabolite abundance here are potentially confounded by the partial replacement of media that occurred periodically during expansion (Methods), thus likely diluting some metabolic byproducts (i.e. formate, lactate) and elevating depleted precursors (i.e. glucose, amino acids). More definitive conclusions of metabolic activity across the expanding cell population can be addressed by a closed system, ideally with on-line process sensors and controls for formate, lactate, along with ethanol and glucose.

### III. Monitoring of T-cell manufacturing with benchtop NMR systems

We demonstrated the ability to identify predictive markers using high-magnetic field NMR spectrometers. However, these are expensive, require a significant amount of resources to house and maintain, and would be the unlikely option for routine monitoring in industrial cell-manufacturing. Another common method, liquid chromatography (LC) coupled to mass spectrometry, has the advantage of a relatively smaller footprint and less upfront cost but it has other drawbacks such as destruction of the sample and difficulty with components in culture media that damage LC columns without extraction. Nevertheless, methods like continuous closed-loop sampling are being developed to address this and might be readily available in the future^35^. Recently, permanent magnet-based NMR spectrometers (benchtop-size) have become available at a lower cost. Many of these are readily configured for flow-through reaction monitoring, which can be leveraged in a closed-cell manufacturing process. To explore the feasibility of such system, we utilized a spectral simulation to evaluate if putative CQAs identified here could theoretically be observed and quantified at a magnetic field strength of 80 MHz (benchtop systems). First, the experimental data acquired at 600 MHz was approximated by creating a simulated mixture of identified metabolites (Fig.5c) and then simulated at 80 MHz (Fig.5d). While the spectral resolution is significantly reduced compared to a spectrum at high-field, there are still numerous features that can be attributed to unique metabolites, including those identified as highly predictive (Fig.5c,d). Although this is promising, there will be challenges to acquiring high-quality data in a closed bioreactor system, i.e. cells/DMS-particles in suspension, media formulation dictated by spectral complexity/overlap, and accurate quantitation of features with high overlap from other signals. However, a dedicated benchtop NMR coupled to a bioreactor could provide a simple system for real-time monitoring of CQAs.

Henceforth, this two-phase approach enabled in-depth characterization and identification of potential CQAs and CPPs for T cells. More sampling is needed to explore aspects like donor-to-donor variability, when available it can be incorporated into this workflow which will be enriched due to its data-driven iterative design that fine-tunes model parameters as more data fits back into it. Providing a powerful framework to optimize a complex experimental space during the cell-manufacturing process, and to facilitate the identification of CPPs and early predictive CQAs from multi-omics, that can be used broadly in the cell therapy and regenerative medicine field to accurately predict end-of-manufacturing quality at early stages.

## Methods

### I. Overall multi-omics study design and development: More details

The first DOE resulted in a randomized 18-run I-optimal custom design where each DMS parameter was evaluated at three levels: IL2 concentration (10, 20, and 30 U/μL), DMS concentration (500, 1500, 2500 carrier/μL), and functionalized antibody percent (60%, 80%, 100%). These 18 runs consisted of 14 unique parameter combinations where 4 of them were replicated twice to assess prediction error. Process parameters for the ADOE were evaluated at multiple levels: IL2 concentration (30, 35, and 40 U/μL), DMS concentration (500, 1000, 1500, 2000, 2500, 3000, 3500 carrier/μL), and functionalized antibody percent (100%) as depicted in Fig.1b. To further optimize the initial region explored (DOE) in terms of total live CD4^+^ T_N_+T_CM_ cells, a sequential adaptive design-of-experiment (ADOE) was designed with 10 unique parameter combinations, two of these replicated twice for a total of 12 additional samples (Fig.1b). The fusion of cytokine and NMR profiles from media to model these responses included 30 cytokines from a custom Thermo Fisher ProcartaPlex Luminex kit and 20 NMR features. These 20 spectral features from NMR media analysis were selected out of approximately 250 peaks through the implementation of a variance-based feature selection approach and some manual inspection steps.

### II. Microcarrier fabrication

Degradable microscaffolds were fabricated as previously described^36^. Briefly, gelatin microcarriers (CuS, GE Healthcare DG-2001-OO) were suspended at 20 mg*/*mL in 1X phosphate-buffered saline (PBS). Sulfo-NHS-biotin (SNB) (Thermo Fisher 21217 or Apex Bio A8001) was dissolved at 10 µM in ultrapure water and 7.5 µL SNB*/*mL PBS was added to carrier suspension and allowed to react for 60 min. After washing the carriers three times in PBS, 40 µg*/*mL streptavidin (Jackson Immunoresearch 016-000-114) was added and allowed to react for 60 min. Biotinylated mAbs against human CD3 and CD28 were combined in a 1:1 mass ratio and added to the carriers at 2 µg mAbs*/*mg carriers. To vary the surface concentration of the antibodies, the anti-CD3/anti-CD28 mAb mixture was further combined with a biotinylated isotype control to reduce the overall fraction of targeted mAbs. mAbs were allowed to bind to the carriers for 60 min. All mAbs were low endotoxin azide-free (Biolegend custom, LEAF specification). Fully functionalized DMSs were washed in sterile PBS and washed once again in the cell culture media to be used for the T cell expansion. The surface concentration of the antibodies was quantified as previously described using a bicinchoninic acid assay (BCA) kit (Thermo Fisher 23227)^36^.

### III. T cell culture (including sample collection)

Cryopreserved primary human T cells were obtained as sorted CD3 subpopulations (Astarte Biotech). T cells were activated by adding DMSs (amount specified by the DOE) at day 0 of culture immediately after thaw. DMSs were not added or removed during the culture and had antibodies that were conjugated in proportions specified by the DOE. Initial cell density was 2.0*10^6^ cells*/*mL in a 96 well plate with 300 µL volume. Media was serum-free TexMACS (Miltentyi Biotech 170-076-307) supplemented with recombinant human IL2 in concentrations specified by the DOE (Peprotech 200-02). Cell cultures were expanded for 14 days as counted from the time of initial seeding and activation. Cell counts and viability were assessed using acridine orange/propidium iodide (AO/PI) and a Countess Automated Cell Counter (Thermo Fisher). Media was added to cultures every 2 days to 3 days in a 3:1 ratio (new volume: old volume) or based on a 300 mg/dL glucose threshold. The ADOE was done using the same feeding schedule as the initial DOE to maintain consistency for validation. Media glucose was measured using a ChemGlass glucometer to confirm cell growth and activation.

### IV. Flow cytometry

At the end of culture, at least 1e5 T cells from each run were washed with PBS once, resuspended in PBS, and stained with Zombie UV (Biolegend, 423107) for 30 minutes at room temperature in the dark at a 1:1000 dilution. Cells were spun and resuspended in FACS buffer (1X PBS, 2% bovine serum albumin, 5 mM EDTA) and were stained with antibodies according to **Table M1** for 60 minutes in the dark at 4C. Cells were then resuspended in fresh FACS buffer, after which they were run on a BD LSR ortessa. All stained was performed in a 96 well v-bottom plate.

**Table M1:** Flow cytometry antibodies.

| Antigen | Fluorophore | Vendor | Cat Number |
| --- | --- | --- | --- |
| CD3 | APC-Fire | Biolegend | 34839 |
| CD4 | PerCP-Cy5.5 | BD | 561438 |
| CCR7 | AF647 | BD | 561438 |
| CD62L | PE | BD | 341012 |

### V. Cytokine measurements

Cytokines were measured using a custom ProcartaPlex Luminex kit (Thermo Fisher). The assay was performed using media samples taken at various time points throughout the T cell culture according to the manufacturer’s instructions with modifications to half the reagent requirements. Briefly, an 8 point standard curve was created with all included standards. 25 μL magnetic beads were added to all required wells and washed three times. 25 μL of each standard or sample was added to the wells and the plate was sealed and spun at 850 rpm for 120 minutes followed by three washes. 12.5 μL detection antibody was added followed by sealing the plate and spinning for 60 minutes at 850 rpm and three washes. 25 μL streptavidin PE was added followed by the same spin and wash steps. 120 μL of reading buffer was added to the plate, the plate was analyzed on a BioPlex 200 (BioRad). Any samples that were majority over-range (denoted as “OOR >” in the output spreadsheet) were deemed too concentrated at run at 1/10th their original concentration to put them within range. All samples were run without technical replicates.

Luminex data was preprocessed using R for inclusion in the analysis pipeline as follows. Any cytokine level that was over-range (“OOR >” in output) was set to the maximum value of the standard curve for that cytokine. Any value that was under-range (“OOR <” in output spreadsheet) was set to zero. All values that were extrapolated from the standard curve were left unchanged.

### VI. NMR metabolomics

#### A. Sample preparation

50 μL of media was collected from each culture at each time point (before media exchange, if applicable), flash-frozen in liquid nitrogen, and stored at -80°C. Samples were shipped to CCRC on dry ice for NMR analysis. Run order of samples was randomized. Samples were prepared in two batches for each rack of NMR samples to be run. For each rack, samples were pulled and sorted on dry ice, then thawed at 4°C for 1 hour. Samples were then centrifuged at 2,990 x g at 4°C for 20 minutes to pellet any cells or debris that may have been collected with the media. 5 μL of 100/3 mM DSS-D6 in deuterium oxide (Cambridge Isotope Laboratories) were added to 1.7 mm NMR tubes (Bruker BioSpin), followed by 45 μL of media from each sample that was added and mixed, for a final volume of 50 μL in each tube. Samples were prepared on ice and in predetermined, randomized order. The remaining volume from each sample in the rack (∼4 μL) was combined to create an internal pool. This material was used for internal controls within each rack as well as metabolite annotation.

#### B. Data collection

NMR spectra were collected on a Bruker Avance III HD spectrometer at 600 MHz using a 5-mm TXI cryogenic probe and TopSpin software (Bruker BioSpin). One-dimensional spectra were collected on all samples using the noesypr1d pulse sequence under automation using ICON NMR software. Two-dimensional HSQC and TOCSY spectra were collected on internal pooled control samples for metabolite annotation.

#### C. Data processing

One-dimensional spectra were manually phased and baseline corrected in TopSpin. Two-dimensional spectra were processed in NMRpipe^37^. One dimensional spectra were referenced, water/end regions removed, and normalized with the PQN algorithm^38^ using an in-house MATLAB (The MathWorks, Inc.) toolbox (https://github.com/artedison/Edison_Lab_Shared_Metabolomics_UGA).

#### D. Feature selection

To reduce the total number of spectral features from approximately 250 peaks and enrich for those that would be most useful for statistical modeling, a variance-based feature selection was performed within MATLAB. For each digitized point on the spectrum, the variance was calculated across all experimental samples and plotted. Clearly-resolved features corresponding to peaks in the variance spectrum were manually binned and integrated to obtain quantitative feature intensities across all samples (Supp.Fig.S24). In addition to highly variable features, several other clearly resolved and easily identifiable features were selected (glucose, BCAA region, etc). Some features were later discovered to belong to the same metabolite but were included in further analysis.

#### E. Metabolite annotation

Two-dimensional spectra collected on pooled samples were uploaded to COLMARm web server^10^, where HSQC peaks were automatically matched to database peaks. HSQC matches were manually reviewed with additional 2D and proton spectra to confirm the match. Annotations were assigned a confidence score based upon the levels of spectral data supporting the match as previously described^11^. Annotated metabolites were matched to previously selected features used for statistical analysis.

#### F. Low-field spectrum simulation

Using the list of annotated metabolites obtained above, an approximation of a representative experimental spectrum was generated using the GISSMO mixture simulation tool.^39,40^ With the simulated mixture of compounds, generated at 600 MHz to match the experimental data, a new simulation was generated at 80 MHz to match the field strength of commercially available benchtop NMR spectrometers. The GISSMO tool allows visualization of signals contributed from each individual compound as well as the mixture, which allows annotation of features in the mixture belonging to specific compounds.

#### G. Unknown identification

Several low abundance features selected for analysis did not have database matches and were not annotated. Statistical total correlation spectroscopy^41^ suggested that some of these unknown features belonged to the same molecules (not shown). Additional multidimensional NMR experiments will be required to determine their identity.

### VII. Machine learning techniques & statistical analysis

#### A. Machine learning modeling

Seven machine learning (ML) techniques were implemented to predict three responses related to the memory phenotype of the cultured T cells under different process parameters conditions (i.e. Total Live CD4+ T_N_ and T_CM_, Total Live CD8+ T_N_+T_CM,_ and Ratio CD4+/CD8+ T_N_+T_CM_). The ML methods executed were Random Forest (RF), Gradient Boosted Machine (GBM), Conditional Inference Forest (CIF), Least Absolute Shrinkage and Selection Operator (LASSO), Partial Least-Squares Regression (PLSR), Support Vector Machine (SVM), and DataModeler’s Symbolic Regression (SR). Primarily, SR models were used to optimize process parameter values based on T_N_+T_CM_ phenotype and to extract early predictive variable combinations from the multi-omics experiments. Furthermore, all regression methods were executed, and the high-performing models were used to perform a consensus analysis of the important variables to extract potential critical quality attributes and critical process parameters predictive of T-cell potency, safety, and consistency at the early stages of the manufacturing process.

Symbolic regression (SR) was done using Evolved Analytics’ DataModeler software (Evolved Analytics LLC, Midland, MI). DataModeler utilizes genetic programming to evolve symbolic regression models (both linear and non-linear) rewarding simplicity and accuracy. Using the selection criteria of highest accuracy (R^2^>90% or noise-power) and lowest complexity, the top-performing models were identified. Driving variables, variable combinations, and model dimensionality tables were generated. The top-performing variable combinations were used to generate model ensembles. In this analysis, DataModeler’s *SymbolicRegression* function was used to develop explicit algebraic (linear and nonlinear) models. The fittest models were analyzed to identify the dominant variables using the *VariablePresence* function, the dominant variable combinations using the *VariableCombinations* function, and the model dimensionality (number of unique variables) using the *ModelDimensionality* function. *CreateModelEnsemble* was used to define trustable model ensembles using selected variable combinations and these were summarized (model expressions, model phenotype, model tree plot, ensemble quality, model quality, variable presence map, ANOVA tables, model prediction plot, exportable model forms) using the *ModelSummaryTable* function. Ensemble prediction and residual performance were respectively assessed via the *EnsemblePredictionPlot* and *EnsembleResidualPlot* subroutines. Model maxima (*ModelMaximum* function) and model minima (*ModelMinimum* function) were calculated and displayed using the *ResponsePlotExplorer* function. Trade-off performance of multiple responses was explored using the *MultiTargetResponseExplorer* and *ResponseComparisonExplorer* with additional insights derived from the *ResponseContourPlotExplorer*. Graphics and tables were generated by DataModeler. These model ensembles were used to identify predicted response values, potential optima in the responses, and regions of parameter values where the predictions diverge the most.

Non-parametric tree-based ensembles were done through the *randomForest, gbm*, and *cforest* regression functions in R, for random forest, gradient boosted trees, and conditional inference forest models, respectively. Both random forest and conditional inference forest construct multiple decision trees in parallel, by randomly choosing a subset of features at each decision tree split, in the training stage. Random forest individual decision trees are split using the Gini Index, while conditional inference forest uses a statistical significance test procedure to select the variables at each split, reducing correlation bias. In contrast, gradient boosted trees construct regression trees in series through an iterative procedure that adapts over the training set. This model learns from the mistakes of previous regression trees in an iterative fashion to correct errors from its precursors’ trees (i.e. minimize mean squared errors). Prediction performance was evaluated using leave-one-out cross-validation (LOO)-R^2^ and permutation-based variable importance scores assessing % increase of mean squared errors (MSE), relative influence based on the increase of prediction error, coefficient values for RF, GBM, and CID, respectively. Partial least squares regression was executed using the *plsr* function from the *pls* package in R while LASSO regression was performed using the *cv*.*glmnet* R package, both using leave-one-out cross-validation. Finally, the *kernlab* R package was used to construct the Support Vector Machine regression models.

Parameter tuning was done for all models in a grid search manner using the *train* function from the *caret* R package using LOO-R^2^ as the optimization criteria. Specifically, the number of features randomly sampled as candidates at each split (mtry) and the number of trees to grow (ntree) were tuned parameters for random forest and conditional inference forest. In particular, minimum sum of weights in a node to be considered for splitting and the minimum sum of weights in a terminal node were manually tuned for building the CIF models. Moreover, GBM parameters such as the number of trees to grow, maximum depth of each tree, learning rate, and the minimal number of observations at the terminal node, were tuned for optimum LOO-R^2^ performance as well. For PLSR, the optimal number of components to be used in the model was assessed based on the standard error of the cross-validation residuals using the function *selectNcomp* from the *pls* package. Moreover, LASSO regression was performed using the *cv*.*glmnet* package with *alpha* = 1. The best lambda for each response was chosen using the minimum error criteria. Lastly, a fixed linear kernel (i.e. svmLinear) was used to build the SVM regression models evaluating the cost parameter value with best LOO-R^2^. Prediction performance was measured for all models using the final model with LOO-R^2^ tuned parameters. **Table M2** shows the parameter values evaluated per model at the final stages of results reporting.

**Table M2:** ML parameter values evaluated.

| ML Model | Tuned Parameter Values |
| --- | --- |
| RF | ntree=c(500,1000,1500,2000,2500)<br>mtry=all possibilities |
| GBM | interaction.depth=c(1:4)<br>n.trees = (1:20)*10<br>shrinkage=c(0.1,0.01, 0.02)<br>n.minobsinnode=c(2:6)<br>bag.fraction=0.5 |
| CIF | mtry=all possibilities<br>ntree*=100<br>minsplit* = 6<br>minbucket* = 3 |
| LASSO | alpha=1<br>lambda=seq(0.001,0.05,by = 0.001) |
| PLSR | ncomp = 1:15 |
| SVM | svmLinear<br>cost=seq(0.05,2,.05) |
|  | *other values besides the ones shown were optimized manually |

#### B. Consensus analysis

Consensus analysis of the relevant variables extracted from each machine learning model was done to identify consistent predictive features of quality at the early stages of manufacturing. First importance scores for all features were measured across all ML models using *varImp* with *caret* R package except for scores for SVM which *rminer* R package was used. These importance scores were percent increase in mean squared error (MSE), relative importance through average increase in prediction error when a given predictor is permuted, permuted coefficients values, absolute coefficient values, weighted sum of absolute coefficients values, and relative importance from sensitivity analysis determined for RF, GBM, CIF, LASSO, PLSR, and SVM, respectively. Using these scores, key predictive variables were selected if their importance scores were within the 80^th^ percentile ranking for the following ML methods: RF, GBM, CIF, LASSO, PLSR, SVM while for SR variables present in >30% of the top-performing SR models from DataModeler (R2≥ 90%, Complexity ≤ 100) were chosen to investigate consensus except for NMR media models at day 4 which considered a combination of the top-performing results of models excluding lactate ppms, and included those variables which were in > 40% of the best performing models. Only variables with those high percentile scoring values were evaluated in terms of their logical relation (intersection across ML models) and depicted using a Venn diagram from the *venn* R package.

## Data availability

The pre-processed set of the data used in this work is available in Supplementary Methods. All NMR data are available at the Metabolomics Workbench^42^ with DOI: http://dx.doi.org/10.21228/M8F982.

## Code availability

Machine learning implementation codes used in this work are available at GitHub (https://github.com/wandaliz/CMaT_TCell_MachineLearning/). DataModeler information can be requested at http://www.evolved-analytics.com/.

## Acknowledgments

The material is based upon work supported by the National Science Foundation under Grant No. EEC-1648035. The work and views presented are those of the authors and do not reflect the views of the National Science Foundation. The research work from N.J.D. and K.R. was also partially supported by funds from The Billie and Bernie Marcus Foundation, The Georgia Research Alliance, and the Georgia Tech Foundation through their support of the Marcus Center for Therapeutic Cell Characterization and Manufacturing (MC3M) at Georgia Tech. N.J.D. would like to thank Melissa Kemp for access to the Bioplex 200 machine and to Levi Wood/Laura Weinstock for the optimized Luminex protocol. M.B.C. would like to thank Hesam Dashti for assistance with getting additional GISSMO compound entries and simulation frequencies uploaded to enable the mixture simulation.

