## Supplementary Tables and Figures for "Predicting T Cell Quality During Manufacturing Through an Artificial Intelligence-based Integrative Multi-Omics Analytical Platform"

**Author information**

Valerie Y. Odeh-Couvertier, Nathan J. Dwarshuis, and Maxwell B. Colonna: These authors contributed equally to this work.

Affiliations

**Department of Industrial Engineering, University of Puerto Rico Mayagüez, Mayagüez, PR, 00681, USA**

Valerie Y. Odeh-Couvertier & Wandaliz Torres-Garcia

**The Wallace H. Coulter Department of Biomedical Engineering, Georgia Institute of Technology, Atlanta, GA, 30318, USA**

Nathan J. Dwarshuis & Krishnendu Roy

**Departments of Genetics and Biochemistry & Molecular Biology, Complex Carbohydrate Research Center, University of Georgia, Athens, GA, 30602, USA**

Maxwell B. Colonna & Arthur S. Edison

**Center for Cellular Immunotherapies, Perelman School of Medicine, University of Pennsylvania, Philadelphia, PA, 19104, USA**

Bruce L. Levine

**Evolved Analytics LLC, Rancho Santa Fe, CA, USA**

Theresa Kotanchek

### *Supplementary Tables*

***Data availability (Both, DOE & ADOE)***

**Supp.Table.S1:** See NatureBME_SuppTableS1.xlsx file containing the summarized data collected.

***Phase 1: Optimization of Process Parameters***

**Supp.Table.S2.** Summary Day 14 Total Live (CD4+, CD8+) $T_N$ and $T_{CM}$ cells and Ratios for DOE/ADOE

| Response | Experiment | Minimum | Median | Mean | Maximum |
|---|---|---|---|---|---|
| Total live CD4+ $T_N$ and $T_{CM}$ cells | DOE: 18-runs | $4.3 \times 10^5$ | $2.5 \times 10^6$ | $2.3 \times 10^6$ | $4.0 \times 10^6$ |
| | ADOE: 12-runs | $7.4 \times 10^5$ | $3.2 \times 10^6$ | $3.1 \times 10^6$ | $4.7 \times 10^6$ |
| Total live CD8+ $T_N$ and $T_{CM}$ cells | DOE: 18-runs | $4.1 \times 10^6$ | $1.2 \times 10^7$ | $1.1 \times 10^7$ | $1.9 \times 10^7$ |
| | ADOE: 12-runs | $7.8 \times 10^6$ | $1.1 \times 10^6$ | $1.2 \times 10^6$ | $1.8 \times 10^7$ |
| Ratio live CD4+/CD8+ $T_N$ and $T_{CM}$ cells | DOE: 18-runs | 0.09 | 0.21 | 0.20 | 0.34 |
| | ADOE: 12-runs | 0.09 | 0.29 | 0.27 | 0.49 |

*Phase 2: Single and Multi-omics Integrative Modeling*

**Supp.Table.S3.** Variables present in >30% of the top-performing Symbolic Regression models from DataModeler (R2≥ 90%, Complexity ≤ 100) for the different end-product responses except for NMR media models at day 4 which chose variables present in >= 40% of various models.

| **Input** | **Ratio CD4+/CD8+ $T_N$+$T_{CM}$ cells** | **Total Live CD4+ $T_N$+$T_{CM}$ cells** | **Total Live CD8+ $T_N$+$T_{CM}$ cells** |
|---|---|---|---|
| **PP +N4** | DMS Conc<br>Functional Mabs %<br>Lactate<br>Formate<br>Histidine<br>Ethanol | IL2 Conc<br>DMS Conc<br>Functional Mab %<br>Lactate<br>Formate<br>Ethanol | IL2 Conc<br>DMS Conc<br>Lactate<br>Formate<br>Histidine<br>BCAAs<br>Ethanol |
| **PP+N6** | Functional Mabs%<br>DMS conc<br>UK 75387<br>Dimethylamine<br>Glycine<br>UK 13653<br>Lactate<br>Histidine | IL2 Conc<br>Functional Mab %<br>DMS Conc<br>Lactate<br>Phenylalanine<br>UK41784 | IL2 Conc<br>DMS Conc<br>Ethanol<br>UK 75387<br>Tyrosine<br>Pyruvate<br>Lactate |
| **PP+S6** | DMS conc<br>GMCSF<br>IL2R<br>MIF<br>IL5<br>Functional Mabs %<br>TNFa | IL Conc<br>IL2R<br>IL13<br>IL15<br>IL17a<br>IFN alpha<br>MIF | IL2 Conc<br>DMS Conc<br>IL15<br>IL17a<br>TNFa |
| **PP+S6+N6** | TNFa<br>IL3<br>Formate<br>Tyrosine<br>GMCSF<br>UK75387 | IL2 Conc<br>IL2R<br>IL13<br>IL15<br>Glycine<br>Histidine<br>IL17a<br>MIF<br>IFN alpha | IL2 Conc<br>IL15<br>DMS Conc<br>IL17a<br>GM CSF<br>UK 15208<br>IL2R<br>TNFa |

**Supp.Table.S4.** Variable combinations across responses from top-performing SR – DataModeler

| Response | Predictors | Top Symbolic Regression DataModeler Combinations |
|---|---|---|
| **Ratio CD4+/CD8+ $T_N$+$T_{CM}$ cells** | **PP +N4** | DMS Conc + Functional Mabs %+Histidine+Formate+Lactate |
| | | IL2 Conc+DMS Conc + Functional Mabs %+Histidine+Formate+Ethanol |
| | | IL2 Conc+DMS Conc + Functional Mabs %+Histidine+Lactate |
| | **PP+N6** | DMS Conc + Functional Mabs%+UK 1.3653+ Dimethylamine+ Glycine+UK 7.5387 |
| | | DMS Conc +Functional Mabs%+ Lactate1+Histidine +UK 7.5387 |
| | **PP+S6** | DMS Conc + Functional Mabs %+GMCSF+IL2R+MIF |
| | | DMS Conc +GMCSF+IL2R+ IL5+MIF |
| | | DMS Conc + Functional Mabs %+GMCSF+TNFa |
| | **PP+S6+N6** | GMCSF+ IL3+TNFa+ Tyrosine+Formate |
| | | IL3+TNFa+ Tyrosine+Formate |
| | | IL3+TNFa+Formate+UK 7.5387 |
| **Total Live CD4+ $T_N$+$T_{CM}$ cells** | **PP +N4** | IL2 Conc+DMS Conc+Functional Mabs%+Ethanol+Lactate |
| | | IL2 Conc+DMS Conc+Functional Mabs%+Ethanol+Formate |
| | | IL2 Conc+DMS Conc+Functional Mabs%+Ethanol+Lactate+Formate |
| | | IL2 Conc+DMS Conc+Functional Mabs%+Ethanol+Formate+Histidine |
| | | IL2 Conc+Functional Mabs%+Ethanol+Dimethylamine+Lactate |
| | **PP+N6** | IL2 Conc + DMS Conc + Functional Mabs% + Lactate + Phenylalanine |
| | | IL2 Conc + DMS Conc + Functional Mabs%+UK 4.1784 |
| | **PP+S6** | IL2 Conc + IL13 + IL15 + IL17a + IL2R |
| | | IL2 Conc + IFN alpha+ IL13 + IL15 + IL2R |
| | | IL2 Conc + IL13 + IL15 + IL2R |
| | **PP+S6+N6** | IL2 Conc+IFN Alpha+ IL13+IL15+Histidine |
| | | IL2 Conc+ IL13+IL15+IL17a+IL2R+Glycine |
| | | IL2 Conc+IL13+IL15+IL2R+MIF+Glycine |
| **Total Live CD8+ $T_N$+$T_{CM}$ cells** | **PP +N4** | IL2 Conc + DMS Conc + Lactate + Ethanol + Histidine + BCAAs |
| | | IL2 Conc + DMS Conc + Functional Mabs% + Lactate + Ethanol + Histidine |
| | | IL2 Conc + DMS Conc + Formate + Ethanol + Glucose + BCAAs |
| | | IL2 Conc + DMS Conc + Formate + Glucose + BCAAs |
| | | IL2 Conc + DMS Conc + Ethanol + Lactate+ Glutamine |
| | **PP+N6** | IL2 Conc + DMS Conc + Ethanol + Pyruvate + UK 7.5387 |
| | | IL2 Conc + DMS Conc + Ethanol +Tyrosine |
| | | IL2 Conc + DMS Conc + Ethanol +Lactate |
| | **PP+S6** | IL2 Conc+ DMS Conc+ IL15+IL17a+TNFa |
| | **PP+S6+N6** | IL2 Conc +DMS Conc + GM CSF+IL15+IL17a+UK 1.5208 |
| | | IL2 Conc +DMS Conc + GM CSF+IL15+IL17a+IL2R |
| | | IL2 Conc +DMS Conc +IL15+IL17a+TNFa |

### *Supplementary Figures*

***Phase 1: Optimization of Process Parameters***

Supp.Fig.S1. Symbolic regression ensemble plots as given by DataModeler optimizing for Total live $CD4^+$ $T_N$+$T_{CM}$ cells.

Supp.Fig.S2. Symbolic regression ensemble plots as given by DataModeler optimizing for Total live CD8+ $T_N$+$T_{CM}$ cells.

Supp.Fig.S3. Single-response model maximum optimization plots by end-point response.

Supp.Fig.S4. Two-way process parameters interactions response contour plots at model maximum

Supp.Fig.S5. Predicted response profiles of a) Total live $CD4^+$ $T_N$+$T_{CM}$ cells versus ratio of $CD4^+$ to $CD8^+$ $T_N$+$T_{CM}$ cells (red is low and green is high value of the DMS process parameter) at the predicted optimum DMS process conditions for total live CD4+ TN+TCM cells – Part I.

Supp.Fig.S6. Predicted response profiles of a) Total live $CD4^+$ $T_N$+$T_{CM}$ cells, b) Total Live $CD8^+$ $T_N$+$T_{CM}$ cells, and c) Ratio of $CD4^+$ to $CD8^+$ $T_N$+$T_{CM}$ cells at the predicted optimum DMS process conditions for Total live $CD4^+$ $T_N$+$T_{CM}$ cells – Part II.

Supp.Fig.S7. Single-response model maximum optimization plots by end-point response

Supp.Fig.S8. Predicted response contour profiles of a) Total live CD4+ $T_N$+$T_{CM}$ cells and b) Ratio of CD4+ to CD8+ $T_N$+$T_{CM}$ cells at the predicted optimum DMS process conditions for Total live CD4+ $T_N$+$T_{CM}$ cells –IL2 Conc=40, DMS Conc=3500, Functional MAB %=100.

Supp.Fig.S9. Predicted response profiles of a) Total live CD4+ $T_N$+$T_{CM}$ cells and b) Ratio of CD4+ to CD8+ $T_N$+$T_{CM}$ cells at the predicted optimum DMS process conditions for Total live CD4+ $T_N$+$T_{CM}$ cells –IL2 Conc=40, DMS Conc=3500, Functional MAB %=100.

Supp.Fig.S10. Predicted response profiles of a) Total live CD4+ $T_N$+$T_{CM}$ cells and b) Ratio of CD4+ to CD8+ $T_N$+$T_{CM}$ cells at the conditions of IL2 Conc=40, DMS Conc=500, Functional MAB %=100.

Supp.Fig.S11. Predicted response profiles of a) Total live CD4+ $T_N$+$T_{CM}$ cells and b) Ratio of CD4+ to CD8+ $T_N$+$T_{CM}$ cells at the conditions of IL2 Conc=40, DMS Conc=2500, Functional MAB %=100.

***Phase 1: Sequential Experimentation based on Optimization Models: Data Characterization***

Supp.Fig.S12. Process parameters impact on end-product responses across the two sequential experiments (DOE, ADOE).

Supp.Fig.S13. Pairwise plots of Total live $CD4^+$ $T_N$+$T_{CM}$ cells.

Supp.Fig.S14. Heatmap display of hierarchical clustering for all three $T_N$+$T_{CM}$ endpoint responses

Supp.Fig.S15. Pearson correlation structure of cytokines day 6 with responses

Supp.Fig.S16. Correlation structure of NMR media features day 4 (a) & 6 (b) with target responses

***Phase 2: Multi-Omics Integrative Approach for Early Predictive Signatures***

Supp.Fig.S17. Multi-omics prediction profiles at day 6 using Symbolic Regression from DataModeler.

Supp.Fig.S18. Overall feature consensus analysis of top-performing features in multi-omics models at day 6
Supp.Fig.S19. Overall feature consensus analysis of top-performing features in single-omics (NMR) models at day 4
Supp.Fig.S20. Overall feature consensus analysis of top-performing features in single-omics (Cytokine-S6) models at day 6
Supp.Fig.S21. Parameter tuning for Ratio of $CD4^+$ to $CD8^+$ $T_N$+$T_{CM}$ cells across all ML models from multi-omics integration at day 6.
Supp.Fig.S22. Parameter tuning for Total live $CD4^+$ $T_N$+$T_{CM}$ cells across all ML models from multi-omics integration at day 6.
Supp.Fig.S23. Parameter tuning for Total live $CD8^+$ $T_N$+$T_{CM}$ cells across all ML models from multi-omics integration at day 6.

***Phase 2: Early Predictive Signatures using NMR Day 4 Modeling***

Supp.Fig.S24. Variance based feature selection of NMR features for computational modeling.
Supp.Fig.S25. Media NMR intensities across monitoring times for total live $CD4^+$ $T_N$+$T_{CM}$ cells.
Supp.Fig.S26. Media NMR intensities across monitoring times for ratio $CD4^+$/$CD8^+$ $T_N$+$T_{CM}$ cells.
Supp.Fig.S27: Media NMR intensities across monitoring times for total live $CD8^+$ $T_N$+$T_{CM}$ cells.
Supp.Fig.S28: Predicted response profiles at the predicted optimum DMS process conditions for Total live $CD4^+$ $T_N$+$T_{CM}$ cells for NMR Models at Day 4 using DataModeler - View 1.
Supp.Fig.S29: Predicted response profiles for NMR Models at Day 4 using DataModeler - View 2.
Supp.Fig.S30: Predicted response profiles for NMR Models at Day 4 using DataModeler - View 3.
Supp.Fig.S31: Response contour plots for NMR media analysis at day 4 using DataModeler – View 1
Supp.Fig.S32: Response contour plots for NMR media analysis at day 4 using DataModeler – View 2
Supp.Fig.S33: Response contour plots for NMR media analysis at day 4 using DataModeler – View 3
Supp.Fig.S34: Response contour plots for NMR media analysis at day 4 using DataModeler – View 4
Supp.Fig.S35: Response contour plots for NMR media analysis at day 4 using DataModeler – View 5
Supp.Fig.S36: Response contour plots for NMR media analysis at day 4 using DataModeler – View 6
Supp.Fig.S37: Response contour plots for NMR media analysis at day 4 using DataModeler – View 6
Supp.Fig.S38: NMR Feature Correlation for SR-DataModeler models for NMR media analysis at day 4
Supp.Fig.S39: Bivarate Plot of NMR features predictive of $CD4^+$ $T_N$+$T_{CM}$ cells, $CD8^+$ $T_N$+$T_{CM}$ cells, and ratio $CD4^+$ $T_N$+$T_{CM}$ to $CD8^+$ $T_N$+$T_{CM}$ cells.

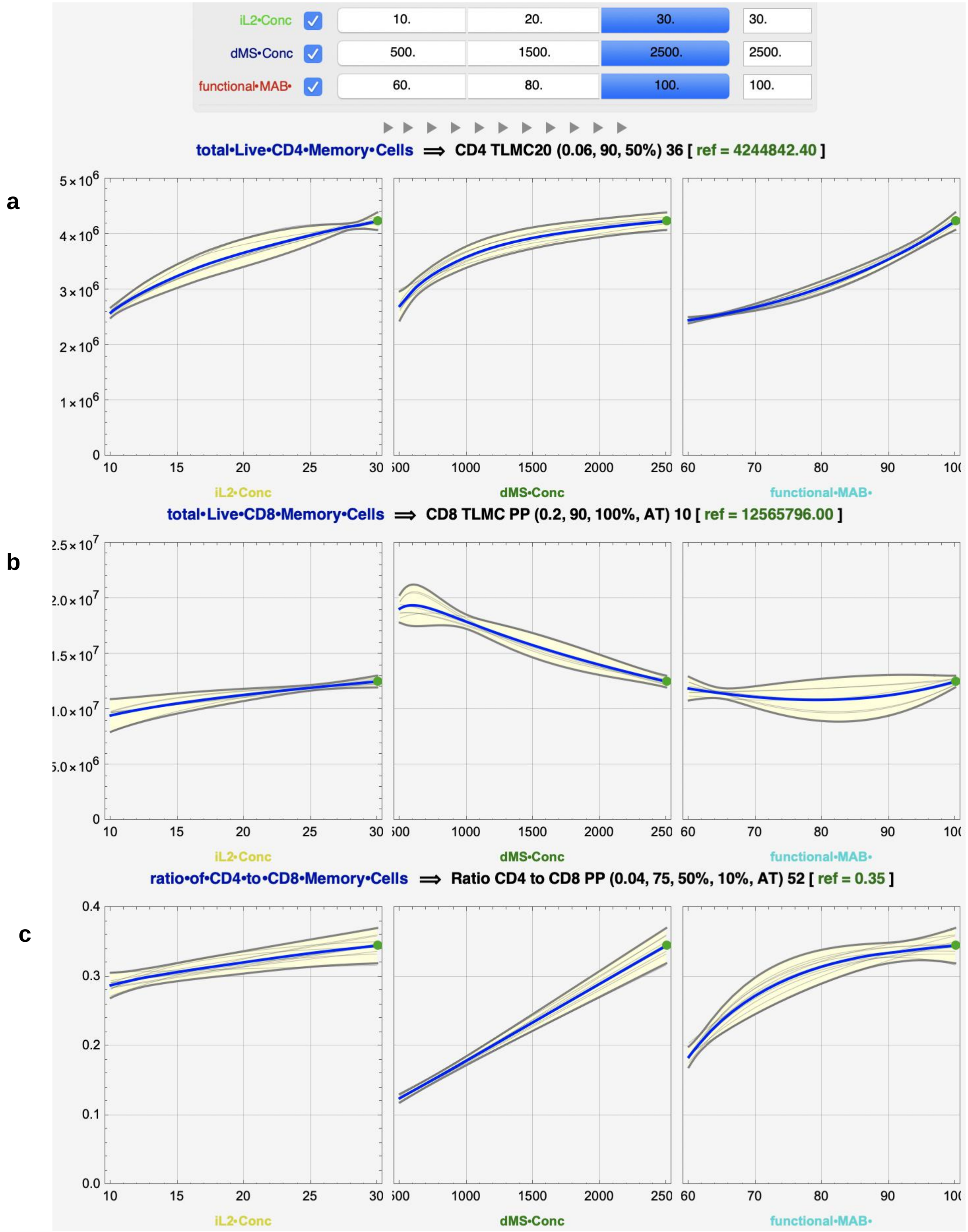


**Supp.Fig.S1. Symbolic regression ensemble plots as given by DataModeler optimizing for Total live $CD4^{+}$ $T_{N}$+$T_{CM}$ cells.** Predicted response profiles of a) Total live $CD4^{+}$ $T_{N}$+$T_{CM}$ cells, b) Total live $CD8^{+}$ $T_{N}$+$T_{CM}$ cells and c) Ratio of $CD4^{+}$ to $CD8^{+}$ $T_{N}$+$T_{CM}$ cells at the predicted optimum for Total live $CD4^{+}$ $T_{N}$+$T_{CM}$ cells.

a

b

c

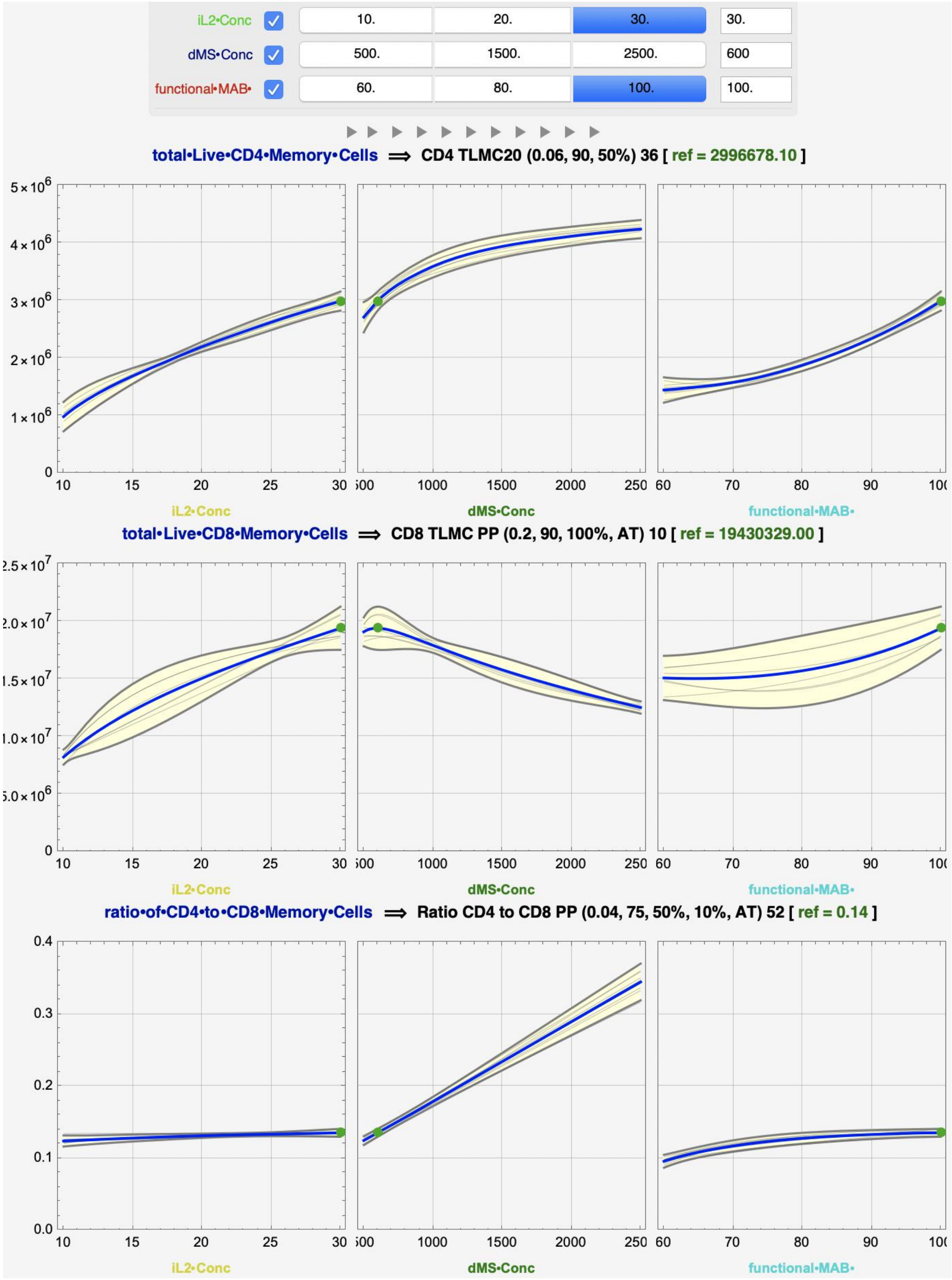


**Supp.Fig.S2. Symbolic regression ensemble plots as given by DataModeler optimizing for Total live CD8⁺ $T_N$+$T_{CM}$ cells.** Predicted response profiles of a) Total live CD4⁺ $T_N$+$T_{CM}$ cells, b) Total live CD8⁺ $T_N$+$T_{CM}$ cells and c) Ratio of CD4⁺ to CD8⁺ $T_N$+$T_{CM}$ cells at the predicted optimum for Total live CD8⁺ $T_N$+$T_{CM}$ cells.

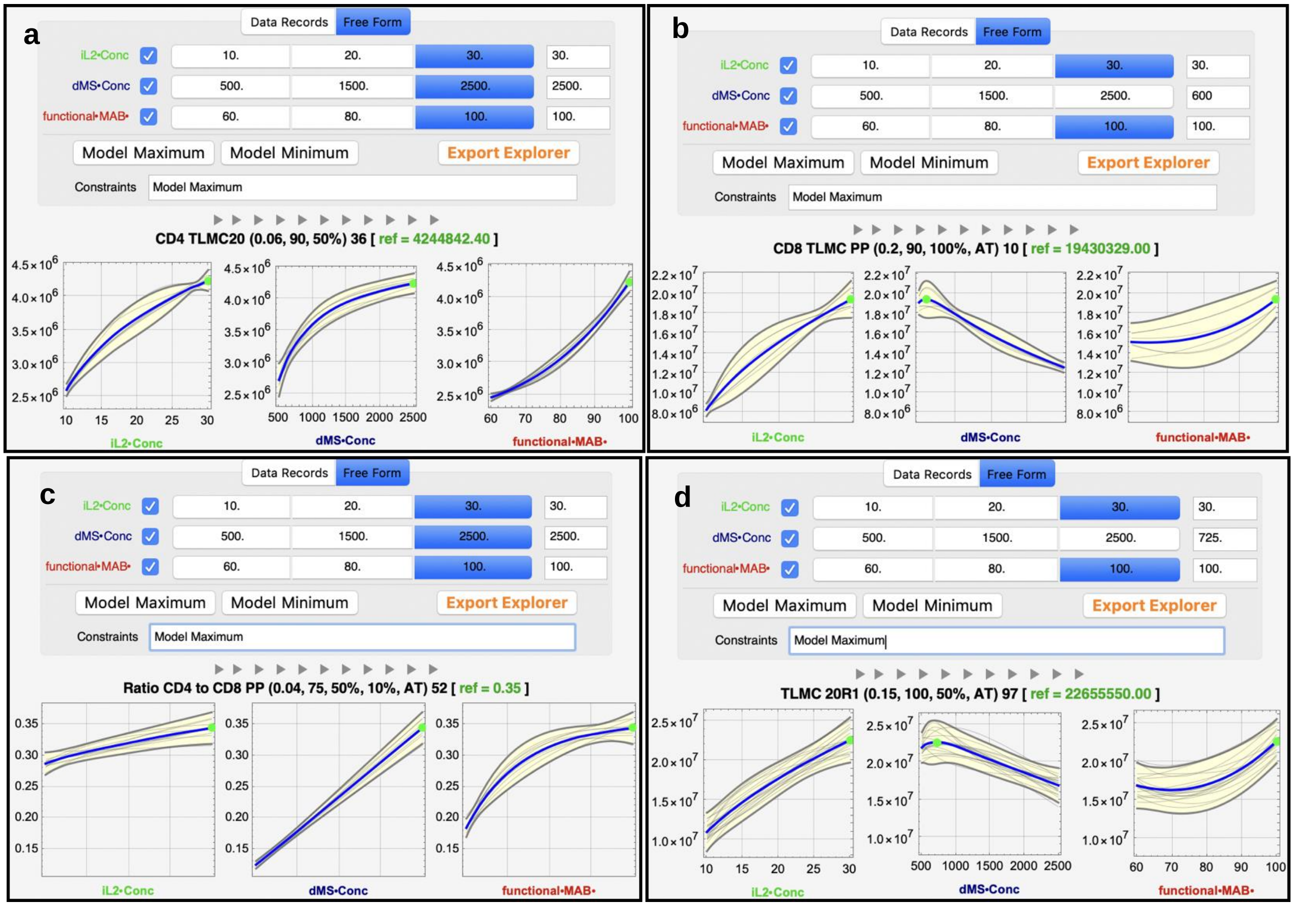


**Supp.Fig.S3. Single-response model maximum optimization plots by end-point response**: a) Total live CD4$^{+}$ $T_N$+ $T_{CM}$ cells, b) Total live CD8$^{+}$ $T_N$+$T_{CM}$ cells c) Ratio of CD4$^{+}$ to CD8$^{+}$ $T_N$+$T_{CM}$ cells, and d) Total live $T_N$+$T_{CM}$ cells.

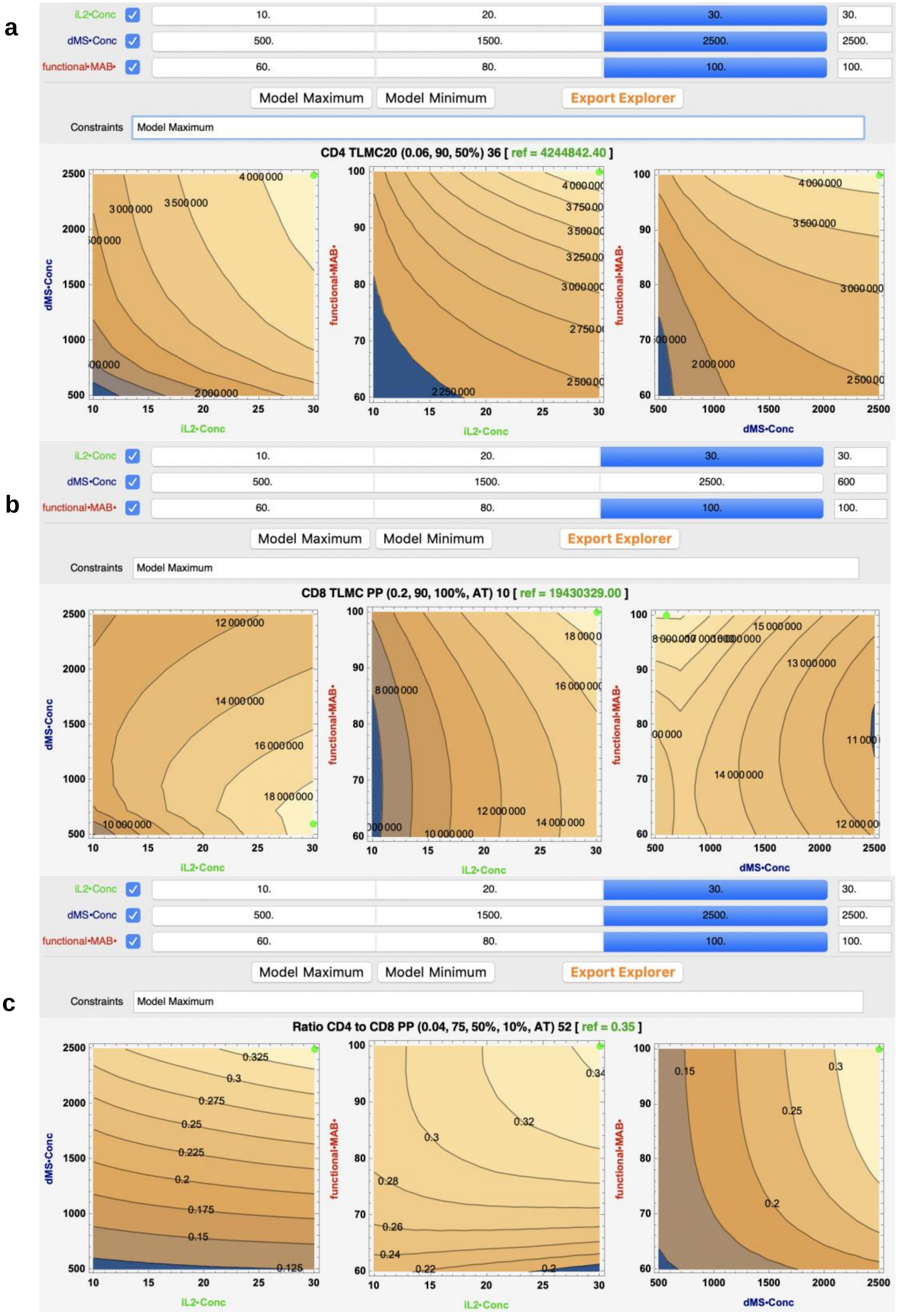


**Supp.Fig.S4. Two-way process parameters interactions response contour plots at model maximum** for: a) Total live CD4$^{+}$ $T_N$+$T_{CM}$ cells, b) Total live CD8$^{+}$ $T_N$+$T_{CM}$ cells, and c) Ratio CD4$^{+}$/CD8$^{+}$ $T_N$+$T_{CM}$ cells.

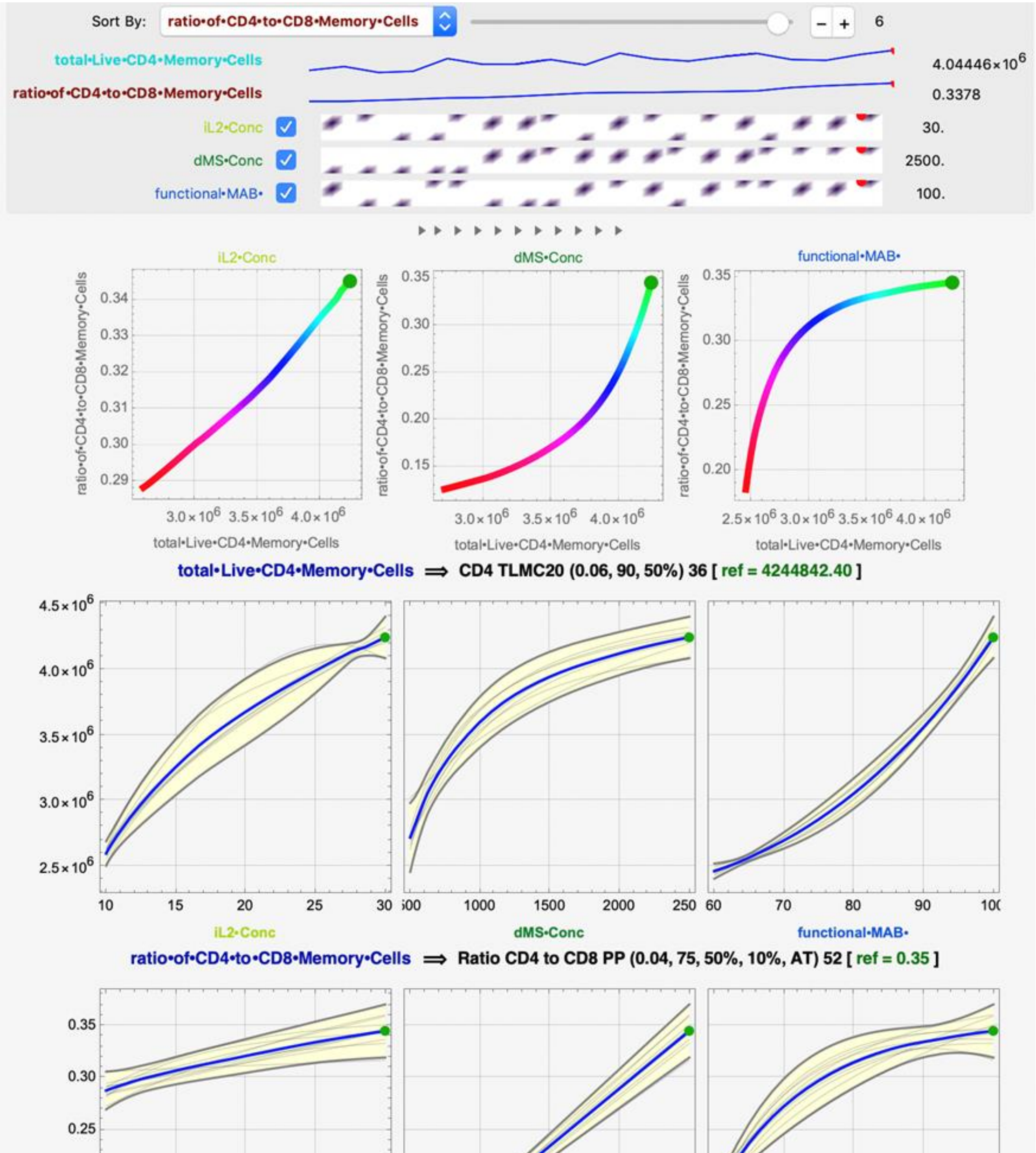


**Supp.Fig.S5. Predicted response profiles of a) Total live CD4$^+$ T$_N$+T$_{CM}$ cells versus ratio of CD4$^+$ to CD8$^+$ T$_N$+T$_{CM}$ cells (red is low and green is high value of the DMS process parameter) at the predicted optimum DMS process conditions for total live CD4$^+$ T$_N$+T$_{CM}$ cells – Part I.** At this setting, the predicted value of total live CD4$^+$ naïve-memory yield was 4.2 x $10^6$ and the predicted ratio of CD4+ to CD8+ T$_N$+T$_{CM}$ cells was 0.35. The measured value of total live CD4+ naïve-memory yield was 4.0 x $10^6$ and the observed ratio of CD4+ to CD8+ T$_N$+T$_{CM}$ cells was 0.34 for samples from the initial experiment (DOE).

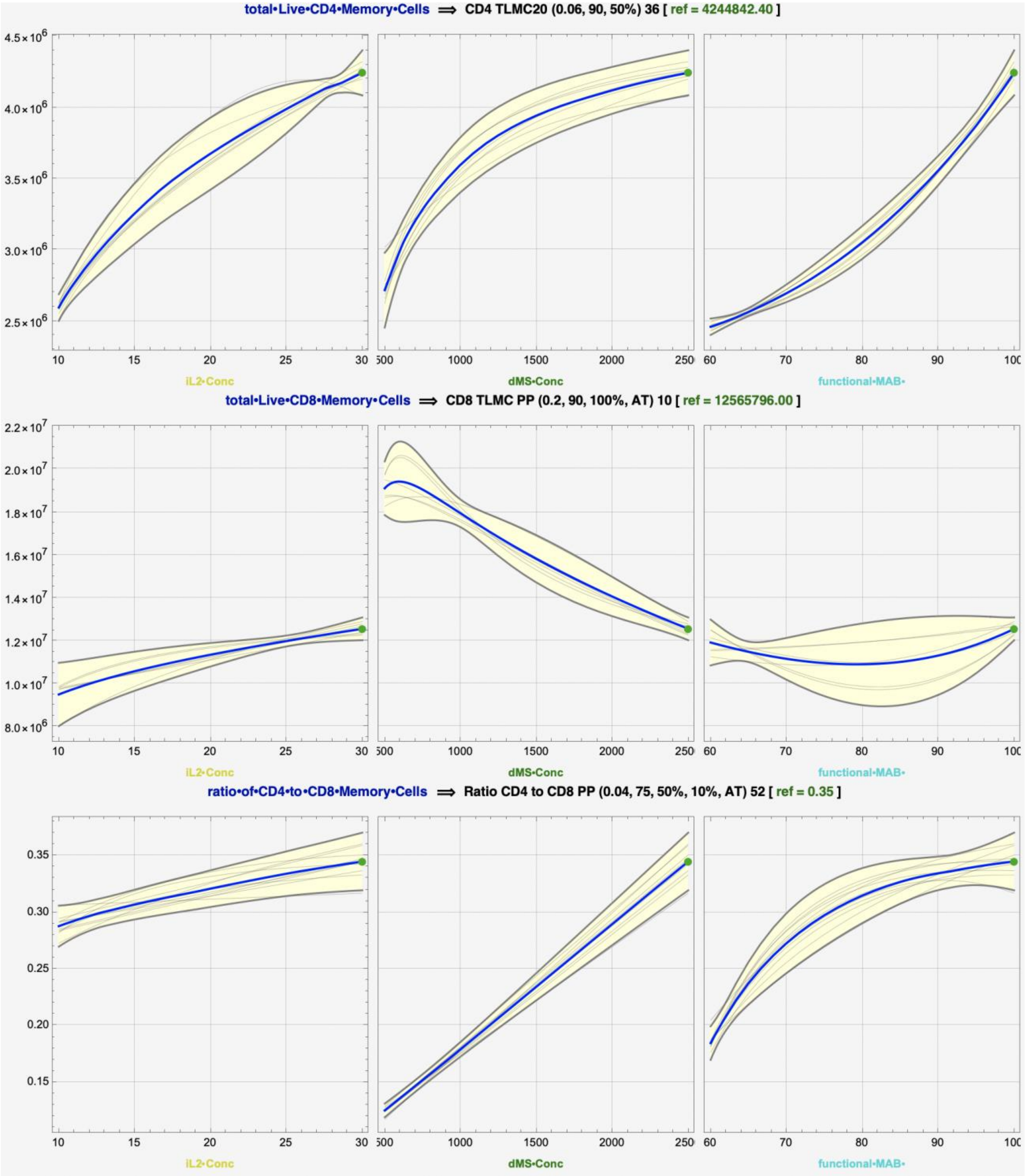


**Supp.Fig.S6. Predicted response profiles of Total live CD4$^+$ $T_N$+$T_{CM}$ cells, Total Live CD8$^+$ $T_N$+$T_{CM}$ cells, and Ratio of CD4$^+$ to CD8$^+$ $T_N$+$T_{CM}$ cells at the predicted optimum DMS process conditions for Total live CD4$^+$ $T_N$+$T_{CM}$ cells – Part II.** At this setting, the predicted value of Total live CD4$^+$ $T_N$+$T_{CM}$ cells is 4.4 x $10^6$, and the predicted values of Total live CD8$^+$ naïve-memory yield and Ratio of CD4$^+$ to CD8$^+$ $T_N$+$T_{CM}$ cells are 1.2 x $10^7$ and 0.35, respectively.

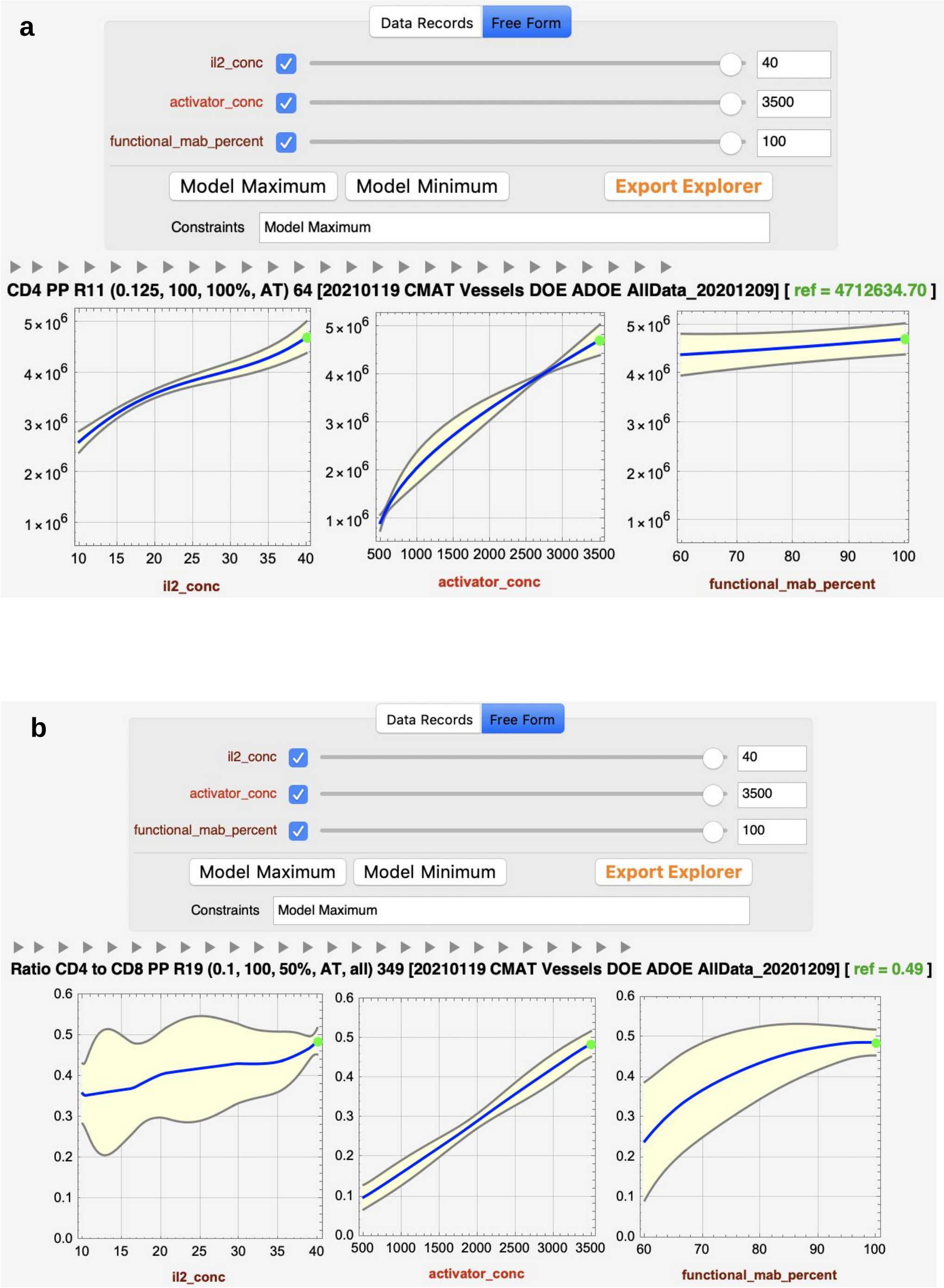


**Supp.Fig.S7. Single-response model maximum optimization plots by end-point response**: a) Total live CD4+ $T_N$+$T_{CM}$ cells and b) Ratio of CD4+ to CD8+ $T_N$+$T_{CM}$ cells.

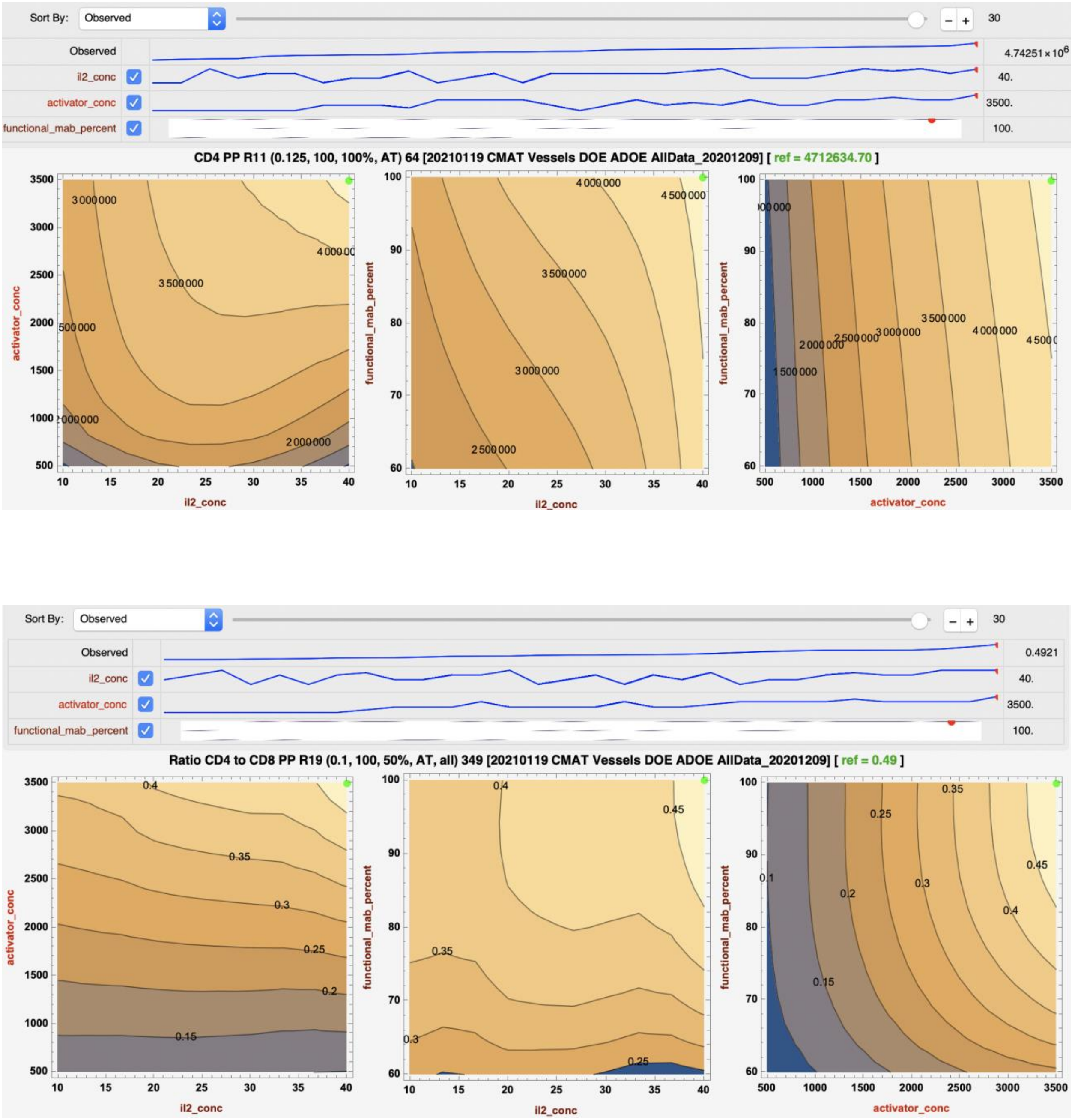


**Supp.Fig.S8. Predicted response contour profiles of a) Total live CD4+ $T_N$+$T_{CM}$ cells and b) Ratio of CD4+ to CD8+ $T_N$+$T_{CM}$ cells at the predicted optimum DMS process conditions for Total live CD4+ $T_N$+$T_{CM}$ cells –IL2 Conc=40, DMS Conc=3500, Functional MAB %=100.** At this setting, the observed values are identical to the 4.7 x $10^6$ predicted optimal value of Total live CD4+ $T_N$+$T_{CM}$ cells and the 0.49 predicted value of the Ratio of CD4+ to CD8+ $T_N$+$T_{CM}$ cells.

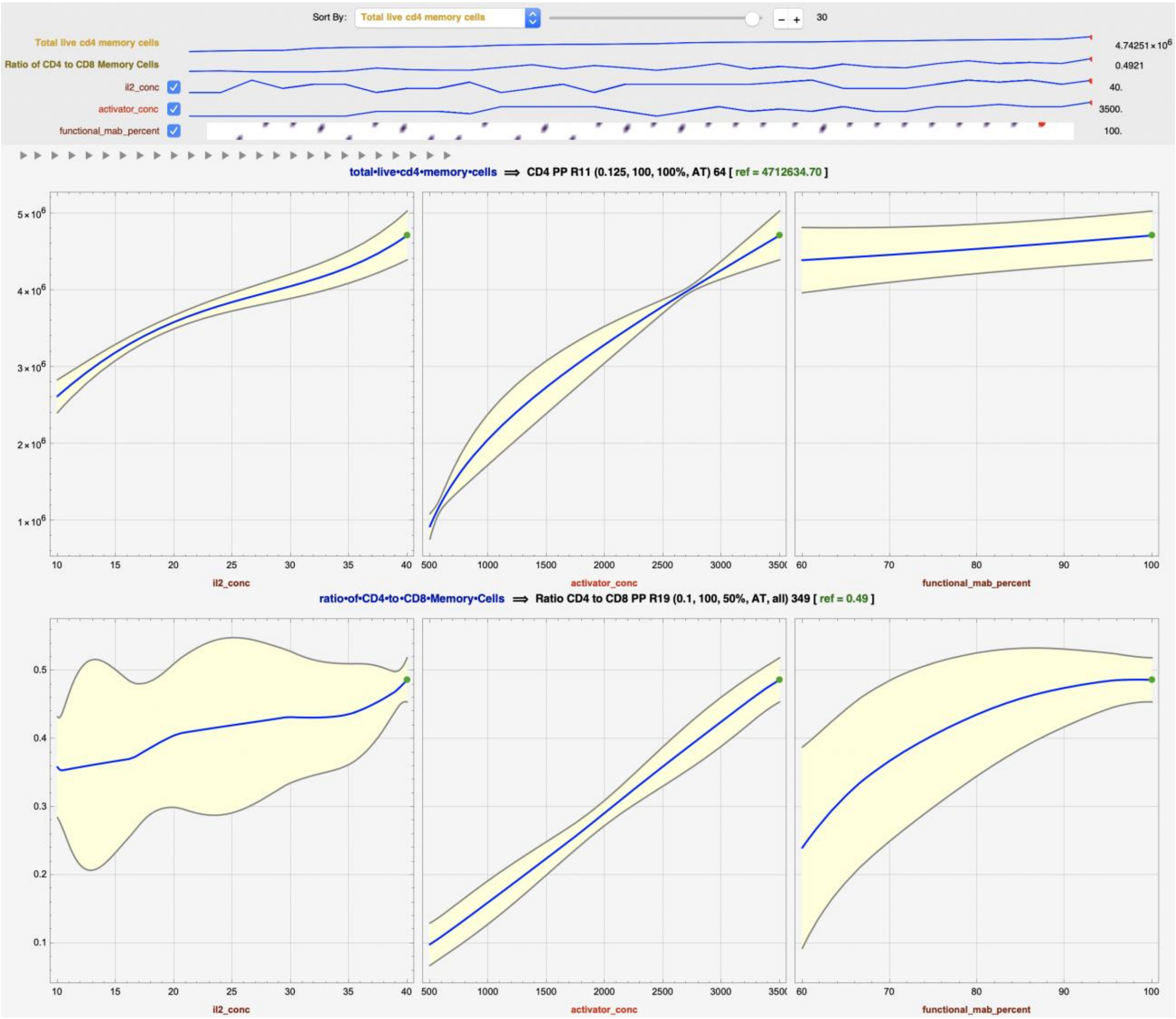


**Supp.Fig.S9. Predicted response profiles of a) Total live CD4+ $T_N$+$T_{CM}$ cells and b) Ratio of CD4+ to CD8+ $T_N$+$T_{CM}$ cells at the predicted optimum DMS process conditions for Total live CD4+ $T_N$+$T_{CM}$ cells –IL2 Conc=40, DMS Conc=3500, Functional MAB %=100.** At this setting, the observed values are identical to the 4.7 x $10^6$ predicted value of Total live CD4+ $T_N$+$T_{CM}$ cells and the 0.49 predicted value of the Ratio of CD4+ to CD8+ $T_N$+$T_{CM}$ cells.

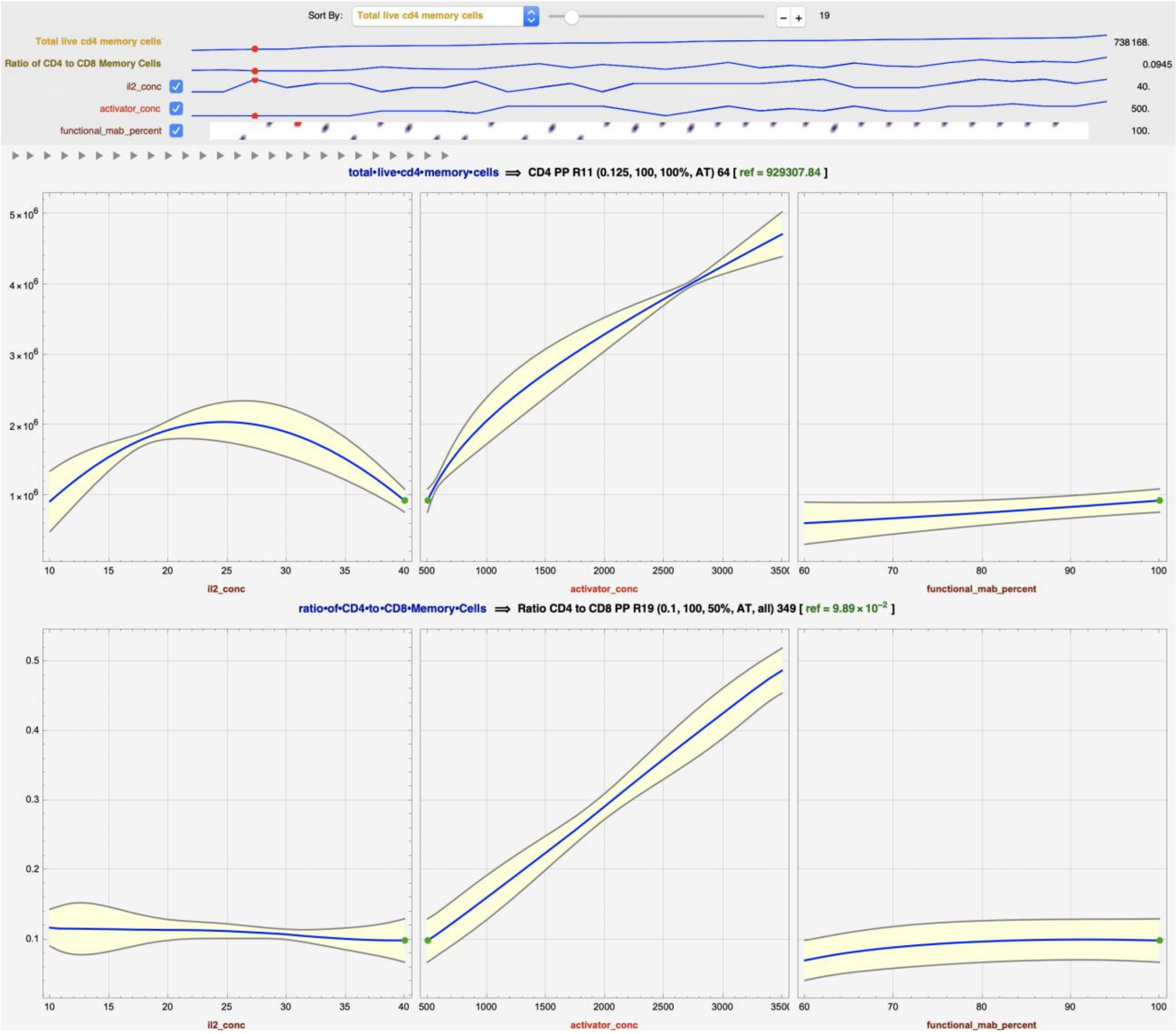


**Supp.Fig.S10. Predicted response profiles of a) Total live CD4+ $T_N$+$T_{CM}$ cells and b) Ratio of CD4+ to CD8+ $T_N$+$T_{CM}$ cells at the conditions of IL2 Conc=40, DMS Conc=500, Functional MAB %=100.** At this setting, the predicted value of Total live CD4+ $T_N$+$T_{CM}$ cells is 0.9 x $10^6$, and the predicted value of the Ratio of CD4+ to CD8+ $T_N$+$T_{CM}$ cells is 0.10. The observed value of Total live CD4+ $T_N$+$T_{CM}$ cells is 0.7 x $10^6$, and the observed value of the Ratio of CD4+ to CD8+ $T_N$+$T_{CM}$ cells is 0.09.

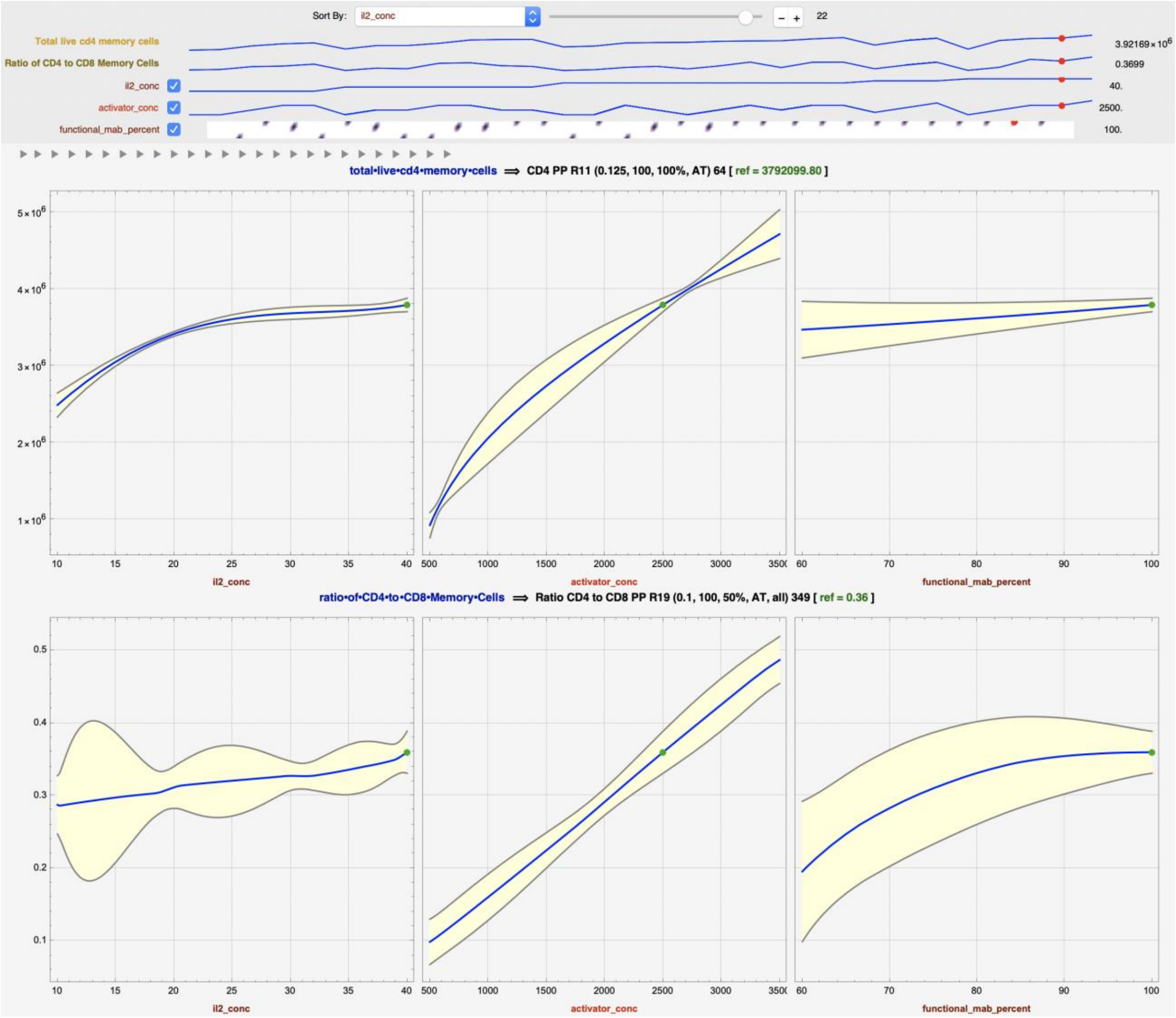


**Supp.Fig.S11. Predicted response profiles of a) Total live CD4+ $T_N$+$T_{CM}$ cells and b) Ratio of CD4+ to CD8+ $T_N$+$T_{CM}$ cells at the conditions of IL2 Conc=40, DMS Conc=2500, Functional MAB %=100.** At this setting, the predicted value of Total live CD4+ $T_N$+$T_{CM}$ cells is 3.8 x $10^6$, and the predicted value of the Ratio of CD4+ to CD8+ $T_N$+$T_{CM}$ cells is 0.36 respectively. The observed value of Total live CD4+ $T_N$+$T_{CM}$ cells is 3.9 x $10^6$, and the observed value of the Ratio of CD4+ to CD8+ $T_N$+$T_{CM}$ cells is 0.37.

***Phase 1: Sequential Experimentation based on Optimization Models: Data Characterization***

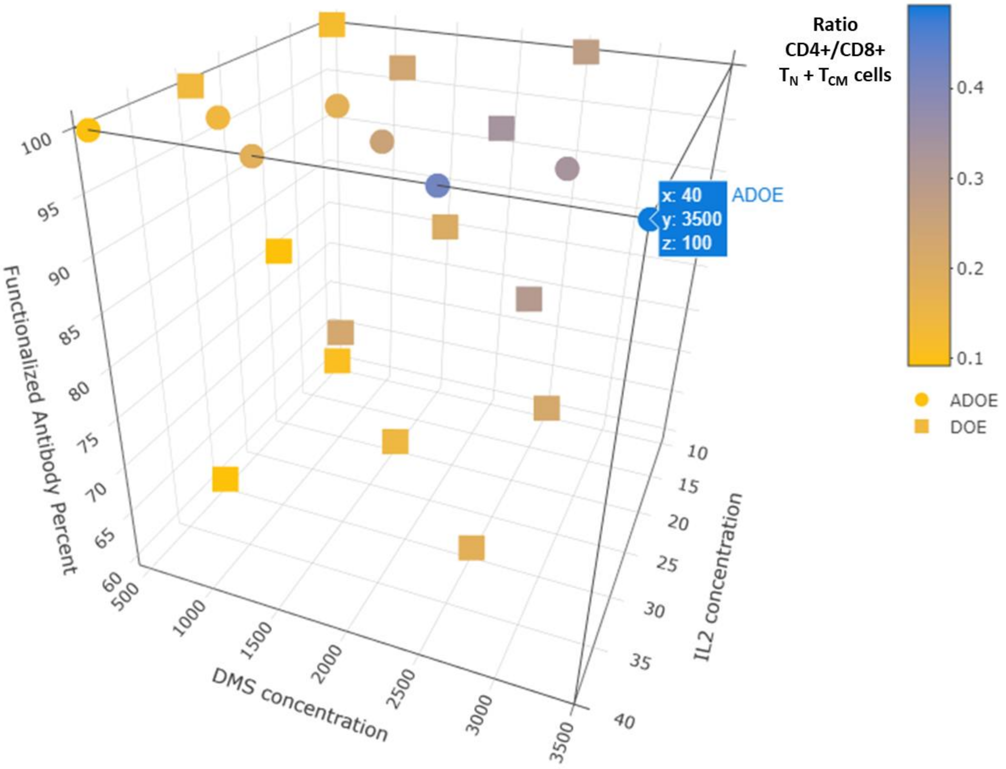


a

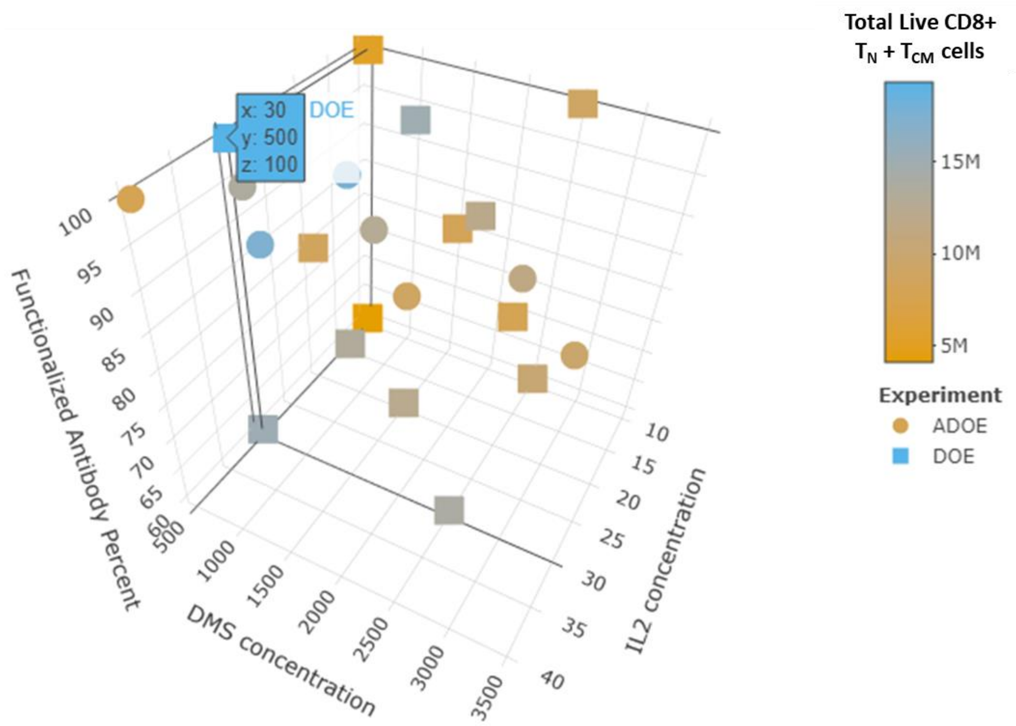


b

**Supp.Fig.S12. Process parameters impact on end-product responses across the two sequential experiments (DOE, ADOE)**. a) Ratio $CD4^{+}/CD8^{+}$ $T_N+T_{CM}$ cells, and b) Total Live $CD8^{+}$ $T_N+T_{CM}$ cells.

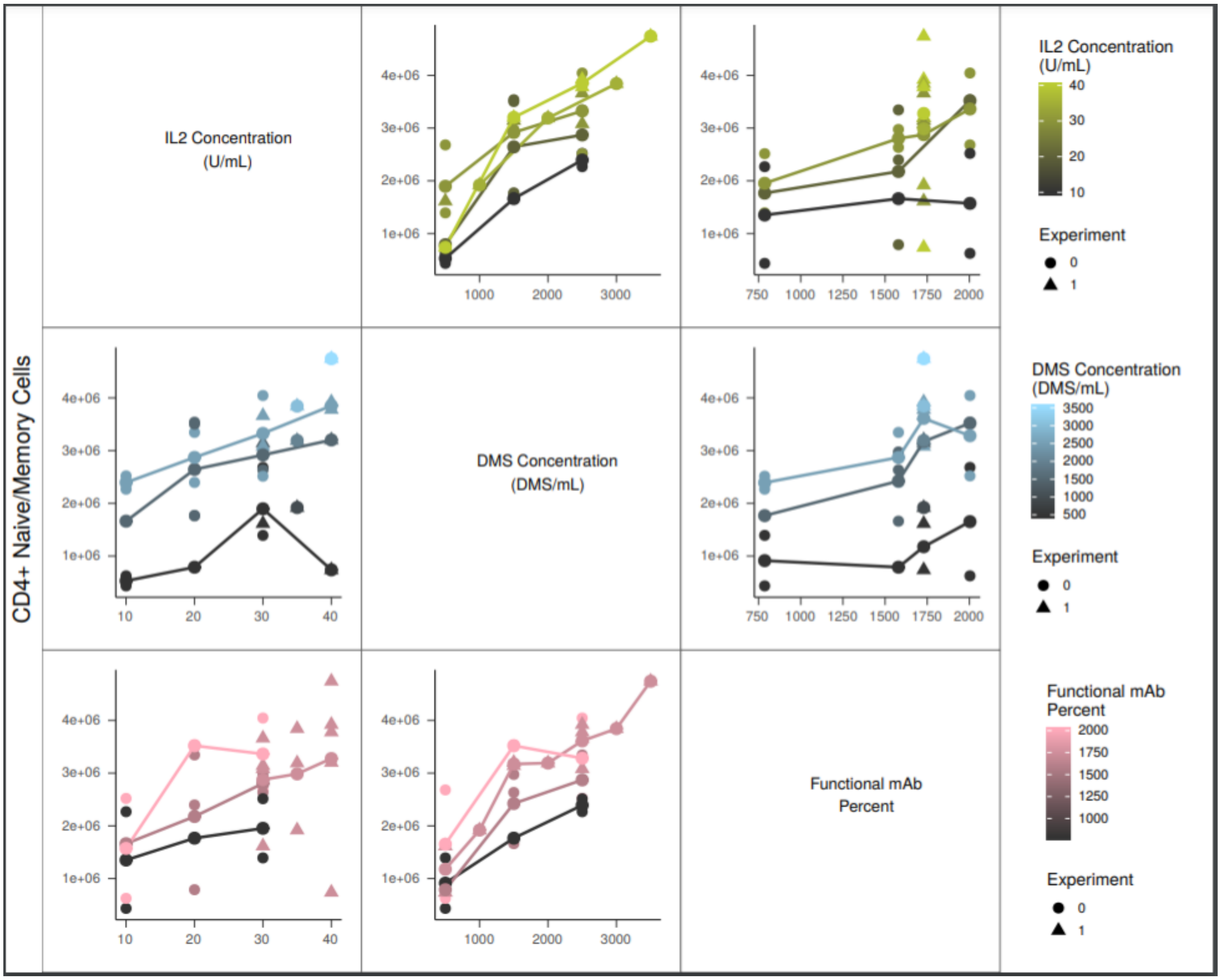


**Supp.Fig.S13. Pairwise plots of Total live CD4⁺ $T_N$+$T_{CM}$ cells.**

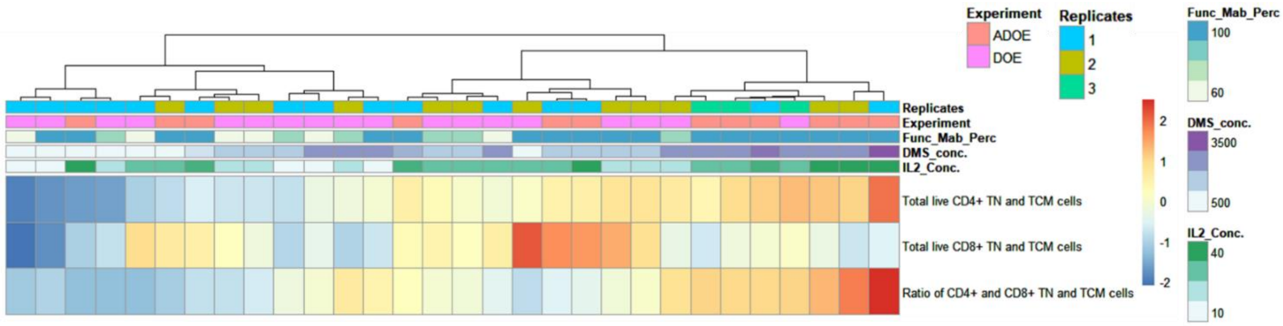


**Supp.Fig.S14. Heatmap display of hierarchical clustering for all three $T_N$+$T_{CM}$ endpoint responses** using Ward.D agglomeration and Euclidean distance.

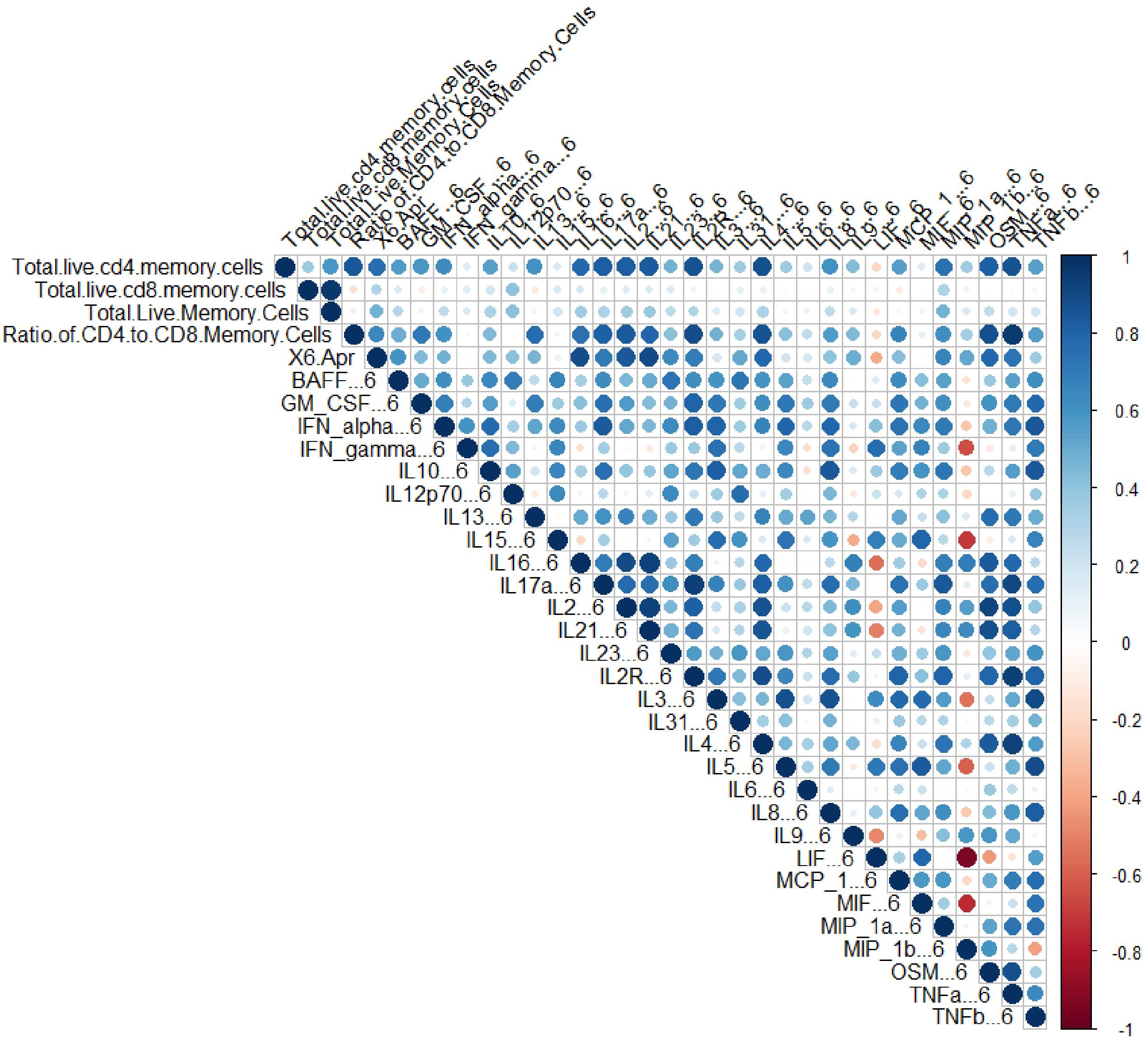


**Supp.Fig.S15. Pearson correlation structure of cytokines day 6 with responses**

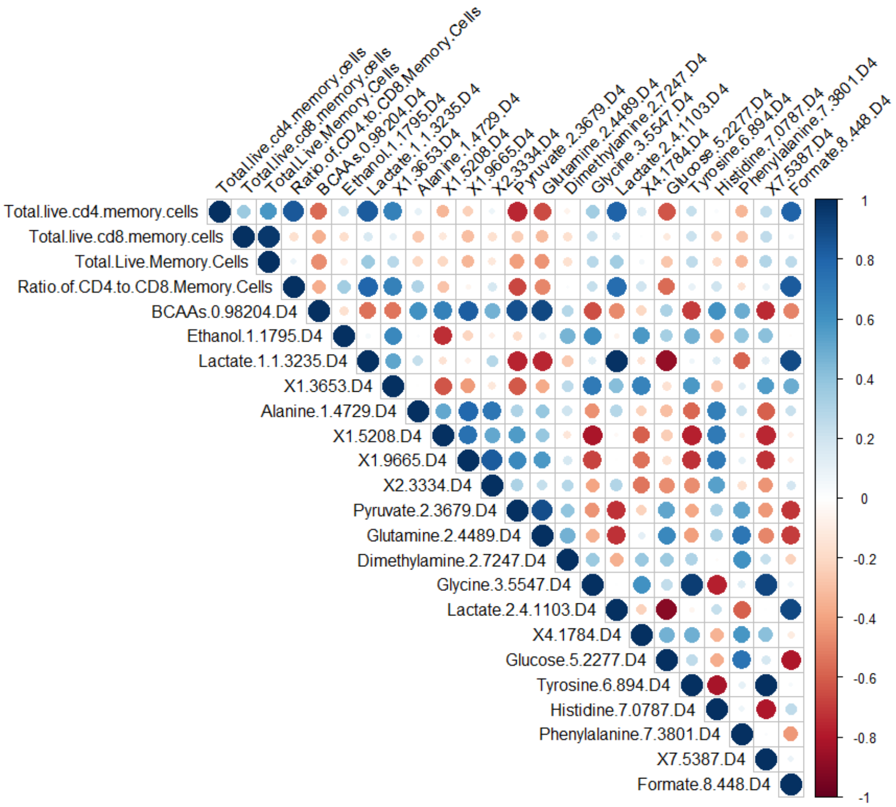


**a**

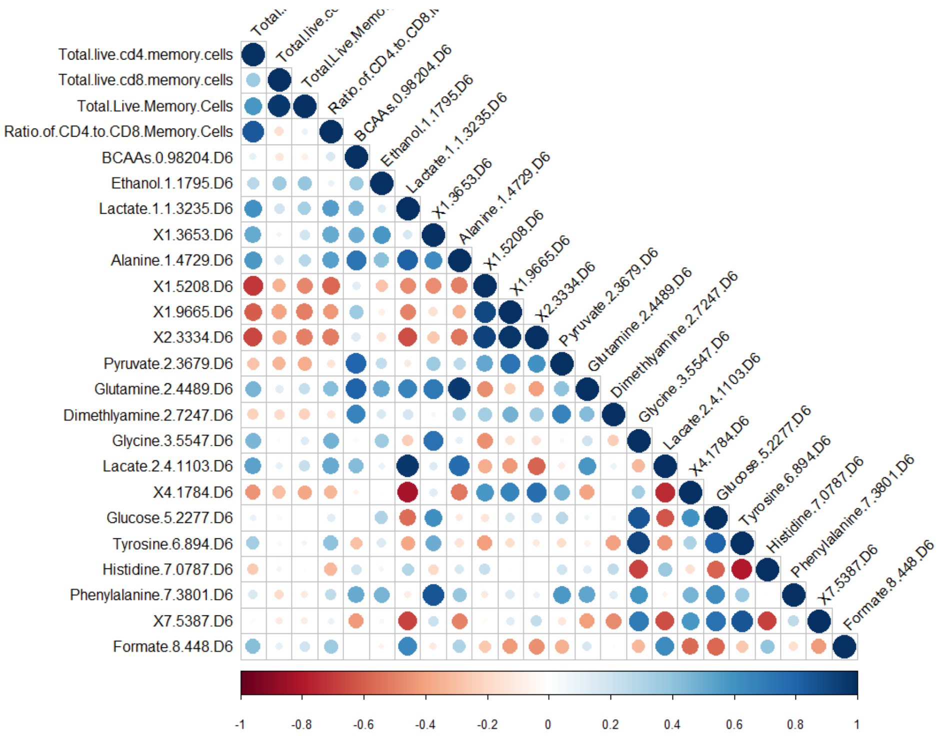


**b**

**Supp.Fig.S16. Correlation structure of NMR media features day 4 (a) & 6 (b) with target responses**

*Phase 2: Multi-Omics Integrative Approach for Early Predictive Signatures*

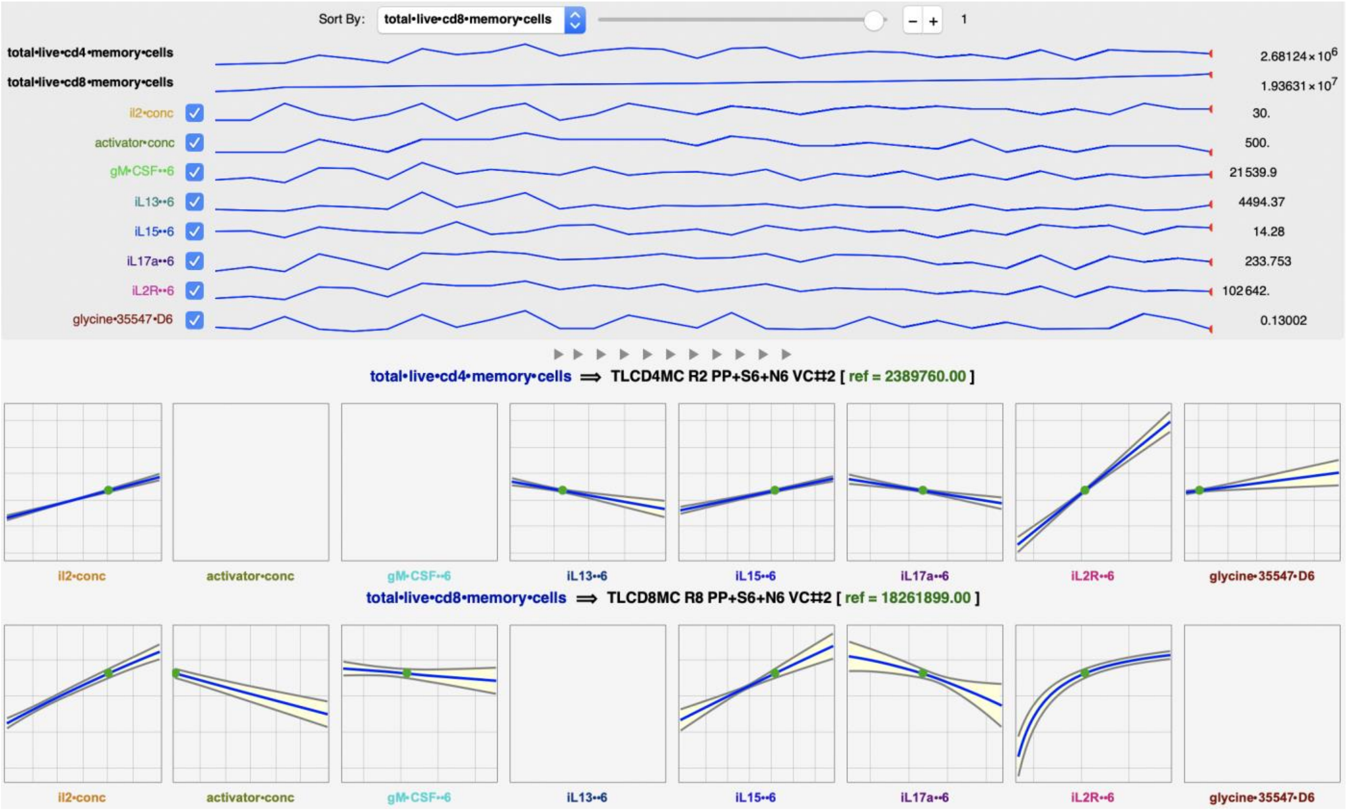


**Supp.Fig.S17. Multi-omics prediction profiles at day 6 using Symbolic Regression from DataModeler.** Maximum observed total live CD8$^+$ T$_N$+T$_{CM}$ cells and predicted values for Total live, CD4$^+$ and CD8$^+$.

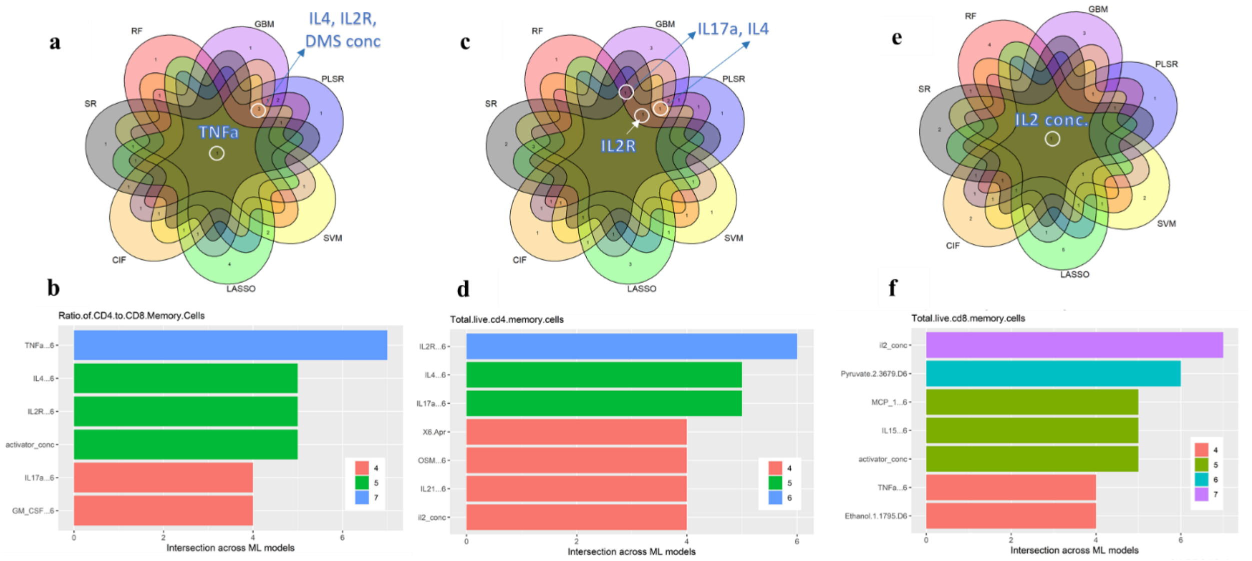


**Supp.Fig.S18. Overall feature consensus analysis of top-performing features in multi-omics models at day 6** for **a-b**) ratio of total live CD4$^+$ to CD8$^+$ T$_N$+T$_{CM}$ cells, **c-d**) total live CD4$^+$ T$_N$+T$_{CM}$ cells, and **e-f**) total live CD8$^+$ T$_N$+T$_{CM}$ cells

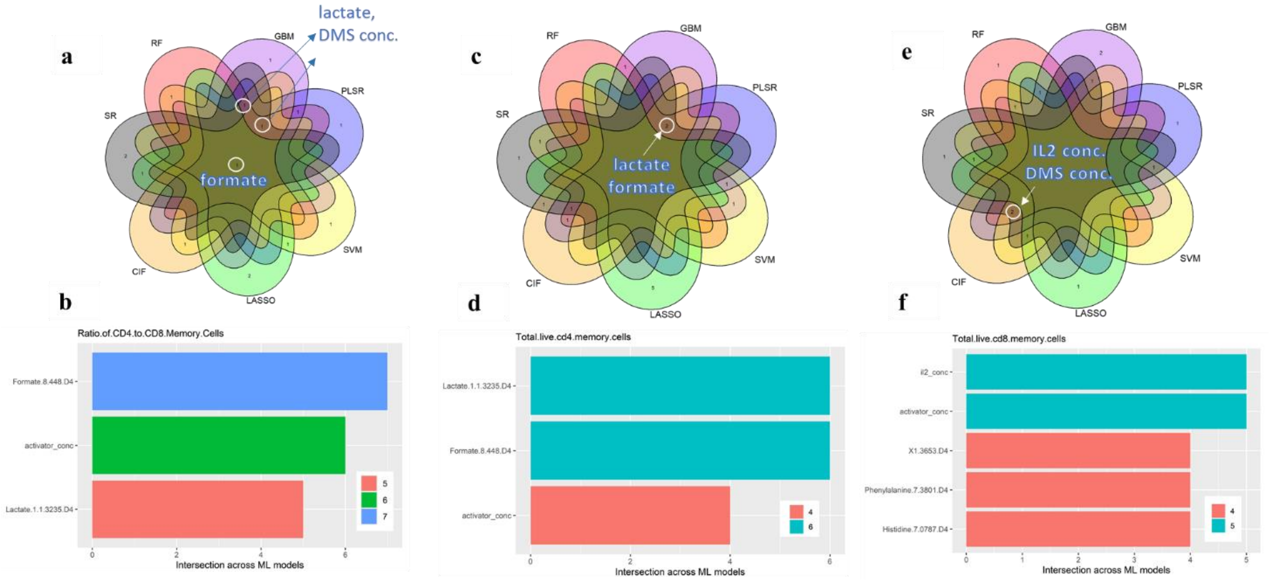


**Supp.Fig.S19. Overall feature consensus analysis of top-performing features in single-omics (NMR) models at day 4** for **a-b**) ratio of total live CD4$^{+}$ to CD8$^{+}$ $T_N$+$T_{CM}$ cells, **c-d**) total live CD4$^{+}$ $T_N$+$T_{CM}$ cells, and **e-f**) total live CD8$^{+}$ $T_N$+$T_{CM}$ cells

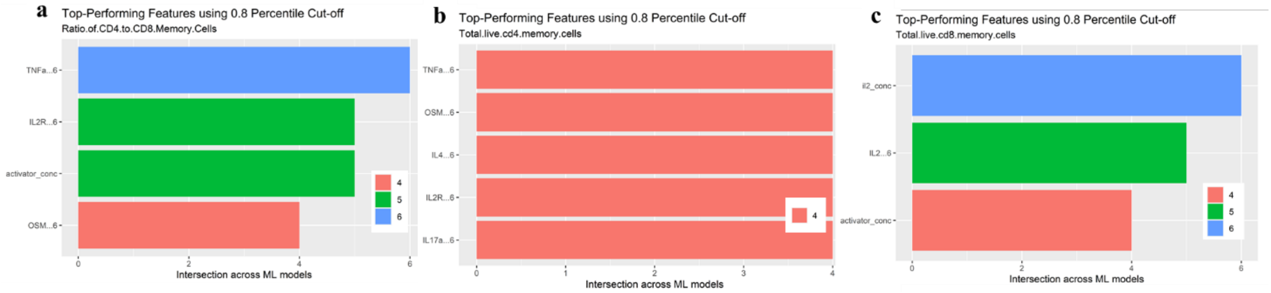


**Supp.Fig.S20. Overall feature consensus analysis of top-performing features in single-omics (Cytokine-S6) models at day 6** for **a**) ratio of total live CD4$^{+}$ to CD8$^{+}$ $T_N$+$T_{CM}$ cells, **b**) total live CD4$^{+}$ $T_N$+$T_{CM}$ cells, and **b**) total live CD8$^{+}$ $T_N$+$T_{CM}$ cells

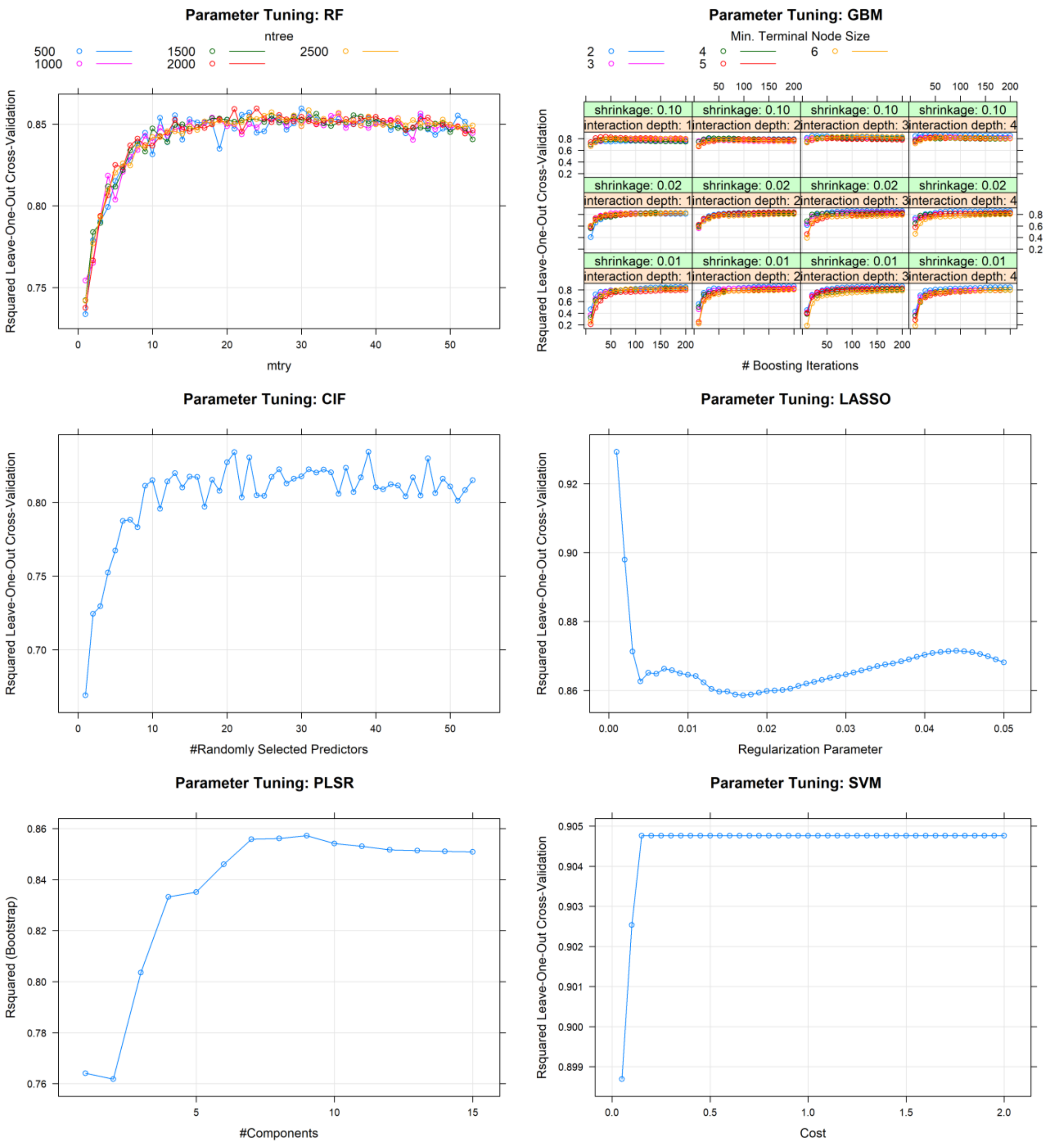


**Supp.Fig.S21. Parameter tuning for Ratio of $CD4^+$ to $CD8^+$ $T_N$+$T_{CM}$ Cells across all ML models from multi-omics integration at day 6.**

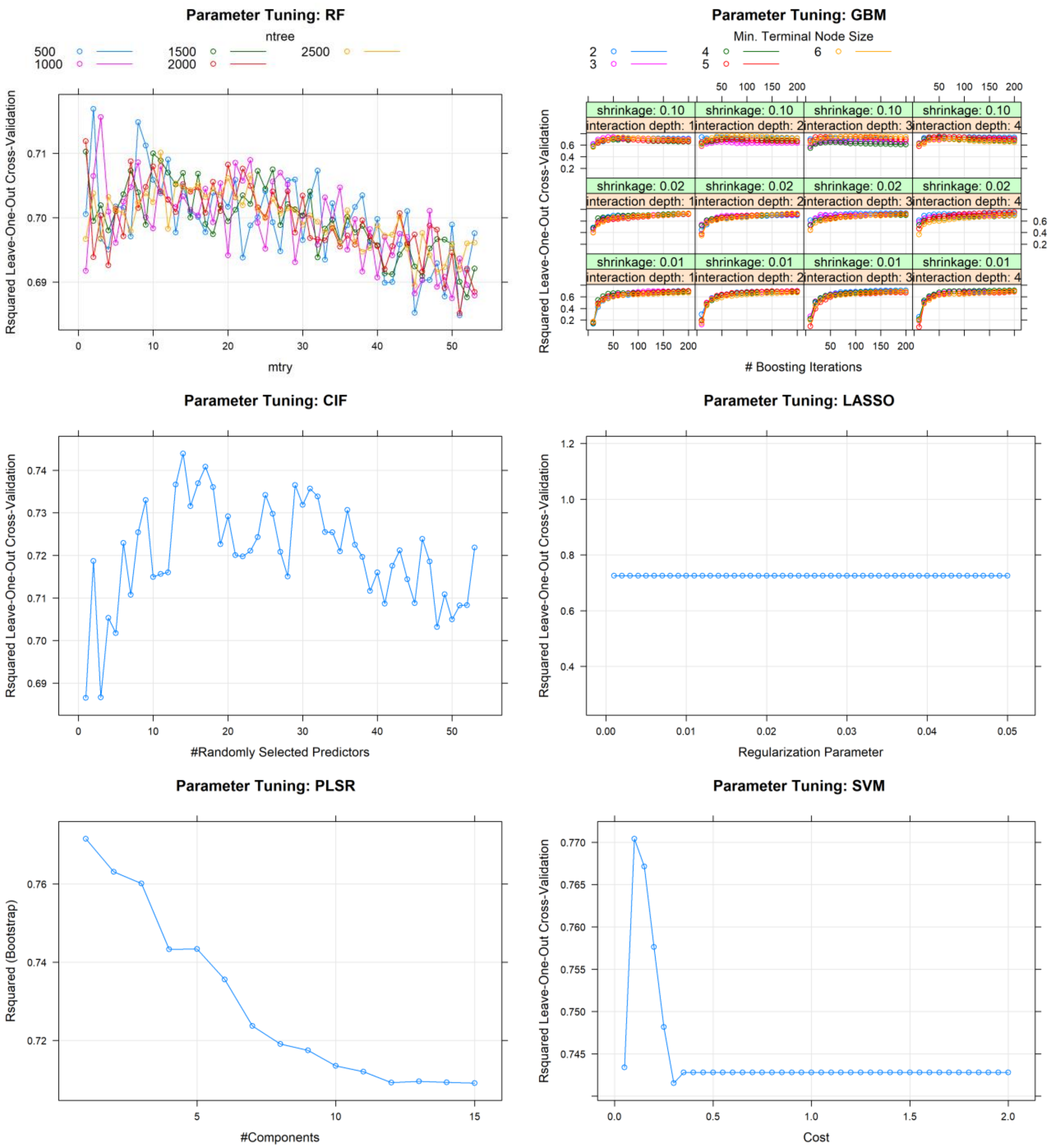


**Supp.Fig.S22. Parameter tuning for Total live $CD4^+$ $T_N$+$T_{CM}$ cells across all ML models from multi-omics integration at day 6.**

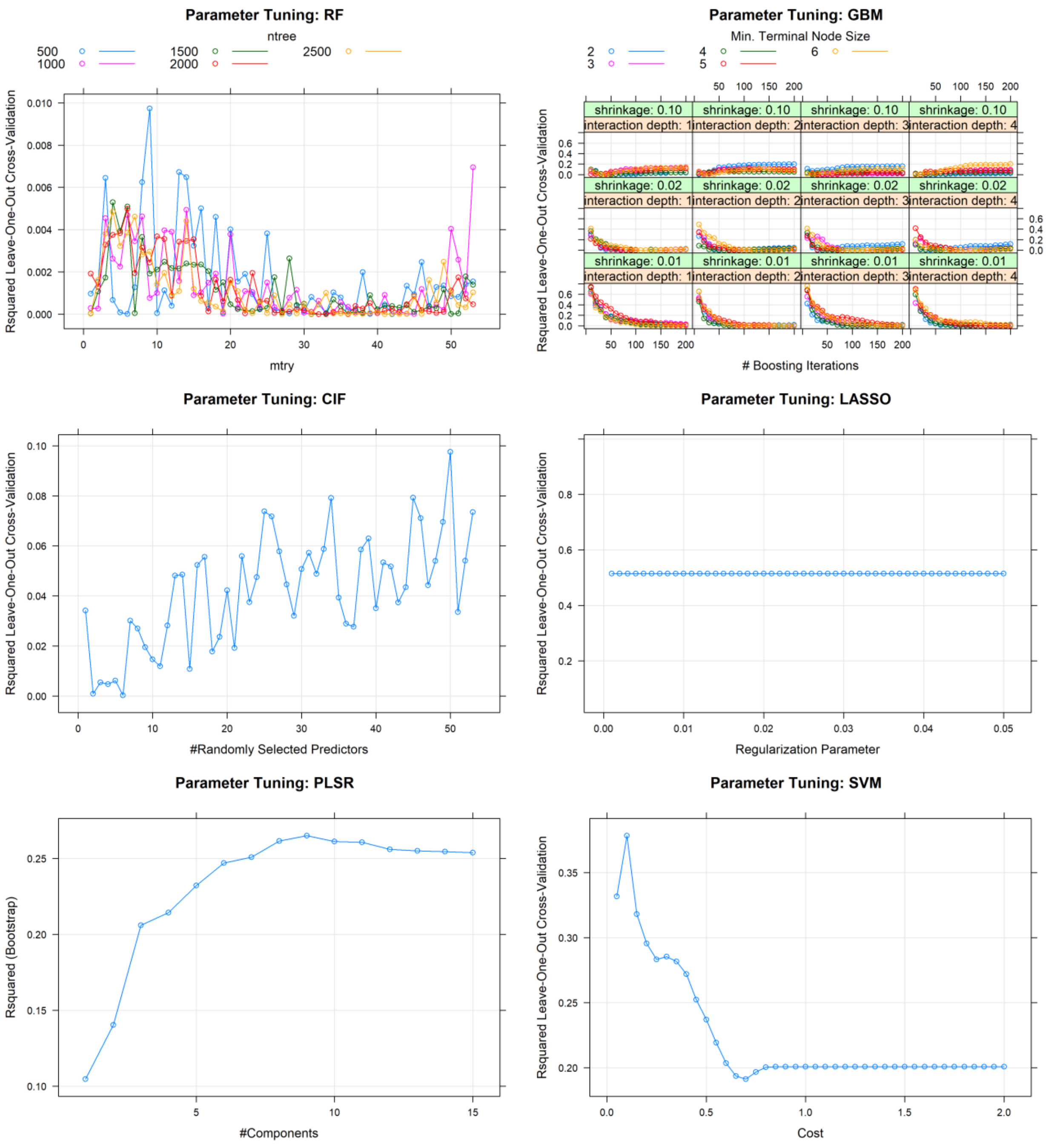


**Supp.Fig.S23. Parameter tuning for Total live CD8$^+$ T$_N$+T$_{CM}$ cells across all ML models from multi-omics integration at day 6.**

*Phase 2: Early Predictive Signatures using NMR Day 4 Modeling*

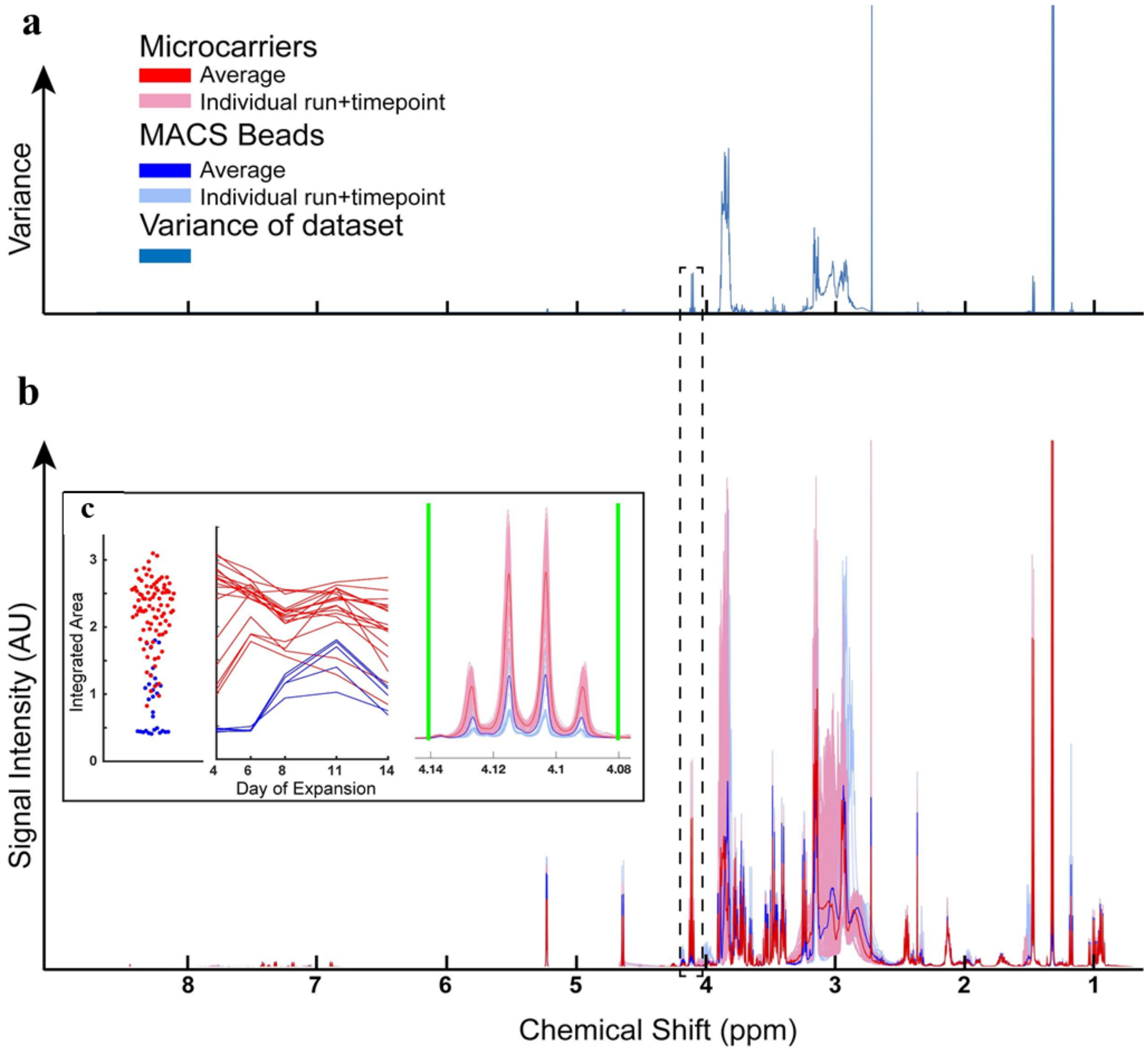


**Supp.Fig.S24. Variance based feature selection of NMR features for computational modeling.** a) Plot of variance across $^{1}H$ NMR spectrum for all experimental samples. X-axis matched to below. b) Plot of all experimental spectra for both microcarrier and MACS bead process runs. Averages shown in bold lines. c) Integration of feature highlighted in dashed box. Far right plot shows overlay of all experimental spectra and averages for both groups. Vertical green lines correspond to boundaries for feature integration. Center plot shows the trajectory of integrated values for individual runs (represented as continuous lines) over the expansion period indicated on X-axis. Far left-plot shows a distribution of integrated values for all samples over all timepoints.

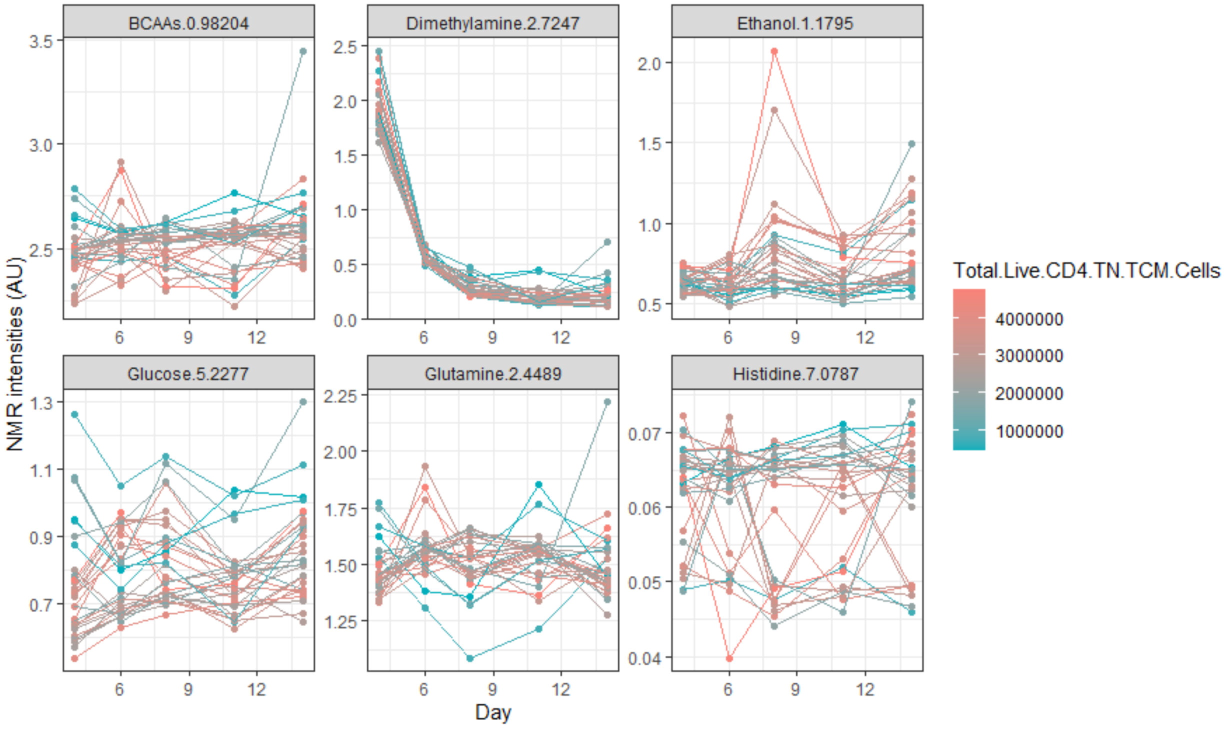


**Supp.Fig.S25. Media NMR intensities across monitoring times for total live CD4$^{+}$ T$_{N}$+T$_{CM}$ cells.**

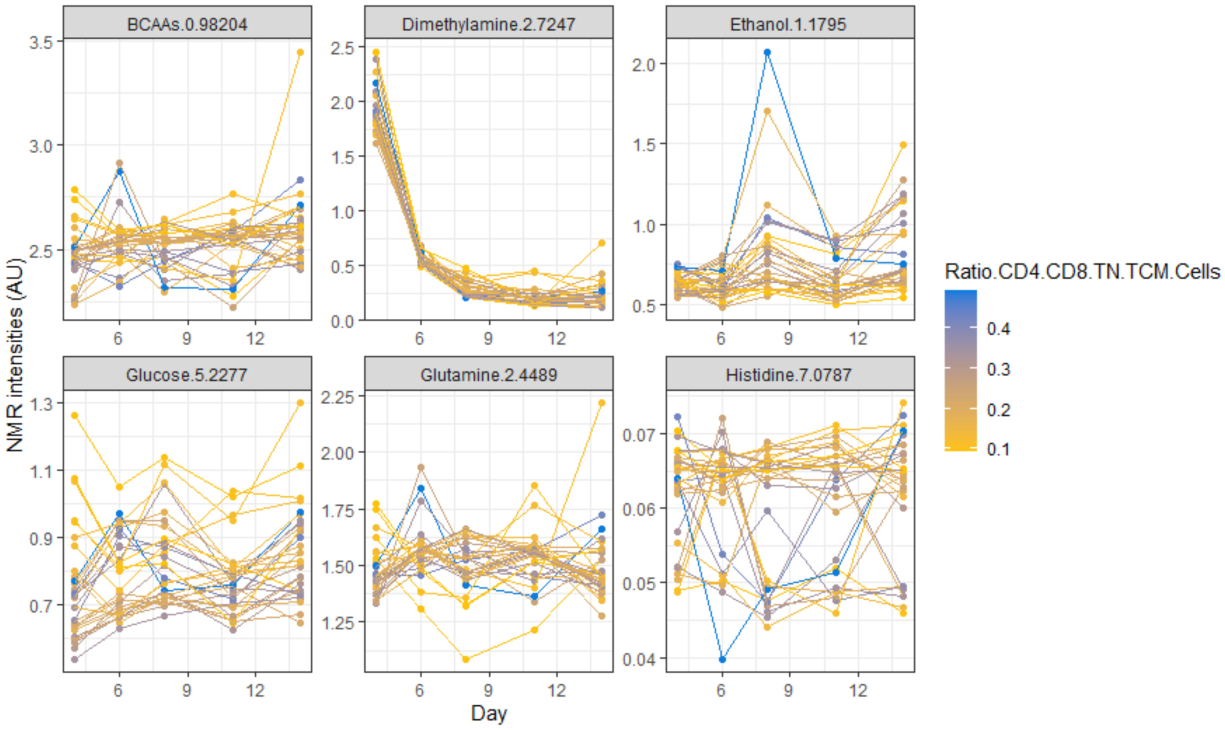


**Supp.Fig.S26. Media NMR intensities across monitoring times for ratio CD4$^{+}$/CD8$^{+}$ T$_{N}$+T$_{CM}$.**

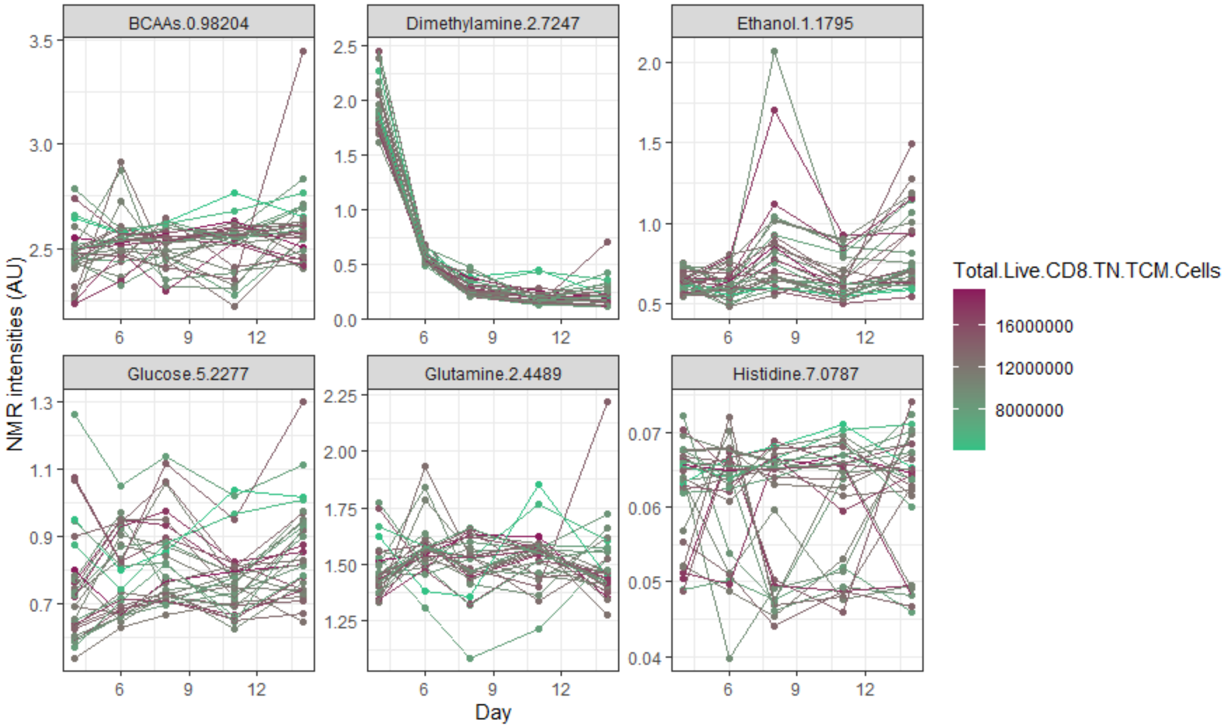


**Supp.Fig.S27: Media NMR intensities across monitoring times for total live CD8$^+$ $T_N$+$T_{CM}$ cells.**

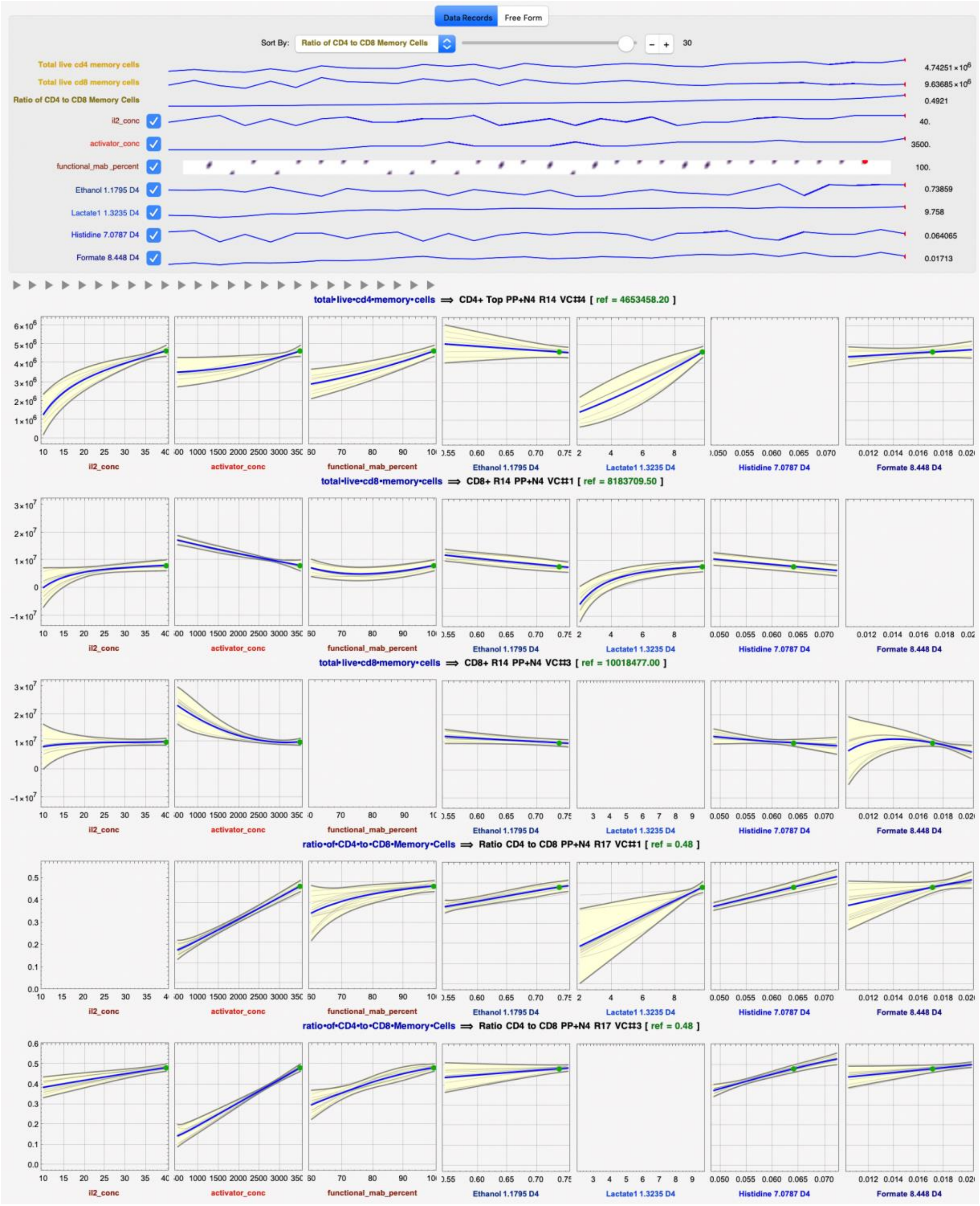


**Supp.Fig.S28: Predicted response profiles at the predicted optimum DMS process conditions for Total live CD4$^{+}$ T$_{N}$+T$_{CM}$ cells for NMR Models at Day 4 using DataModeler - View 1.** At this setting, the predicted value of a) Total live CD4+ T$_{N}$+T$_{CM}$ cells is 4.7 x $10^{6}$ vs observed 4.7 x $10^{6}$ ; Total live CD8+ naïve-memory yield is b) 0.8 x $10^{7}$ and c) 1.0 x $10^{7}$ vs observed 1.0 x $10^{7}$; Ratio of CD4+ to CD8+ T$_{N}$+T$_{CM}$ cells is d) 0.48 and e) 0.48 vs the observed 0.49.

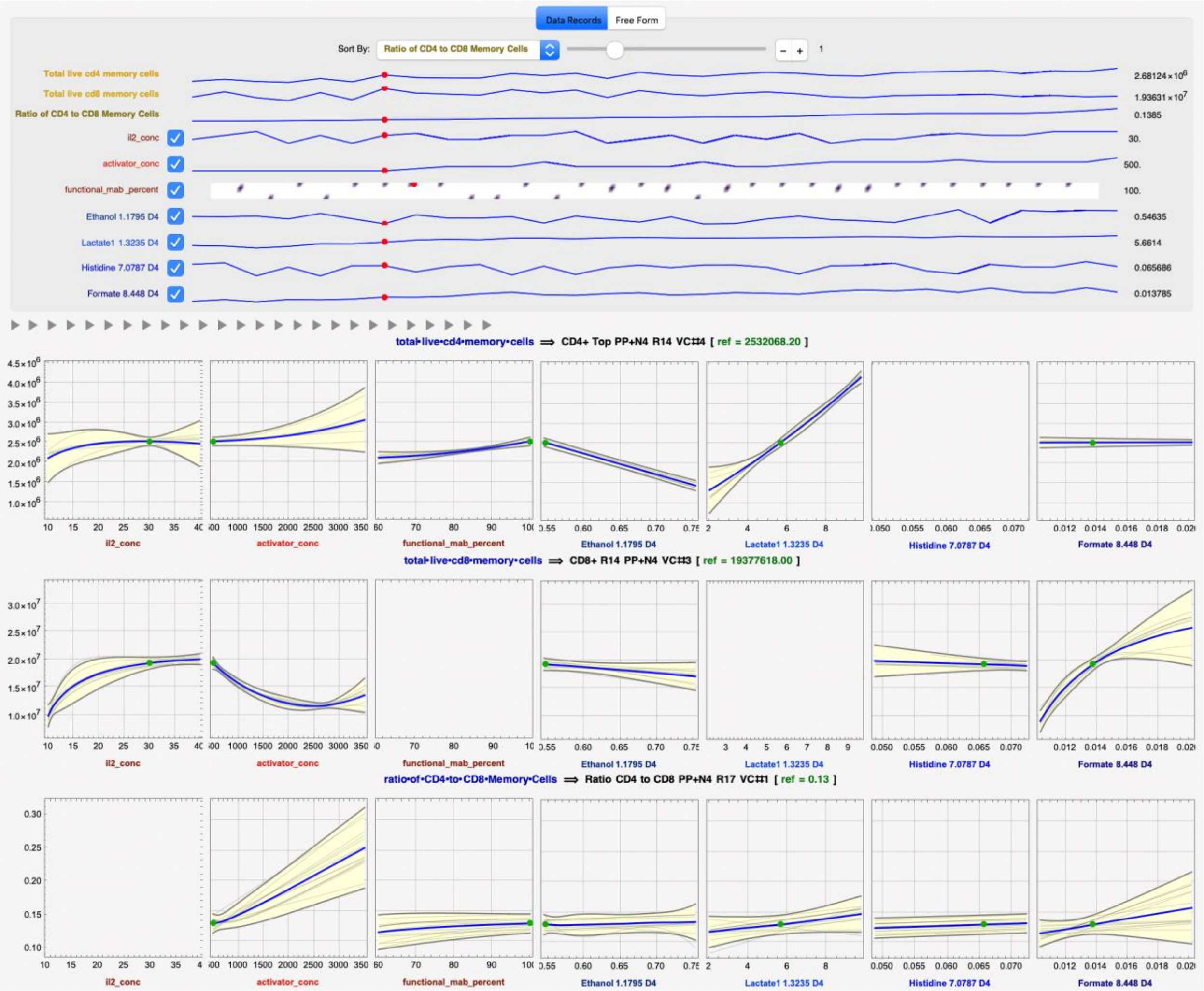


**Supp.Fig.S29: Predicted response profiles for NMR Models at Day 4 using DataModeler - View 2.** At this setting, the predicted value of a) Total live CD4+ $T_N$+$T_{CM}$ cells is 2.5 x $10^6$ vs observed 2.7 x $10^6$ ; the predicted value of b) Total live CD8+ naïve-memory yield is 1.9 x $10^7$ vs observed 1.9 x $10^7$ , and the predicted value of the c) Ratio of CD4+ to CD8+ $T_N$+$T_{CM}$ cells is 0.13 vs observed 0.14.

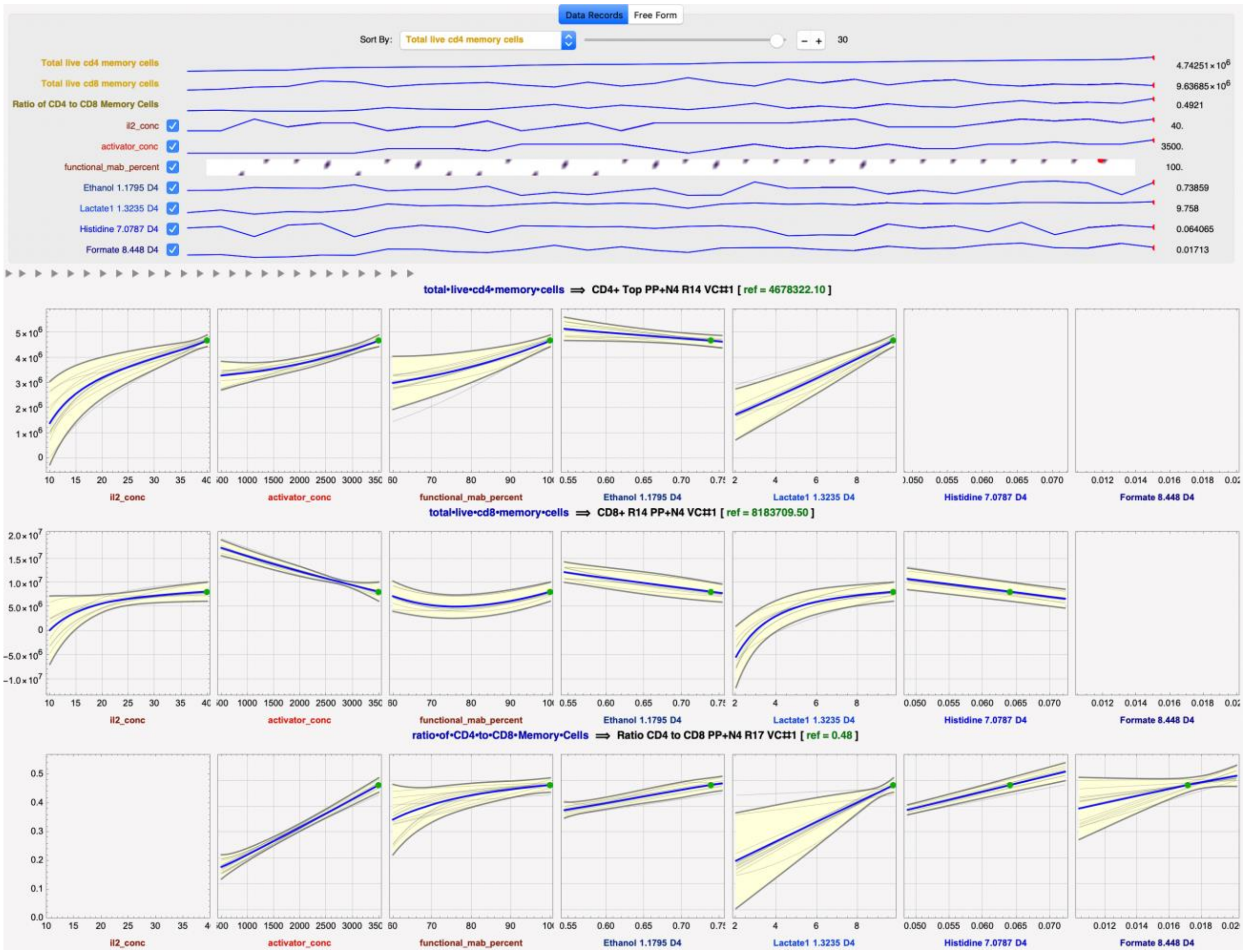


**Supp.Fig.S30: Predicted response profiles for NMR Models at Day 4 using DataModeler - View 3.** At this setting, the predicted value of a) Total live CD4+ $T_N$+$T_{CM}$ cells is 4.7 x $10^6$ vs observed 4.7 x $10^6$ ; the predicted value of b) Total live CD8+ naïve-memory yield is 1.0 x $10^7$ vs observed 0.8 x $10^7$ , and the predicted value of the c) Ratio of CD4+ to CD8+ $T_N$+$T_{CM}$ cells is 0.48 vs observed 0.48.

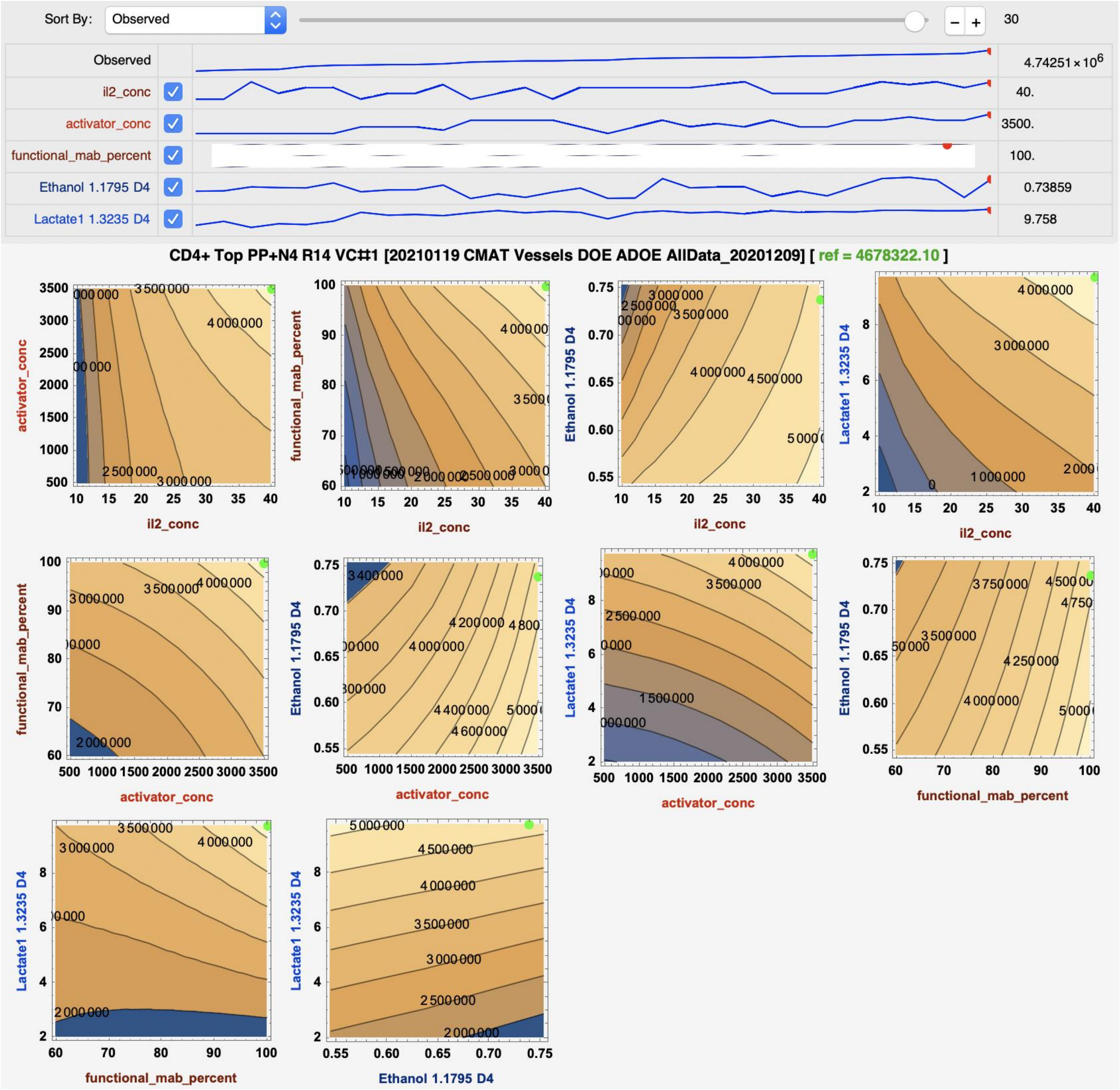


**Supp.Fig.S31: Response contour plots for NMR media analysis at day 4 using DataModeler – View 1**

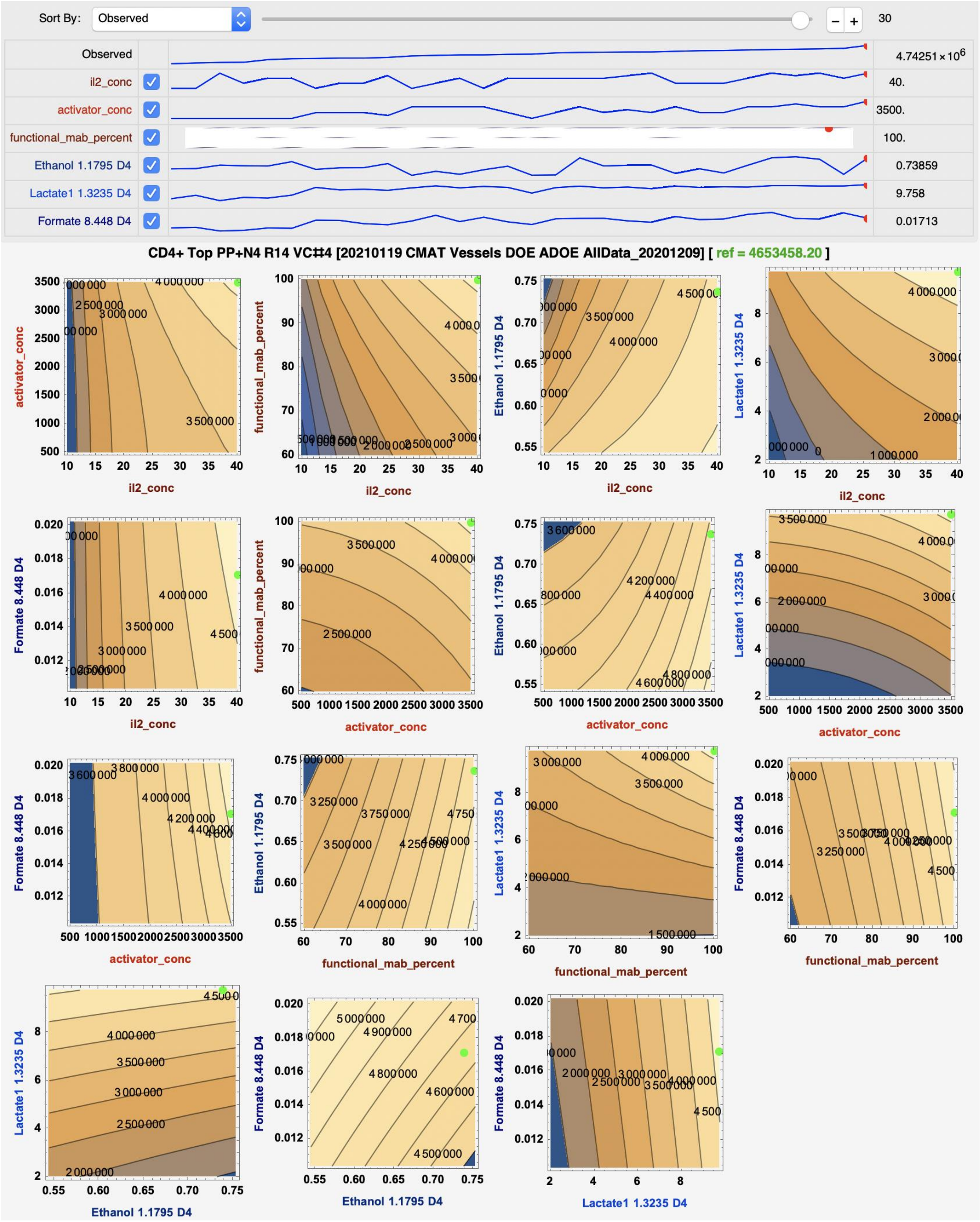


**Supp.Fig.S32: Response contour plots for NMR media analysis at day 4 using DataModeler – View 2**

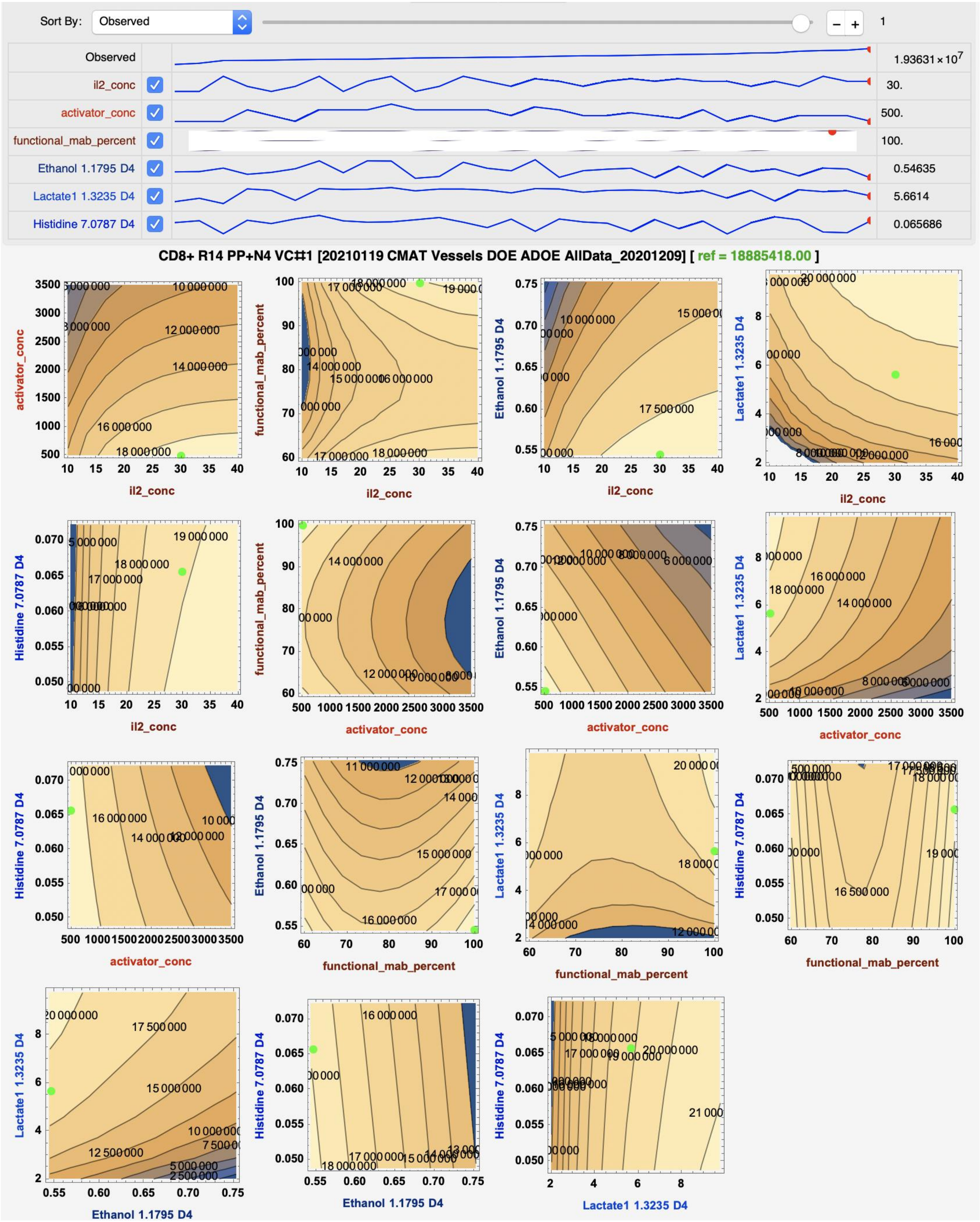


**Supp.Fig.S33: Response contour plots for NMR media analysis at day 4 using DataModeler – View 3**

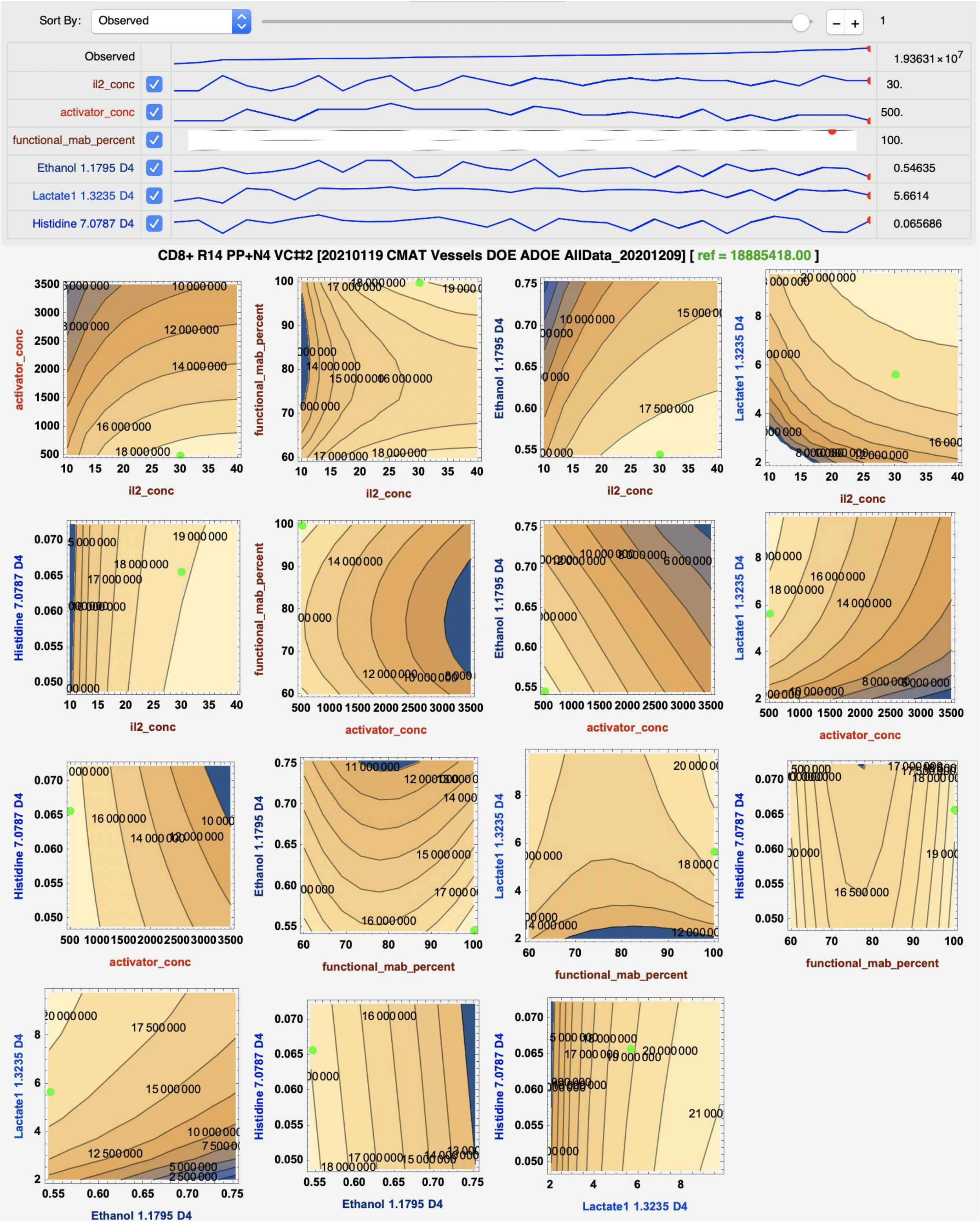


**Supp.Fig.S34: Response contour plots for NMR media analysis at day 4 using DataModeler – View 4**

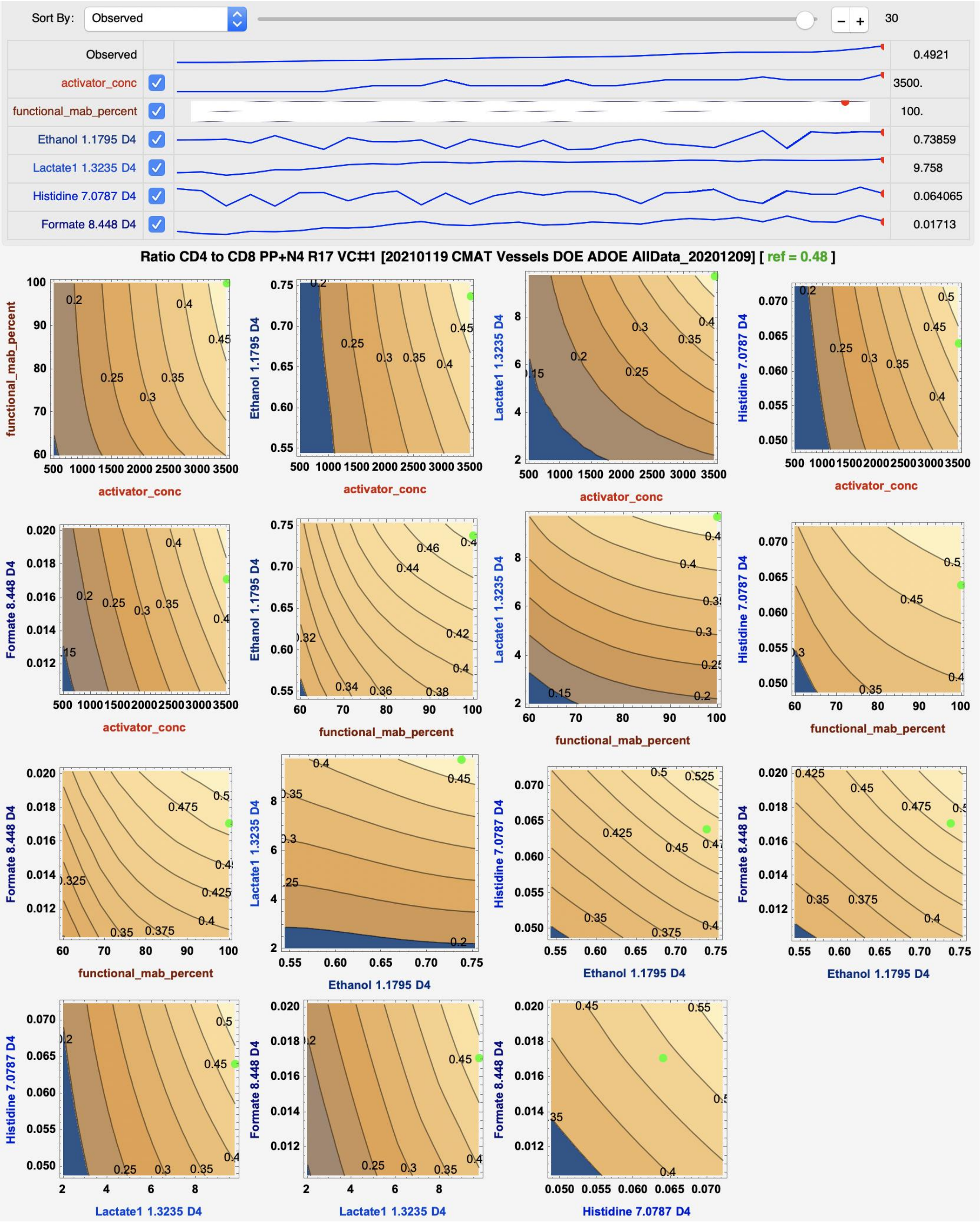


**Supp.Fig.S35: Response contour plots for NMR media analysis at day 4 using DataModeler – View 5**

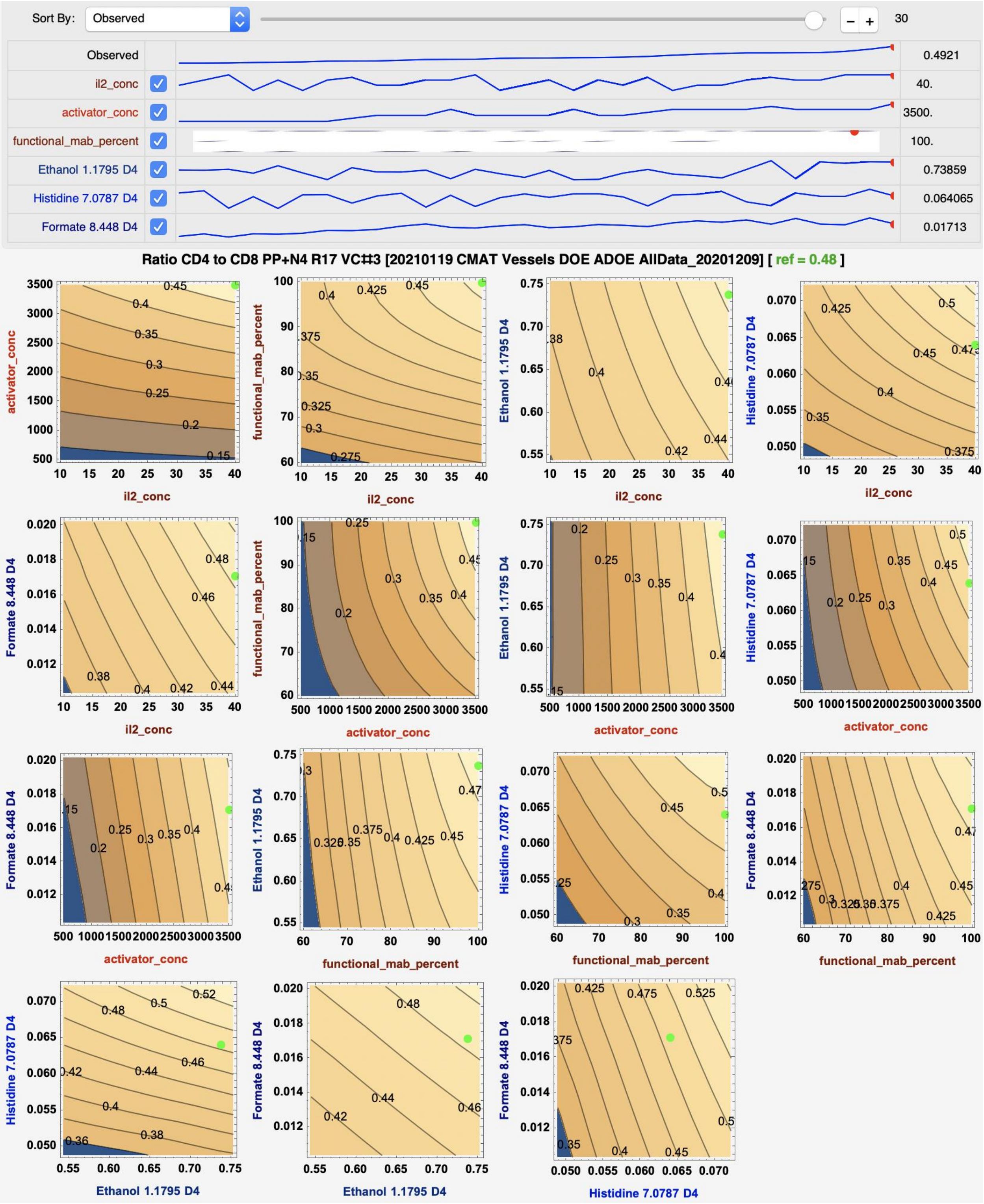


**Supp.Fig.S36: Response contour plots for NMR media analysis at day 4 using DataModeler – View 6**

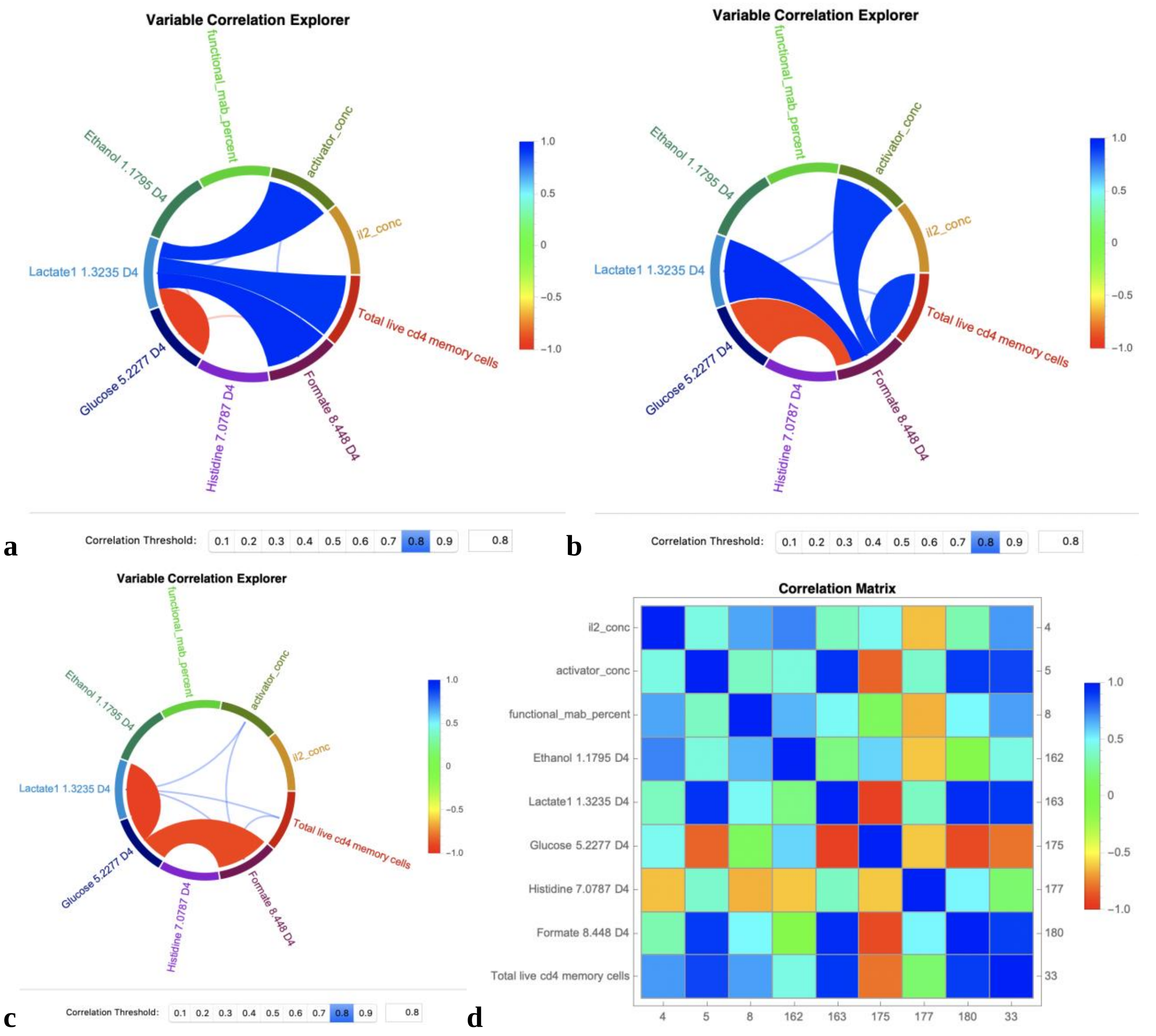


**Supp.Fig.S38: NMR Feature Correlation for SR-DataModeler models for NMR media analysis at day 4**: **a** lactate is strongly positively correlated with formate, DMS Conc and Total Live $CD4^{+}$ and negatively correlated with glucose; **b** formate is strongly positively correlated with lactate, DMS Conc and Total Live $CD4^{+}$ and negatively correlated with glucose; **c** glucose is negatively correlated with lactate and formate; **d** NMR correlation Matrix.

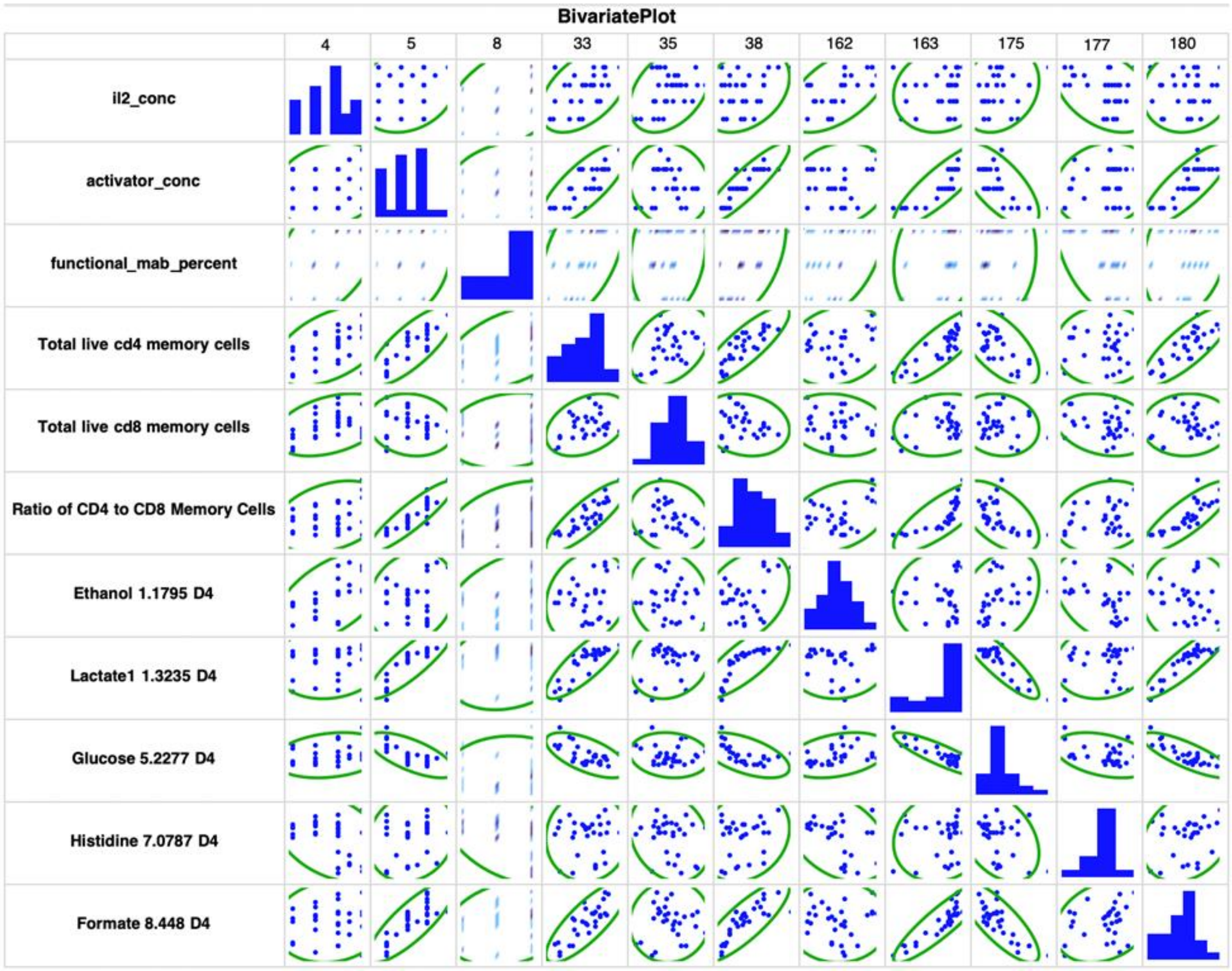


**Supp.Fig.S39: Bivarate Plot of NMR features predictive of CD4$^{+}$ $T_{N}$+$T_{CM}$ cells, CD8$^{+}$ $T_{N}$+$T_{CM}$ cells, and ratio CD4$^{+}$ $T_{N}$+$T_{CM}$ to CD8$^{+}$ $T_{N}$+$T_{CM}$ cells.**
